# Multidimensional variation and population stratification across 8000 complete human centromeres

**DOI:** 10.64898/2026.07.22.740206

**Authors:** Yanqing Sun, Shijie Wan, Lei Nie, Dan Yu, Feifei Zhou, Yiqing Yang, Xiangyu Yang, Anguo Liu, Quanyu Chen, Kezhi Fu, Qingyang Ni, Yaoxi He, Bing Su, Yafei Mao, Kai Ye, Xiaofei Yang, Guojie Zhang, Dongya Wu

## Abstract

Human centromeres are indispensable for the faithful segregation of chromosomes during cell division, yet their highly repetitive nature has historically precluded comprehensive characterization, leaving fundamental questions about their sequence diversity, evolution trajectories and function dynamics unresolved. Here, we generated 6,312 complete human centromere sequences from 320 phased genome assemblies in Asian Pan-Genome project phase 1. By integrating the assemblies from the Human Pangenome Reference Consortium (HPRC) and Human Genome Structural Variation Consortium (HGSVC), we constructed a multidimensional genetic variation map encompassing over 8,000 gapless centromeres. Centromeric satellite arrays account for 4.19% to 6.01% of the whole genome, with substantial variations in size and architecture across chromosomes. Using a refined alpha satellite clustering approach that captures global diversity, we identified 195 higher-order repeat (HOR) arrays, 56.4% of which are absent from the T2T-CHM13 reference genome. Extensive structural variations across multiple dimensions exhibit population stratification, including centromeric haplotypes (CenHaps), ultra-large pericentric inversions spanning up to 36.6 Mbp, and inter-chromosomal HOR sharing that reflects sequence exchange among chromosomes. Integrating CENP-A CUT&Tag experiments and long-read-based DNA methylation profiles, we demonstrate that 16.8% of centromeres harbor multiple potential kinetochore assembly sites, and CenHap-specific local HOR homogenization is associated with kinetochore positioning. Despite global suppression of recombination at centromeres, we observed asymmetric linkage disequilibrium flanking centromeres and an ancient recombination event within the centromere of chromosome 19. Furthermore, contrary to the prevailing assumption of high mutation rates in centromeres, our estimates based on stringent orthology reveal no significantly higher single-base substitution rates for centromeres relative to flanking pericentromeric regions with substantial variations across chromosomes, despite extraordinary structural plasticity. Collectively, these multi-scale centromeric variations provide a global view of human centromere diversity and population stratification, fundamentally redefine centromere evolution through a dual-track model balancing structural innovation with mutational constraint, and establish an essential resource for investigating centromere biology and a baseline reference for diagnosing centromere-associated disorders.

## Introduction

Centromeres are essential chromosomal loci that ensure the faithful segregation of chromosomes during cell division by serving as assembly sites for the kinetochore complex^1^. Defects in their establishment, inheritance, or maintenance compromise genome stability, and are implicated in developmental disorders and cancers^1–3^. Despite their conserved function across eukaryotes, human centromeres have remained enigmatic due to their extreme repetitiveness, dominated by tandemly arranged alpha satellite (αSat) DNA organized into higher-order repeat (HOR) arrays^4–6^. These highly homogeneous sequences have historically evaded comprehensive characterization, leaving fundamental questions regarding centromeric diversity, evolution, and functional dynamics unresolved^7,8^.

Recent advances in long-read sequencing technologies have enabled the generation of the first complete human genome assembly (T2T-CHM13), providing an unprecedented view of human peri/centromeric sequences^9,10^. Follow-up efforts, including T2T-CN1^11^, T2T-CHM1^12^, T2T-CQ^13^, Q100-HG002^14,15^, RPE1v1.1^16^and initiatives from HPRC and HGSVC^17^, have begun to unravel centromeric diversity across individuals. However, several critical limitations persist. Current assemblies exhibit disappointingly low centromere completion rates, leaving substantial peri/centromeric diversity uncharacterized. Existing human αSat/HOR annotation approaches using hidden Markov models with limited sequence diversity have failed to capture population-specific and individual-specific variants that are crucial for understanding human evolution and disease susceptibility^9,18,19^. Additionally, the inability to systematically compare centromeric architectures across populations has obscured their evolutionary dynamics, including the longstanding debate over whether centromeres evolve under relaxed or constrained mutation rates. Finally, the relationship between centromeric structural variation and functional consequences remains poorly defined, despite well-documented links to infertility and chromosomal disorders^2,20–23^.

Here, we generate 6,312 complete human centromeres from 320 phased genome assemblies of East Asian individuals in Asian Pan-Genome project phase 1^24^ (APGp1), and present a multidimensional variation map encompassing over 8,000 gapless centromere sequences, by integrating previous assemblies from HPRC year 1 (HPRCy1) and HGSVC phase 3 (HGSVC3). Using a newly developed reference-free annotation pipeline, we systematically characterize multi-scale structural diversity and define population-stratified centromeric haplotypes that may modulate kinetochore positioning. Our analyses uncover inter-chromosomal HOR sharing, document pericentric inversions spanning entire centromeres, and delineate recombination and single-base substitution rates in human centromeres. These findings establish critical clinical reference ranges for evaluating meiotic stability, and fundamentally reshape our understanding of human centromere evolution.

## Results

### A comprehensive atlas of human complete centromere sequences

Leveraging the combined power of high-depth PacBio high-fidelity (HiFi; ∼51×) and Oxford Nanopore Technology (ONT) ultra-long (>100 Kbp) sequencing reads (∼62×), we assembled 6,312 gapless centromere sequences from 320 phased East Asian (EAS) genomes in APGp1, accounting for 85.8% of all 7,360 centromeres (**Supplementary Fig. 1a**; **Supplementary Table 1**). This achievement marks a dramatic leap forward from previous efforts, surpassing the centromere gaplessness rates of HPRCy1^25^ (20.6%) and HGSVC3^17^ (66.3%), and establishes the most comprehensive collection of complete human centromere sequences to date. With an average single-base quality value (QV) of 85.9, our centromere sequences meet the highest standards for genomic accuracy (**Supplementary Fig. 1b)**. We subjected each assembly to rigorous validation using comprehensive approaches, including Genome Continuity Inspector^26^, Nucfreq^27^, Flagger^25^, GAVISUNK^28^, and VerityMap^29^ (**Supplementary Note 1**; **Supplementary Figs. 1c-1e**; **Supplementary Table 1**). Collectively, an average of 19.7 centromeres per haplotype assembly were confirmed as completely assembled, with chromosomal representation averaging 273 centromeres per euchromosome and 153 per sex chromosome (**Supplementary Table 1**). These high-quality assemblies provide an unprecedented foundation for dissecting the architectural complexity and evolutionary dynamics of human centromeres at the population scale.

Canonical human centromeric satellite arrays, including 171-bp αSat^4,5^, classical human satellites^30,31^ (HSat1, Hsat2 and Hsat3), beta satellites^32^ (βSat) and gamma satellites^33^ (γSat), collectively accounted for 4.19%-6.01% of each APGp1 whole-genome haplotype assembly, yet their size distributions exhibited striking chromosome-specific patterns (**Fig. 1a**; **Supplementary Fig. 2**; **Supplementary Tables 2 and 3**). Specifically, αSat arrays, the core of functional centromeres, averaged 3.66 ± 1.40 Mbp (mean ± s.d., standard deviation) per centromere, with chromosome 19 (5.67 ± 0.94 Mbp) and chromosome 1 (5.49 ± 0.88 Mbp) having the largest arrays and chromosome Y the smallest (0.75 ± 0.30 Mbp). The three largest satellite arrays in the human genome include two HSat2 arrays on chromosome 1 (3.16-23.91 Mbp) and chromosome 16 (2.09-19.82 Mbp), along with one HSat3 array on chromosome 9 (0.76–20.69 Mbp), all of which exhibit substantial size variability across individuals. HSat1, the most AT-rich satellite, was predominantly localized to chromosomes 3 (1.76 ± 0.96 Mbp), 4 (1.41 ± 0.56 Mbp), and 13 (1.65 ± 0.95 Mbp), with additional individual-specific expansions on acrocentric chromosomes 14, 15, 21 and 22 (4.57-5.39 Mbp). The two additional satellite families, βSat (3.31 ± 1.09 Mbp) and γSat (0.57 ± 0.54 Mbp), generally displayed conserved array sizes, though βSat arrays on chromosome 1 showed a distinctive bimodal distribution (**Supplementary Fig. 2**), indicative of two divergent clusters within the EAS population. Similar bimodal distributions were observed for βSat arrays on chromosome 13, 14, 15, and 21, though for these acrocentric centromeres, the apparent bimodality may reflect residual assembly incompleteness near the telomeric ends rather than true biological divergence (**Supplementary Fig. 2**).

**Figure 1.**
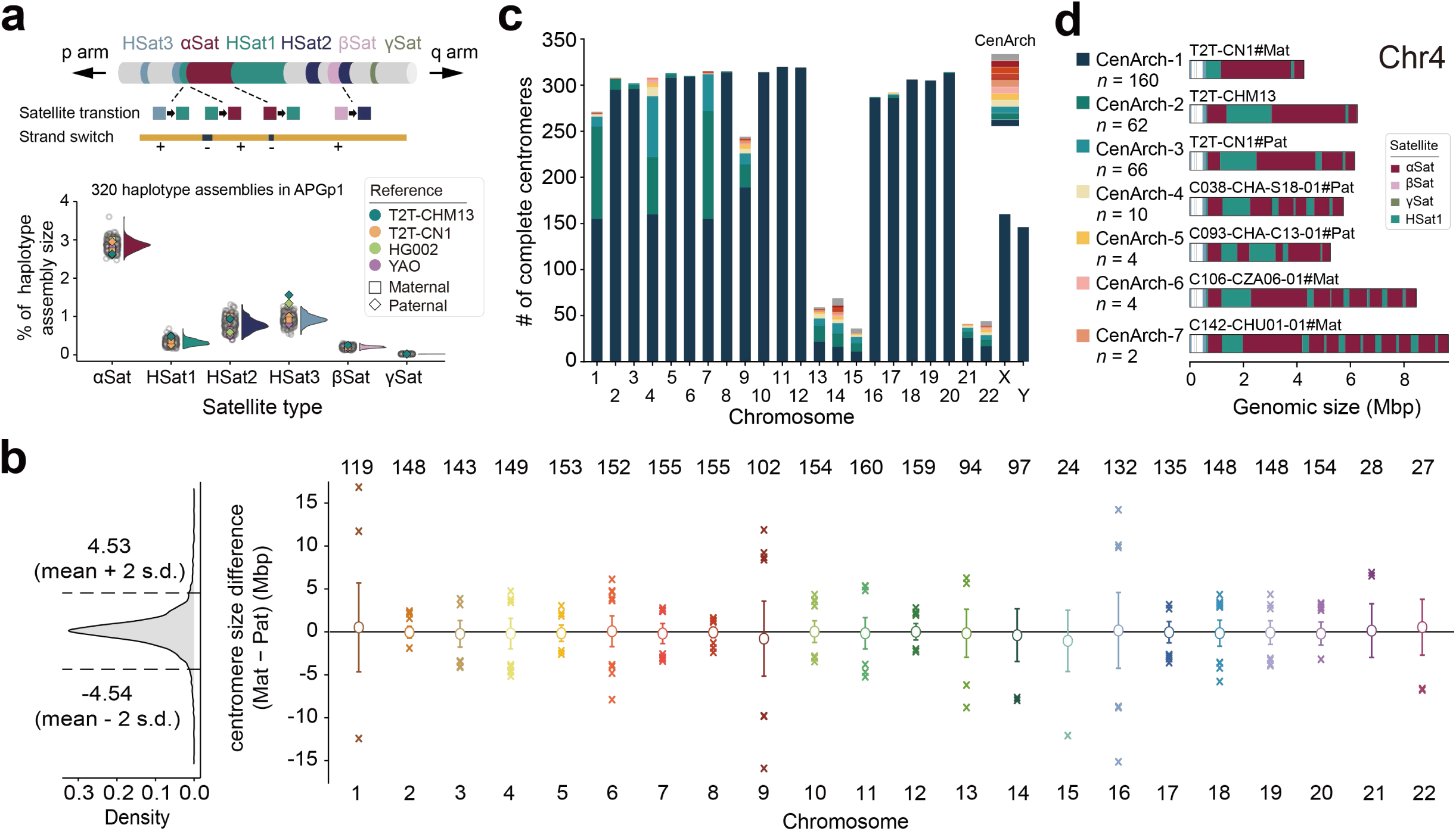
Size and structure variations of centromeric satellite arrays across the APGp1 assemblies. **a**, Schematic diagram of human centromere structure (upper) and the proportional composition of distinct centromeric satellite types across APGp1 haplotype assemblies. Previously published T2T-level assemblies (T2T-CHM13, T2T-CN1, YAO and Q100-HG002) are highlighted with squares (maternal) and diamonds (paternal). Open grey circles denote the satellite array proportions for each APGp1 haplotype assembly. **b**, Density distribution of centromere size differences between matched maternal (Mat) and paternal (Pat) centromere pairs with fully gapless assemblies. Dashed lines demarcate the mean ± 2 standard deviations (s.d.) interval. For each chromosome, circles denote mean values, and error bars indicate ± 1 s.d. Haplotype pairs falling outside the mean ± 2 s.d. intervals are annotated as outliers (cross symbols). The number of complete maternal-paternal centromere pairs per chromosome is labeled at the top. **c**, Distribution of centromere architectures (CenArchs) across chromosomes in APGp1 cohort. Colours and architecture indices denote the chromosome-specific centromere architectures, ordered by decreasing haplotype frequency.For acrocentric chromosomes (13, 14, 15, 21 and 22), beyond the gapless criterion, we additionally removed any centromere lacking annotated telomere or rDNA to avoid misclassification due to incomplete assembly of peri-telomeric satellite sequences. **d**, Representative CenArchs derived from 302 complete centromeres on chromosome 4 in APGp1. The number of haplotype assemblies assigned to each CenArch is indicated on the left. Reference assemblies are used as representative structures where applicable. CenArchs are numbered in descending order of global frequency.

Centromere size comparison between EAS assemblies from APGp1 and previous studies (HPRCy1 and HGSVC3) revealed no obvious batch effects, enabling us to investigate the population differences in centromeric satellite array sizes (**Supplementary Fig. 3**). To control for potential bias arising from unequal superpopulation sample sizes, we performed a balanced random downsampling strategy (see **Methods**). Comparison between EAS and Africans (AFR) superpopulations revealed significantly different patterns across 12 chromosomes (*p* < 0.01, Wilcoxon rank-sum test; **Supplementary Figs. 4 and 5; Supplementary Table 4**; **Methods**). Specifically, CENX αSat arrays are significantly smaller in AFR compared to EAS (mean *p* = 3.9×10^-9^). Significant size reductions were also observed when compared to other superpopulations, including European (EUR, mean *p* = 0.008), South Asian (SAS; mean *p* = 0.001), and American (AMR, mean *p* =0.0002; **Supplementary Fig. 4**). Conversely, HSat3 arrays on chromosome 9 are significantly smaller in EAS than those in AFR (mean *p* =0.0052), directly contributing to reduced overall CEN09 sizes in EAS (**Supplementary Figs. 5 and 6**).

Given that large size discrepancies between homologous centromeres may induce meiotic abnormalities and contribute to conditions such as trisomy^21^, we investigated maternal-paternal centromere size differences across 160 healthy EAS individuals. The vast majority (94.3%; 2,581 of 2,736) of homologous centromere pairs fell within the mean ± 2 s.d. range (-4.54 - 4.53 Mbp, **Fig. 1b**), establishing a baseline for normal variations. αSat arrays show comparable distributions, with a mean ± 2 s.d. range of ∼2.37 Mbp and rare outliers displaying variations up to ∼6 Mbp (**Supplementary Fig. 7**). Notably, we identified nine parental centromere pairs with size differences exceeding 10 Mbp that were also classified as statistical outliers, warranting further clinical follow-up to assess potential health risks to the individuals involved (**Fig. 1d**). Decomposing the parental size differences for these outlier pairs by satellite types revealed that eight of the nine per-chromosome outliers were concentrated on chromosomes 1, 9, and 16, driven primarily by HSat array expansion and contractions (**Supplementary Table 5**). Such patterns were also commonly observed in other superpopulation (**Supplementary Table 5**). We detected no significant systematic maternal-or paternal-allele length bias across any comparison (all BH-adjusted *p* > 0.05), providing no evidence for directional asymmetry (**Supplementary Fig. 7**). These results establish critical reference ranges that may inform risk assessment for meiotic abnormalities based on homologous centromere size differences.

Beyond satellite DNA, centromeres harbor a complex landscape of transposable elements (TEs) and genes that contribute to their functional diversity^9,34^. An average of 32.8 ± 2.2 Mbp of TEs (15.2 ± 1.0% of centromeric regions) were annotated per haplotype, with LINE/L1 being the most abundant by size (14.3 ± 0.9 Mbp, 6.6% of centromeres) and SINE/Alu showing the highest insertion frequency (133.7 ± 8.8 counts per Mbp; **Supplementary Table 6**). TE insertion patterns show clear preference among satellite arrays (**Supplementary Fig. 8a**), where SINE/Alus are predominantly enriched in βSat (∼37 counts per Mbp) and HSat3 (∼81 counts per Mbp) arrays, while LTR/ERV1s are enriched in γSat arrays (∼20 counts per Mbp). While TE insertion numbers are broadly conserved across populations, variable patterns are observed, for example, LINE/L1 within CEN07 αSat arrays and CEN16 HSat2 arrays, and Alu/SVA within CEN04 HSat3 arrays (**Supplementary Fig. 8b**). We also identified ∼4,115 centromeric gene/pseudogene elements per haplotype, encompassing ∼204 candidate protein-coding genes. A core set of 39 genes is universally conserved across all assemblies, comprising 16 uncharacterized genes, 22 members of Zinc Finger (ZNF) family, and *DUX4* homologs, the latter being a key pathogenic driver of Facioscapulohumeral Muscular Dystrophy^35^. Intriguingly, *DUX4* homologs within the βSat regions of acrocentric chromosomes exhibited discrete, multi-layered copy numbers (e.g., ∼20 and ∼90 copies on chromosome 14; **Supplementary Table 7**). In contrast, *DUX4* homologs within βSat regions of other chromosomes (e.g., chromosomes 3, 5 and 9) remained stable at the single-copy level (**Supplementary Fig. 9a**). The distinction between discrete-layered and stable *DUX4* homologs is mirrored in their phylogeny: high-copy active arrays formed a complex, rapidly evolving clade, whereas low-copy centromeric paralogs and functional subtelomeric homologs formed compact, stable clades (**Supplementary Fig. 9b**).

### Arrangements of human satellite arrays

Human centromere sequences exhibit extraordinary structural plasticity, characterized by frequent rearrangements, including satellite inversions, insertions and deletions that generate remarkable architectural diversity across individuals and populations^9,11^. To systematically quantify these structural variations (SVs), we first analyzed sequence strand switches across all APGp1 centromeres (**Supplementary Fig. 10a**). βSat arrays, predominantly localized to chromosome 1 and five acrocentrics, along with HSat3 arrays in CEN09, notably exhibited hundreds of strand switches across assemblies (**Supplementary Fig. 10b**; **Supplementary Table 8**). The number of switch breakpoints significantly correlated with array lengths for both βSat (Pearson’s *r*= 0.90, *p* = 0) and HSat3 (*r* = 0.90, *p* = 2.1×10^-195^) arrays (**Supplementary Figs. 10c** and **10d**). Notably, these strand switches are randomly distributed across the entire array rather than concentrated at specific hotspots, suggesting they arise as stochastic byproducts of the repeat array expansion process (**Supplementary Fig. 10e**). Within αSat arrays, ∼89.6% of strand switches localized to monomeric regions, composed of divergent, non-repetitive αSat monomers. However, we also detected strand switches in active HOR arrays, which serve as the primary functional units of the centromere where kinetochore proteins preferentially associate, across nine chromosomes (**Supplementary Table 9**). Among these, we confirmed the presence of a polymorphic megabase-scale inversion (1.42 ± 0.32 Mbp) inside the active HOR array of CEN01^9,36^, present in 55 haplotype assemblies in APGp1 with an allele frequency (AF) of 20.3% (**Fig. 1c**; **Supplementary Figs. 11a and 11b**). Notably, one haplotype assembly (C112-CUG04-01#Pat) exhibited not only inversions but also a large subsequent duplication spanning both the inverted active HOR array and adjacent inactive HOR regions (**Supplementary Figs. 11c and 11d**).

We next characterized transitions between distinct satellite array types to understand the higher-order architecture of centromeres (**Fig. 1a**). Across the genome, fifteen chromosomes (2, 3, 5, 6, 8, 10, 11, 12, 16, 17, 18, 19, 20, X and Y) displayed a nearly fixed architecture, with over 90% of haplotypes sharing this structure (**Supplementary Figs. 12 and 13**). Notably, the centromeric architecture of CEN03 in T2T-CHM13 was shared by only two haplotype assemblies in our dataset, indicating that T2T-CHM13 does not represent human population majority for this chromosome (**Supplementary Fig. 13**). Other nine chromosomes (1, 4, 7, 9, 13, 14, 15, 21 and 22) exhibited substantially higher organization variability, with 7, 7, 6, 10, 9, 17, 11, 6 and 13 distinct architectures, respectively. This extraordinary diversity reveals that centromeric satellite array organization is far more polymorphic than previously thought.

CEN04 represented a striking example of polymorphism driven by recurrent αSat-HSat1 transitions (**Fig. 1d**), revealing a complex evolutionary history of satellite array interleaving. Approximately 51.9% (*n* = 160) of centromeres contained a single HSat1 block inserted within the αSat array, while 19.5% and 21.1% (*n* = 62 and 66) harbored two and three discrete HSat1 blocks, respectively. Notably, the remaining 20 centromeres exhibited more complex αSat–HSat1 interleaved structures, with up to nine distinct HSat1 array insertions in two haplotype assemblies (**Supplementary Fig. 13**). Pairwise alignment dotplots enabled identification of duplicated segments within these arrays. Notably, duplicated segments were not localized to αSat-HSat1 junctions, and the inconsistencies in their boundaries and contents across individuals indicated that these duplications likely arose from recurrent independent events (**Supplementary Fig. 14**).

In contrast to CEN04, structural polymorphism in CEN01 was defined primarily by the absence, single-copy, or two-copy βSat arrays (**Supplementary Fig. 13**). A large inversion relative to T2T-CHM13 was dominant in the population with an AF of 0.81 across 271 haplotype assemblies, higher than previous estimate using Strand-seq^36^ (**Fig. 1c**). Among them, 127harbored singles while 92 contained two βSat arrays. The single βSat array is likely derived from a deletion event that fused two ancestral βSat arrays, rather than representing either ancestral array alone (**Supplementary Fig. 15**). We discovered a strong statistical association between the two seeming independent structural features. 87.3% (48 of 55) of centromeres carrying the αSat array inversion also harbored the single-βSat array configuration (*p* = 2.61×10^−6^, chi-square test), implying these two structural variants either arise on the same ancestral haplotype or are functionally linked through unknown mechanistic constraints (**Supplementary Table 10**).

### Large peri/centromeric inversions

Recurrent large inversions represent one of the SV forms richly clustered in peri/centrometric regions, driven through non-allelic homologous recombination between segmental duplications (SDs) that flank these dynamic genomic territories^17,36–38^. While previous studies have documented the existence of such inversions, their comprehensive characterization across diverse populations has remained limited by the lack of complete, high-quality centromere assemblies. Here, leveraging our high-quality peri/centromeric sequences, we systemically identified and characterized large inversions (>10 Kbp) within regions spanning each centromere plus 5 Mbp of flanking sequences on both arms. A final curated set of 194 peri/centromeric large inversions across 538 haplotype assemblies was identified, including two pericentric inversions that span both chromosome arms (**Supplementary Table 11**). These inversions are restricted to a subset of chromosomes, mainly occurring on chromosomes 2, 7, 9 and 16, while six chromosomes (3, 4, 5, 6, 8 and 18) show complete absence of large peri/centromeric inversions (**Supplementary Table 11**). The polymorphism of these inversions was primarily driven by sequence similarities between inverted SDs or palindrome structures, which were enriched at the breakpoints of large inversions (**Supplementary Fig. 16**). Compared to large inversions at the chromosome arms^24^ (*n* = 160), the peri/centromeric inversions had higher population frequencies and larger physical sizes, potentially reflecting weaker functional constraints of peri/centromeric SVs (**Supplementary Fig. 17**). Among all inversions, 19 are specific to EAS, including a 403-Kbp peri/centromeric inversion on chromosome 19 observed in 12 EAS assemblies (**Supplementary Fig. 18**; **Supplementary Table 11**).

The peri/centromere of chromosome 7 emerged as a hotspot of inversion activity, harboring 14 large inversions, including two rare >1-Mbp ones (**Fig. 2a**; **Supplementary Table 11**). Notably, one assembly C037-CHA-S17-01#Mat from APGp1 carries an ultra-large pericentric inversion (∼7.6 Mbp, 7p11-q11). This massive arrangement is independently observed in a single AFR assembly (NA20355#hap2 from HGSVC3; **Supplementary Fig. 19**). This large pericentric inversion was novelly identified among humans, and Nucfreq, GCI, GAIVISUNK and BioNano data collectively support the high confidence of assembly accuracy at this region (**Supplementary Fig. 20**). Alignments of long reads and Hi-C chromosomal interaction maps also provide clear inversion signals (**Fig. 2b**). A pair of inverted SDs were detected in the flanks of the inversion breakpoints, suggesting the homologous sequences have likely mediated this ultra-large rearrangement through ectopic recombination (**Supplementary Fig. 16**). Among the common inversions, several displayed population stratification in their allele frequencies (**Fig. 2c**). Whether these patterns arise from genetic drift or selection remains to be investigated.

**Figure 2.**
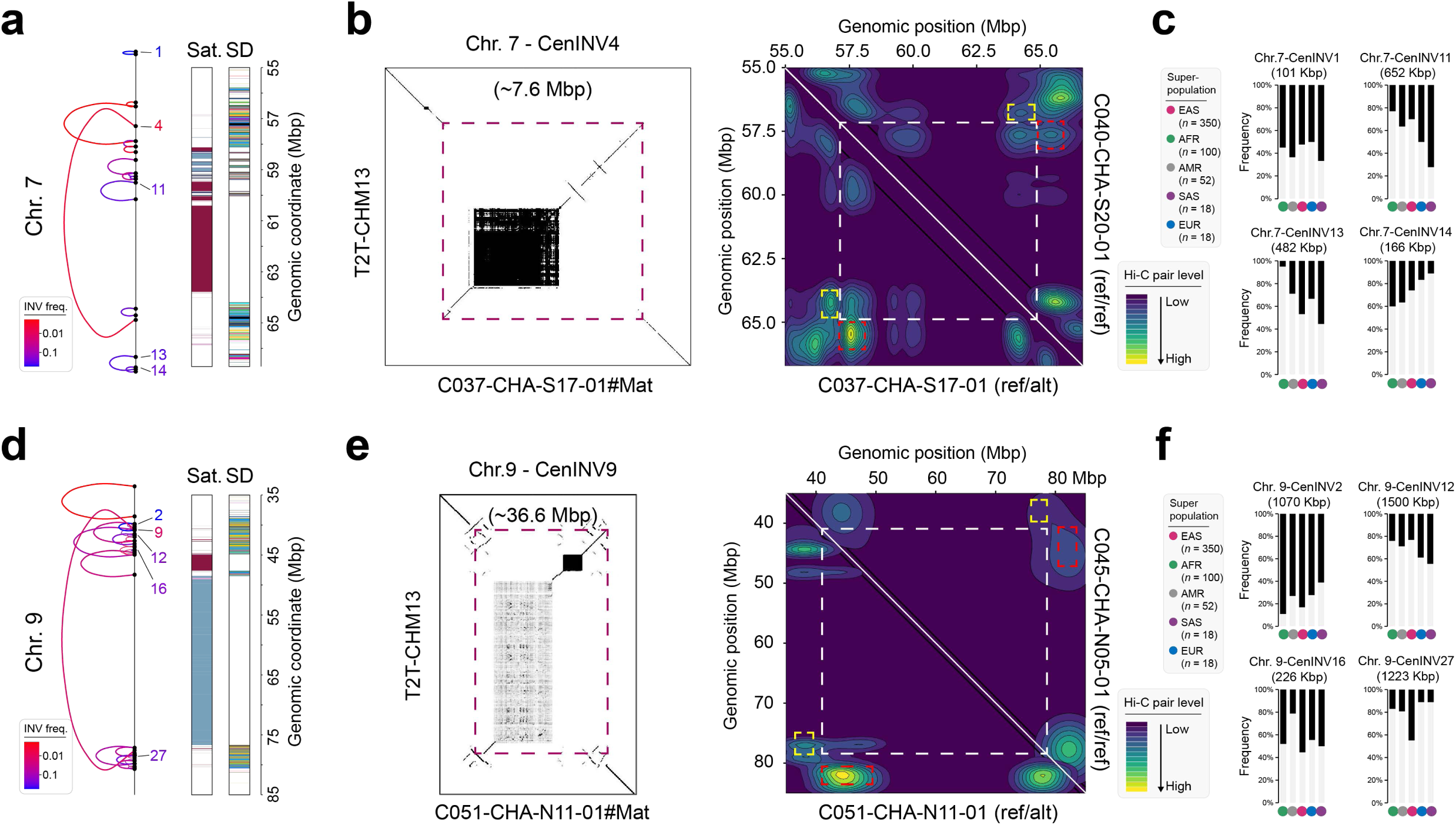
Recurrent peri/centromeric inversions on human chromosomes 7 and 9. **a** and **d**, Large inversions (>10 Kbp) mapped within the peri/centromeric regions of chromosomes 7 and 9. Two ends of each link represent the breakpoints of the inversion, with the color spectrum scaling with the inversion frequency (INV freq.) calculated across all investigated genomes. To provide structural context, the centromere satellite (Sat.) track and the segmental duplication (SD) track from T2T-CHM13 are shown for each region. Individual inversions are sequentially named according to their linear physical coordinates. **b** and **e**, Nucleotide sequence dotplots of ultra-large inversion regions and Hi-C-based interaction maps in CEN07 and CEN09. C040-CHA-S20-01 / C045-CHA-N05-01 (reference-allele carrier) and C037-CHA-S17-01 / C051-CHA-N11-01 (inversion-allele carrier) have distinct chromosomal interactions when Hi-C data are mapped against T2T-CHM13. **c** and **f**, Allele frequencies of the four most frequent peri/centromeric inversions on chromosomes 7 and 9, stratified by five global superpopulations. The dark bars denote frequencies of inversion alleles in each superpopulation.

Likely driven by multiple nested SDs, CEN09 presents an even more complex inversion landscape, with at least 32 peri/centromeric inversions, forming an intricate inversion complex (**Fig. 2d**; **Supplementary Table 11**). Within this complex, we identified another ultra-large pericentric inversion spanning ∼36.6 Mbp (p11-q21), detected heterozygously in ten individuals, including six EAS and four AFR assemblies (three from APGp1, two from HPRCy1 and five from HGSVC3), while notably no carriers were observed in AMR, EUR and SAS superpopulations (**Fig. 2e**; **Supplementary Figs. 19 and 21**). Previous investigations reported this inversion in ∼1% of the population and it is probably associated with infertility and repeated abortions^39,40^. In our dataset, we observed a frequency of 1.43% (5/349) in EAS and 3.09% (3/97) in AFR among unrelated individuals. Notably, we observed transmission of this inversion in two families (HG00512/HG00514 from EAS, and HG01890/HG01891 from AFR), where both the father and offspring carried the inversion in heterozygous state (**Supplementary Table 11**), indicating successful meiotic transmission across generations, despite its enormous size. This finding is particularly significant given the theoretical concerns about meiotic complications that might arise from pairing difficulties between inverted and non-inverted homologous during prophase I, especially for an inversion of this magnitude that encompasses the entire centromere^41^. Additionally, the CEN09 region displayed numerous smaller inversions that illustrate the ongoing structural dynamism of this chromosome, with several displaying population stratification, including a ∼1.2-Mbp inversion present at higher frequency in EAS compared to other ancestries (**Fig. 2f**). We observed no other previously reported pericentric inversions associated with abnormality^42^. The functional consequences of these peri/centromeric inversions—especially pericentric ones—on human health and disease represent a particularly intriguing area for further research.

### HOR diversity of human alpha satellites

Human αSats are organized in periodicity, forming HOR arrays, which can be further subdivided into distinct layers composed of HOR structural variants (StVs), the variant forms of canonical HOR units arising from internal monomer rearrangement^43^. To mitigate reference bias in annotating HOR units and arrays across multi-ancestry genomes, we developed a novel analytical workflow, aggregating ∼244 million αSat sequences from APGp1, HPRCy1, HGSVC3 and four reference-level complete assemblies (**Fig. 3a**; **Supplementary Note 2**). Leveraging a greedy, heuristic centroid-based algorithm, we assigned these sequences into 29,773 representative clusters, ensuring rare variants, population-specific variants and common sequences are all equally prioritized throughout the process (**Fig. 3a**; **Methods**). We then employed two complementary tools for HOR inference: graph-based HORmon^44^ and hierarchical tandem repeat mining (HTRM) method-based HiCAT^45^ (**Supplementary Note 2**). Of all defined monomer clusters, 3,722 were organized into HOR arrays, representing 95.8% of all αSat sequences. Notably, 2,197 clusters (59.0%) of these clusters are absent in T2T-CHM13, demonstrating the limited diversity of the single reference genome. Phylogeny of representative monomers revealed eight distinct groups, exhibiting 96.9% concordance with established suprachromosomal families (SF) taxonomy^18,19,46^ (**Supplementary Fig. 22**; **Supplementary Table 12**). We validated our clusterings by reconstructing known HOR arrays, and successfully recalled 57 of the 60 previously annotated HOR arrays in T2T-CHM13, six of which were merged with sister HOR arrays due to differences in detection methodology (**Supplementary Table 13**). The three undetected arrays had monomer divergence exceeding the identity threshold (**Supplementary Note 2**).

**Figure 3.**
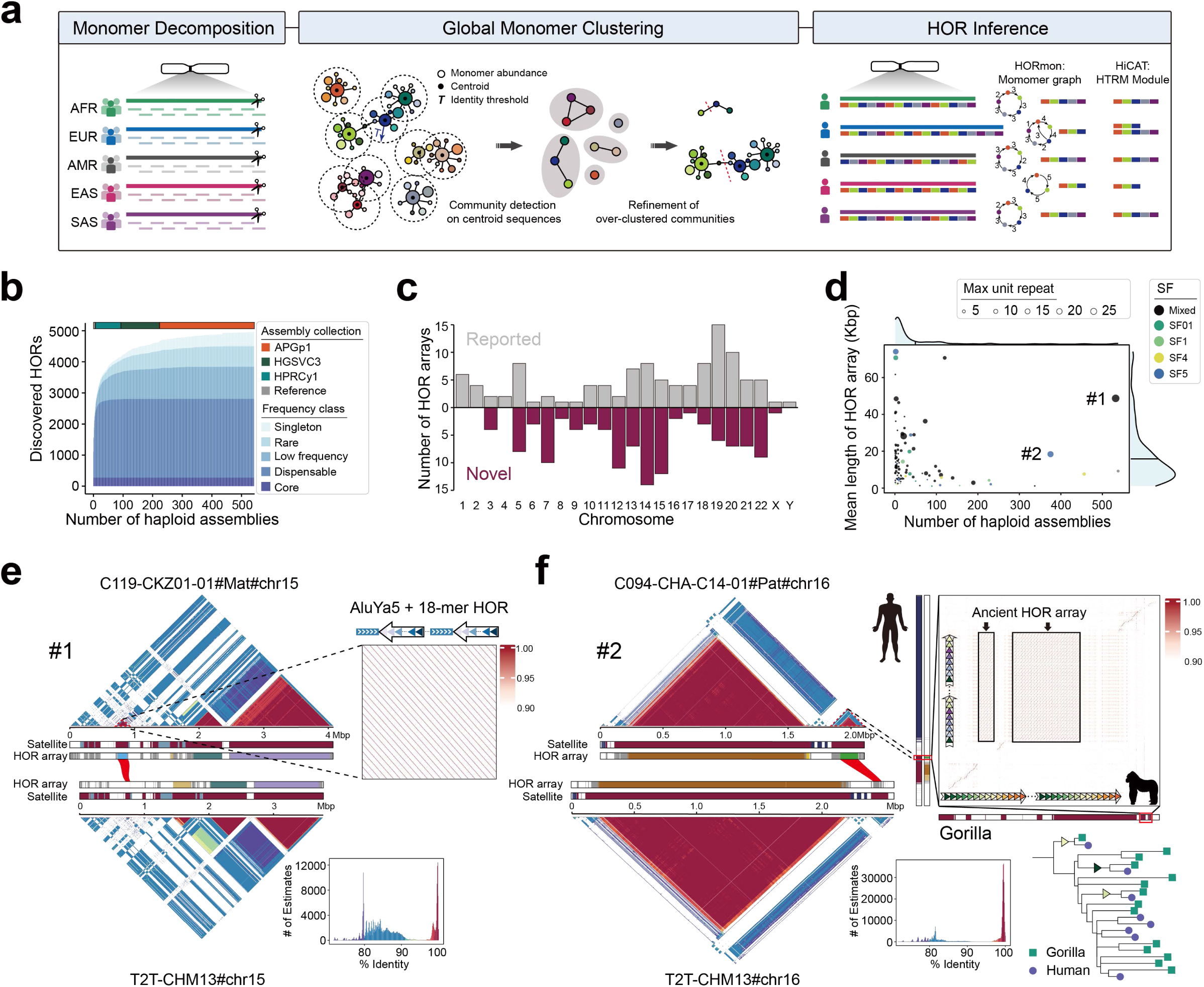
Pangenomic HOR profiling across 543 haplotype assemblies reveals ancient HOR arrays. **a**, Overview of HOR profiling workflow. This workflow includes monomer decomposition, global monomer clustering and HOR inference. Briefly, all αSat sequences from all assemblies are split and clustered into αSat monomer clusters. By utilizing graph and hierarchical tandem repeat mining (HTRM) based methods (HORmon and HiCAT), repeat strings with clustering labels are annotated as HORs. **b**, Growth curve of HOR StVs discovered from increasing number of haplotype assemblies by HORmon. Genome assemblies from APGp1, HPRCy1, HGSVC3 and reference-level assemblies are included. Five categories: core (>90%), dispensable (>5%), low-frequency (>1%), rare (present in multiple haplotype assemblies, but at <1% frequency) and singleton (observed in only a single haplotype assembly), are determined by population frequency. References include T2T-CHM13, T2T-CN1, Q100-HG002 and YAO. **c**, Chromosomal distribution of HOR arrays. Novel arrays unreported in T2T-CHM13 are highlighted in magenta. **d**, Correlations between average length of novel HOR arrays and its prevalence across haplotype assemblies. The suprachromosomal families (SFs) are highlighted by different colors. Arrays labeled as’Mix’ indicate that the monomers constituting the HOR array are derived from more than one SF.’#1’and’#2’denote the two cases illustrated in **e** and **f**, respectively. **e**, A novel CEN15 HOR array in assembly C119-CKZ01-01#Mat, absent in T2T-CHM13. This HOR unit is composed of AluYa5 and 18-mer. **f**, A novel CEN16 HOR array identified in the assembly C094-CHA-C14-01#Pat. Sequence identity maps generated by StainedGlass reveal that C094-CHA-C14#Pat contains an independent evolutionary layer with high intra-array identity, whereas the syntenic region in T2T-CHM13 lacks this layer. The zoom-in monomer identity heatmap (central track) shows clear co-linearity between the two species in the flanking regions, but in the central region each species exhibits its own expanded HOR array (an 8-mer array in human and a 14-mer array in gorilla). Different coloured triangles in the track denote distinct monomer variants. Phylogenetic tree constructed from the consensus sequences of all monomers. Three monomer clades show well-supported monophyly between human and gorilla sequences, indicating intermixing of orthologous monomers. These clades are marked by triangles on their ancestral branches, and their colors correspond to the monomer colors used in the HOR structure track above.

Across all assemblies, HORmon and HiCAT identified 4,971 and 5,512 HOR StVs in 113 and 195 HOR arrays, respectively (**Supplementary Tables 14** and **15)**. Accumulation curves of HOR StVs plateaued across assemblies, confirming the representativeness of our centromere sequences in capturing global HOR diversity (**Fig. 3b**; **Supplementary Fig. 23a**). APGp1 contributed 86.4% of all HORmon-annotated types, with 333 unique to this project (**Supplementary Fig. 24a**). Among HORmon StVs, 468 (9.4%) were singletons, and 274 (5.5%) were shared by >90% of haplotype assemblies (*n* > 489), constituting a core repertoire of human HOR units (**Supplementary Fig. 24b**). Core HORs exhibited significantly higher copy numbers than other frequency categories (*p* < 2.2×10^-16^, Wilcoxon signed-rank test; **Supplementary Figs. 23d and 24c**). However, rare HORs also showed dramatic individual expansions, exemplified by a 4-mer StV in CEN07 expanding over 1,300 times in an AFR assembly (NA19238#hap2) nested within the canonical 6-mer HOR array, highlighting individual-specific centromeric dynamism (**Supplementary Fig. 24d**). Among superpopulations, 488 HOR StVs exhibited significant copy number divergence between AFR and EAS haplotypes (*p* < 0.05, Wilcoxon signed-rank test), with functional and evolutionary consequences yet to be determined. Additionally, 396 and 431 HOR StVs were specific to AFR and EAS, respectively, including 47 common (AF > 0.05) in AFR and 24 in EAS (**Supplementary Table 14**).

Our pan-genomic approach enabled the discovery of 110 previously unreported HOR arrays absent from T2T-CHM13, predominantly localized to CEN14 (*n* = 14), CEN15 (*n* = 12), and CEN12 (*n* = 11), averaging 15.84 Kbp per haplotype and were shared among ∼49 haplotypes (**Figs. 3c** and **3d**; **Supplementary Table 16**). For example, a distinct CEN15 HOR array, present in 532 out of 543 assemblies (98.0%), formed a unique evolutionary structure (average length 48.7 Kbp) composed of an 18-mer αSat array along with an AluYa5 element (**Fig. 3e**). Majority (98/110) of novel HOR arrays originated from SF4, SF5, SF6 or their chimeric derivatives, representing evolutionary HOR relics of ancestral centromeric structures (**Fig. 3d**). Notably, a newly identified SF5-derived 8-mer HOR array in human CEN16 exhibited synteny with a 14-mer HOR array in gorilla^47^, with extensive monomer intermixing in phylogenies indicating shared ancestry predating the human-gorilla split approximately 10 million years ago (**Fig. 3f**). Flanking sequences of these arrays displayed high genomic synteny, supporting a shared evolutionary origin and subsequent lineage-specific expansion. No comparable expansion exists in chimpanzee and bonobo genomes^47^, potentially suggesting lineage-specific loss or incomplete sampling in these great apes (**Supplementary Fig. 25**).

### Inter-chromosomal HOR sharing landscape

Leveraging our comprehensive centromere assemblies and reference-free HOR annotation, we characterized the landscape of HOR arrays shared across human chromosomes. We identified 25 distinct HOR arrays shared by at least two chromosomes, with the majority clustering into two prominent centromere groups: CEN13/14/21/22 (*n* = 15) and CEN01/05/16/19 (*n* = 8; **Figs. 4a-4c**; **Supplementary Table 17**). Additionally, we detected two rare transchromosomal events, including a canonical inverted 16-mer HOR array (∼202 Kbp) translocating from CEN20 to CEN17 (**Supplementary Fig. 26**).

**Figure 4.**
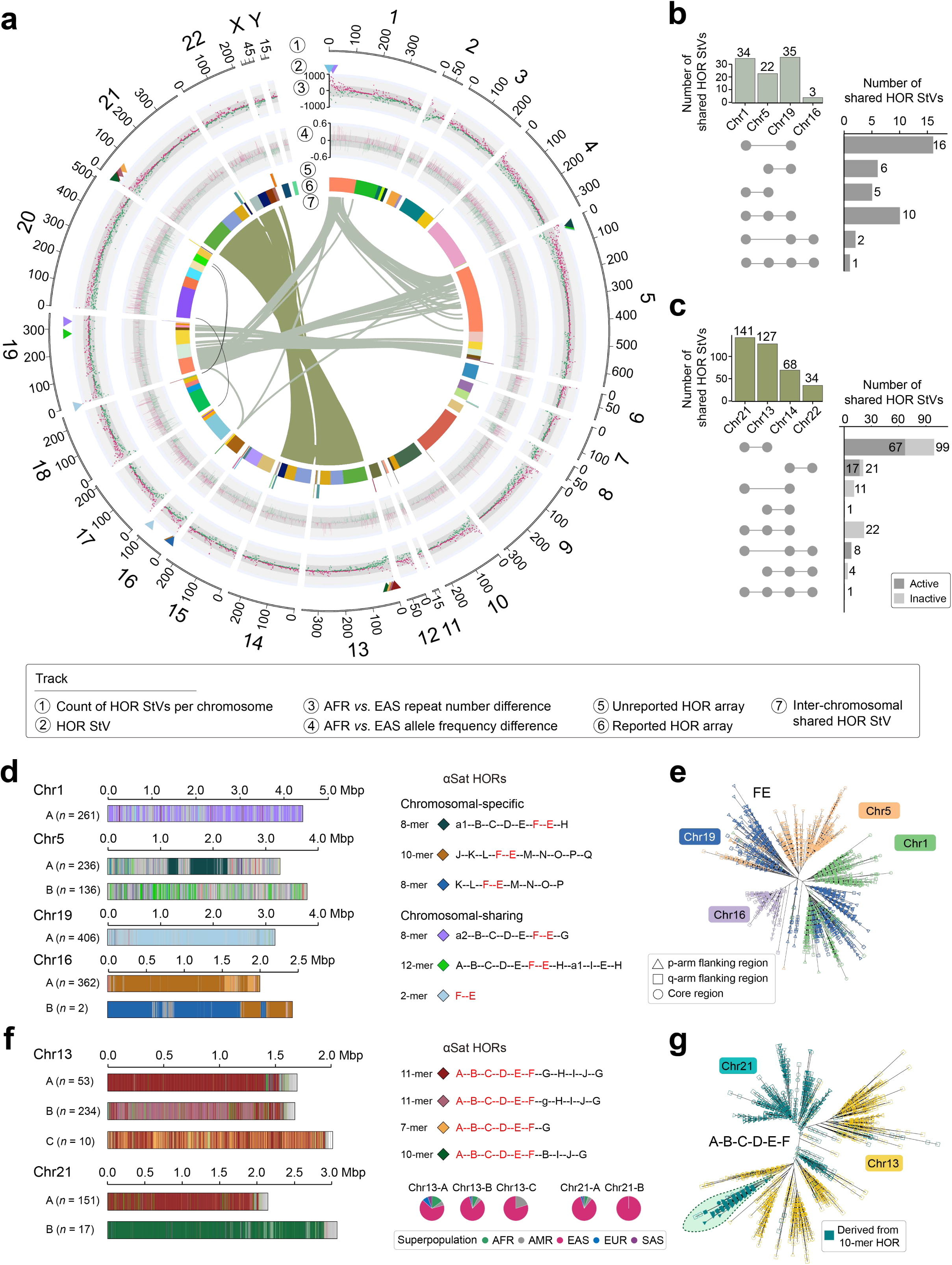
Inter-chromosomal sharing landscape sheds light on the evolutionary history of centromeres. **a**, Circos plot illustrating genome-wide inter-chromosomal sharing of HOR structural variants (StVs) annotated by HORmon. Concentric tracks from outer to inner: Track 1, number of HOR StVs per chromosome; Track 2, colored triangles represent HOR StVs shared between chromosomes, with colors matching those shown in **d** and **f**; Track 3, inter-population differences in mean StV repeat number between AFR (green) and EAS (magenta) superpopulations, plotted as log_10_-transformed absolute values. Shading denotes transformed ranges: dark gray<1, light gray 1-2.5, light blue 2.5-3; Track 4, StV allele frequencies in AFR and EAS. Background colors denote difference ranges: dark gray (<0.2), light gray (0.2-0.5), and light blue (0.5-0.6); Track 5, unreported HOR arrays, with distinct colors representing different HOR array; Track 6, previously reported HOR arrays, colored by array type; Track 7, connecting ribbons representing inter-chromosomal shared StVs. **b** and **c**, Upset plots quantifying intersections of shared HOR StVs for representative chromosomes. Dark grey bars denote StV sharing between active arrays; light grey bars denote sharing between inactive arrays. **b**, StV intersections for CEN01, CEN05, CEN16 and CEN19. **c**, StV intersections for CEN13, CEN14, CEN21 and CEN22. **d**, Major centromeric haplotypes (CenHaps) and dominant StVs for CEN01, CEN05, CEN16 and CEN19. StVs are depicted by diamond symbols, colour-matched to CenHap track color schemes at the left and to the shared-StV triangles in (**a**, Track 2). Distinct StV monomer compositions are labeled alphabetically (A-Q). A conserved 2-mer FE HOR (red) is shared across all four chromosomes. **e**, Phylogenetic reconstruction of the 2-mer FE HOR. Tip colours denote chromosomal origin. Shared-segment sequences from three regions of each active array are used: p-arm-flanking sequences (0-200 Kbp from the array start, indicated by triangles), core central sequences (midpoint ± 100 Kbp, circles) and q-arm-flanking sequences (terminal 200 Kbp, squares). **f**, Major CenHaps and dominant StVs for CEN13 and CEN21, highlighting a distinct haplotype in EAS characterized by near-complete replacement with a 10-mer HOR structure. StV symbols, colour coding and alphabetical monomer labeling follow the conventions defined in (**d**) (A-G). A 6-mer segment (A-B-C-D-E-F; hor_A-F, red) is shared among all four dominant StVs. **g**, Phylogenetic tree of partially shared hor_A-F sequences. Tip colours and region-based shape conventions (triangles, squares, circles) follow the scheme in (**e**). Filled symbols indicate sequences derived from the 10-mer HOR, while open symbols indicate sequences from the other three HOR StVs. The phylogeny reveals embedding of a CEN21-derived 10-mer HOR within CEN13 HOR sequences, indicative of inter-chromosomal transfer.

Within the CEN01/05/16/19 group, seven of the eight shared arrays were inactive and restricted to CEN05 and CEN19, while one active array was shared across all four chromosomes (**Fig. 4a**). Higher resolution analysis revealed that only a small fraction of StVs were shared between CEN05 and CEN19 (HORmon: 8.8%; HiCAT: 3.2%) in the five inactive arrays, indicating these arrays accumulated extensive chromosome-specific variants following an ancestral transfer event, rather than undergoing recent sequence exchange (**Supplementary Table 17**). In the shared active array, we identified both chromosome-specific StVs and sequence variants (SqVs; **Fig. 4b**; **Supplementary Note 2**). Alignments of shared HOR StVs revealed that CEN19 exhibited the highest SqV polymorphism, supporting a previous hypothesis that CEN19 represents the most ancestral centromere among these chromosomes^9^ (**Supplementary Figs. 27 and 28**). We further found a 2-mer HOR variant designated as FE (where F and E denote the two constituent monomer types) actively maintained across all four centromeres yet exhibiting copy number divergence spanning orders of magnitude, from 16 ± 20 in CEN16 to 8,977 ± 2,499 in CEN19 (**Supplementary Fig. 27b**). Phylogenetic analyses revealed that 2-mer HORs from CEN16 were nested within those from CEN19, with similar pattern observed for CEN01 and CEN05, suggesting the CEN16’s dimeric HORs originated from CEN19 (**Fig. 4e**). Additional evidence from reciprocal monomer incorporation between flanking regions of active arrays in CEN19 and CEN16, which was not observed between CEN16 and two other centromeres, further supported their closer evolutionary relationship (**Supplementary Fig. 29**). Notably, these FE HORs were preferentially retained as core components of active arrays in both CEN16 and CEN19, with subsequent amplification in CEN19 and CEN01, leading to extended dimer stretches (**Fig. 4d**).

In contrast, the CEN13/14/21/22 acrocentric group exhibited substantially higher sharing, with 15 arrays and 167 StVs (HiCAT; 391 for HORmon), collectively occupying >90% of the total length of HOR arrays per centromere (**Fig. 4c**; **Supplementary Table 17**). Importantly, CEN13/21 and CEN14/22 subclusters were largely distinguishable by their patterns of shared inactive and active arrays, respectively (**Figs. 4a** and **4c**). To determine whether this high homology arises from genetic drift or ongoing inter-acrocentric recombination, we identified chromosome-specific SqVs and reconstructed phylogenies using predominant HORs from active arrays. For CEN13/21, the canonical 11-mer HOR was universally shared across 239 complete assemblies. While shared StVs were common, we observed a 7-mer HOR (shared by 233 haplotypes) showing marked expansion (occupied over 30% of the active array) in 10 CEN13 haplotypes, whereas a 10-mer HOR (shared by 10 haplotypes) nearly completely replaced canonical structure in 17 CEN21 haplotypes (**Fig. 4f**). Notably, these 10-mers were EAS-specific (**Fig. 4f**). Phylogenetic analysis of randomly sampled shared HORs revealed distinct clustering for most CEN13 and CEN21 sequences, with one key exception: a monophyletic clade of 10-mer HOR sequences from CEN21, embedded within CEN13 HORs (**Fig. 4g**). This pattern, along with significantly reduced SNP divergence between the CEN21 10-mer HOR and the CEN13 11-mer HOR (37.0 ± 5.8) compared to the canonical CEN21 11-mer (38.2 ± 6.5), may reflect a potential transfer event from CEN13 active arrays to CEN21 (**Supplementary Fig. 30**). For CEN14/22, canonical 8-mer HORs showed nearly complete coverage of both active arrays and greater homogeneity than CEN13/21 (**Supplementary Fig. 31**). Unlike observations in T2T-CHM13, population-level phylogenies failed to distinguish CEN14 and CEN22 HORs, implying recent divergence, recurrent exchange, or selection-mediated conservation (**Supplementary Fig. 31**). These findings demonstrate that inter-chromosomal HOR sharing is not merely an ancient evolutionary relic, but an ongoing dynamic process. Acrocentric chromosomes, in particular, exhibit highly fluid sequence exchange—an event that continuously reshapes their centromeric architectures across human populations.

### Population stratification of centromere haplotypes

The extreme repetitiveness of centromeric satellite arrays, coupled with the scarcity of population-scale complete assemblies, severely constrains our understanding of their structural variation and evolutionary dynamics across human populations. Here, we developed HORSCAN, a computational tool that enables pairwise, single-satellite-aware collinearity analysis for αSat arrays by integrating satellite clustering categories and HOR structure information (**Methods**). Using synteny distance to quantify pairwise centromere similarity, we classified centromere sequences into discrete centromere haplotypes (CenHaps; **Supplementary Fig. 32**). Across chromosomes, two primary architectural features that define CenHaps were generally observed: variation in HOR array size (e.g., CEN03, CEN07, CEN12), and differences in HOR StVs (e.g., CEN04, CEN05, CEN08, CEN10 and CEN17; **Supplementary Figs. 32-40**). Inferring genetic relationships among size-driven CenHaps is hindered by recurrent independent expansion and contraction in αSat HOR arrays, whereas StV-driven CenHaps offer stable molecular signatures for detailed evolutionary history reconstruction and population stratification characterization. For chromosomes with pronounced HOR StVs, we performed unsupervised, alignment-free *k*-mer clustering as an independent validation and found high concordance with our assigned CenHap classification (**Supplementary Fig. 41**).

Taking CEN08 as an example, we identified three major CenHaps with distinct ancestry-specific distribution (**Fig. 5a**; **Supplementary Fig. 33**). CenHap-A was predominantly characterized by 7-mer and 19-mer HOR StVs, with the 7-mer forming a haplotype-specific evolutionary layer. CenHap-B, present in 205 assemblies, was defined by a canonical 11-mer (D8Z2) at the periphery of active arrays and a distinct layer of a 15-mer HOR StV. CenHap-C was identified across 20 assemblies, predominantly of AFR ancestry (*n* = 19), and exhibited a pronounced contraction of the canonical 11-mer HOR (**Figs. 5b** and **5c**; **Supplementary Fig. 33**). Clear ancestry-specific differentiation was observed: EAS lineages were enriched in CenHap-A and its subtypes (A1-A3), while AFR assemblies were predominantly in CenHap-B and CenHap-C, particularly subclades B2, B3, and C2, reflecting divergent demographic histories (**Fig. 5d**; **Supplementary Fig. 33**). Phylogenetic reconstruction utilizing flanking sequences with chimpanzee and bonobo outgroups dated the divergence between CenHap-A and CenHap-B to approximately 251-399 thousand years ago (**Supplementary Fig. 34**). Additionally, phylogenetic analysis of HOR sequences and genomic distribution identified the 11-mer HOR unit as the ancestral configuration, which gave rise to 7-mer (mn_A-G) and 8-mer (mn_A-C_G-K; **Fig. 5b**; **Supplementary Fig. 33**). These variants subsequently coalesced into the mosaic 15-mer (mn_A-G_A-C_G-K) characteristic of CenHap-B, while the 7-mer variant experienced localized expansion, further diversifying the centromeric landscape (**Supplementary Fig. 33**).

**Figure 5.**
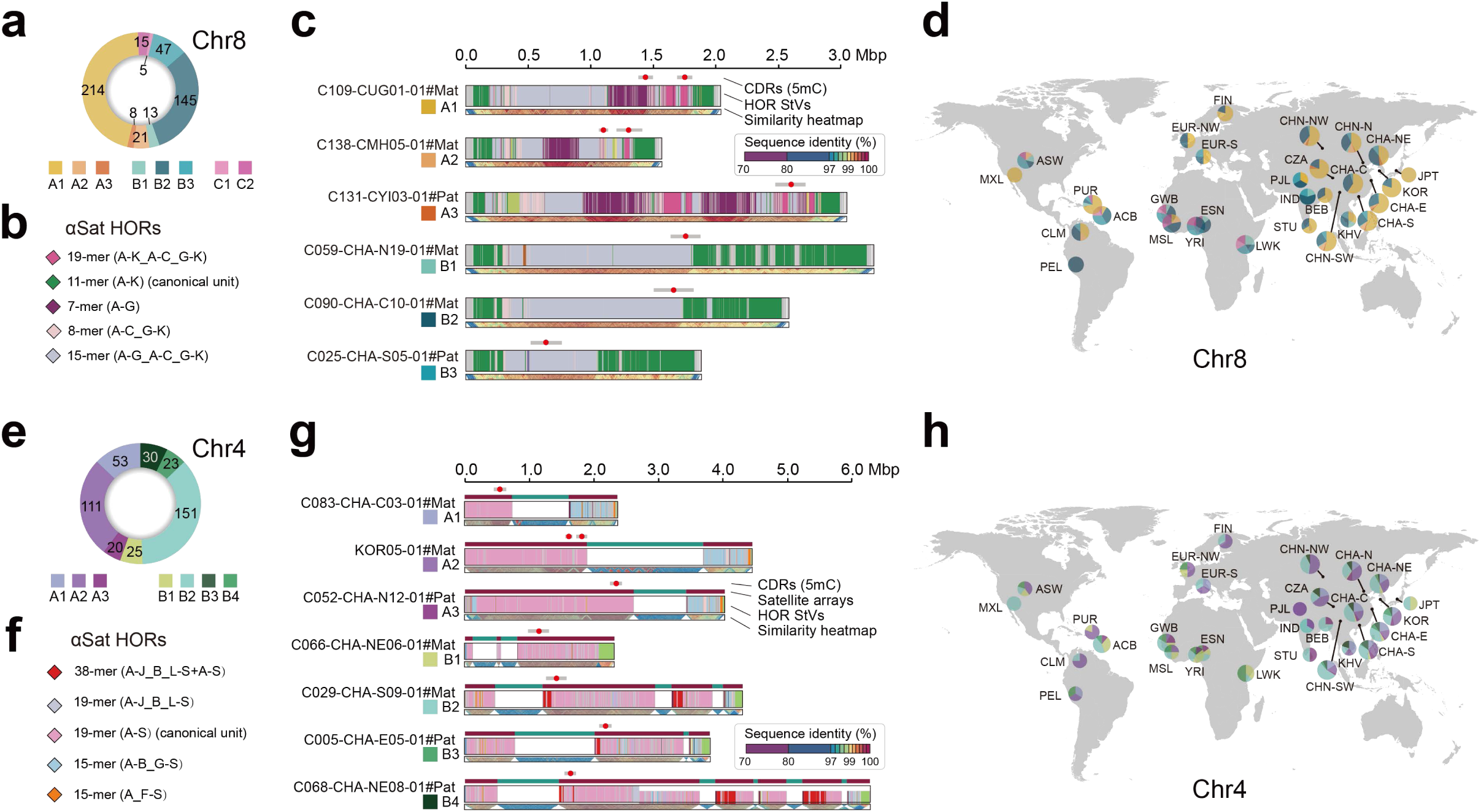
Population stratification of synteny-based centromere haplotypes. Number of haplotype assemblies assigned to CenHaps for CEN08 (**a**) and CEN04 (**e**). Major classes (A-C) reflect distinct HOR structural variants (StVs) layer composition; subclasses (e.g., A1-A3) are further defined using synteny distance. **b** and **f**, Major HOR StVs in CEN08 (**b**) and CEN04 (**f**), depicted as diamond symbols. Monomers are denoted by alphabetical labels to reflect distinct structural features. The canonical units represent the inferred ancestral monomer arrangement from which contemporary centromeric HOR arrays evolve. Continuous letter ranges (e.g., A-G) denote uninterrupted contiguous monomer arrangement, while underscored discontinuous ranges (e.g., A-C_G-K) denote the concatenated non-adjacent segments, with internal monomer deletions (D-F deletion in the example). **c** and **g**, Representative full-length CenHap examples for CEN08 (**c**) and CEN04 (**g**). Tracks display the centromere dip regions (CDRs), along with sequence similarity heatmaps (representing local homogenization). For CEN04, blank intervals in the HOR StVs track colocalize with HSat1 satellite tracks, demonstrating the association between HSat1 array insertion frequency and CenHap classification, with enriched insertions in CenHap-B. **d** and **h**, population-level CenHap composition for complete centromeres of CEN08 (**d**, *n* = 468) and CEN04 (**h**, *n* = 413). Pie charts show CenHap frequency distributions across human populations, with color schemes consistent with the corresponding CenHap classification panels (**a** and **e**). Detailed population descriptions are provided in Methods.

CEN04 exhibited pronounced CenHap bifurcation into two clusters (**Fig. 5e**), distinguished by HSat1A block insertions. 84.7% (194/229) of CenHap-B haplotypes contained ≥2 αSat-HSat1A interleaved structures, *versus* a single insertion in 98.4% (181/184) of CenHap-A carriers (**Figs. 5f** and **5g**; **Supplementary Fig. 35**). AFR samples were highly enriched in CenHap-B, while other superpopulations show balanced distribution. Notably, the EAS-enriched subcluster B4 had frequent HSat1A insertions, suggesting population-specific duplication (**Fig. 5h**). These two CenHaps diverged approximately 321-553 thousand years ago (**Supplementary Fig. 36**), establishing them as evolutionarily stable configurations maintained across human populations through selective or demographic forces that remain to be fully elucidated.

Additionally, certain CenHaps were characterized by inactive array insertions splitting active arrays, predominantly in CEN17 and CEN19 (**Supplementary Figs. 37-40**). In CEN19, insertions of two inactive arrays were identified in 42 haplotype assemblies, consistent with prior reports^3,17^. In CEN17, insertions occurred both in 65 CenHap-A and 6 CenHap-B haplotypes, with strong population specificity: 61/65 CenHap-A carriers were from EAS superpopulation, and all 6 CenHap-B carriers were from AFR (**Supplementary Fig. 37**).

### Epigenetic dynamics among centromeres

The epigenetic state of centromeres governs kinetochore assembly during chromosome segregation, making it essential for maintaining genome stability^48,49^. In most eukaryotes, the centromere-specific histone variant CENP-A serves as the definitive marker of functional centromeres^43,50–52^, with its occupancy strongly correlating with locally hypomethylated regions termed centromere dip regions (CDRs). These CDRs are primarily localized within highly homogenized HOR arrays^9,12,53^, yet their dynamics across individuals and populations remains poorly characterized. To map these functional centromeric regions, we performed CENP-A CUT&Tag sequencing on induced pluripotent stem cells (iPSCs) from five individuals and leveraged the CpG methylation kinetics signals of blood samples from PacBio HiFi sequencing to define CDRs across all 160 APGp1 individuals (**Supplementary Table 18**). Remarkably, 90.2% of centromeres (165 out of 183) exhibited high consistency between the CENP-A peaks and CDRs, validating methylation-based CDR calling as a robust proxy for kinetochore positioning (**Fig. 6a**; **Supplementary Fig. 42**; **Supplementary Table 18**). It should be noted that current CENP-A enrichment peaks represent an underestimate due to limits of short-read alignments in distinguishing extremely homologous inter and intra-centromere regions for a small subset of chromosomes (e.g., chromosome Y; **Supplementary Table 19**).

**Figure 6.**
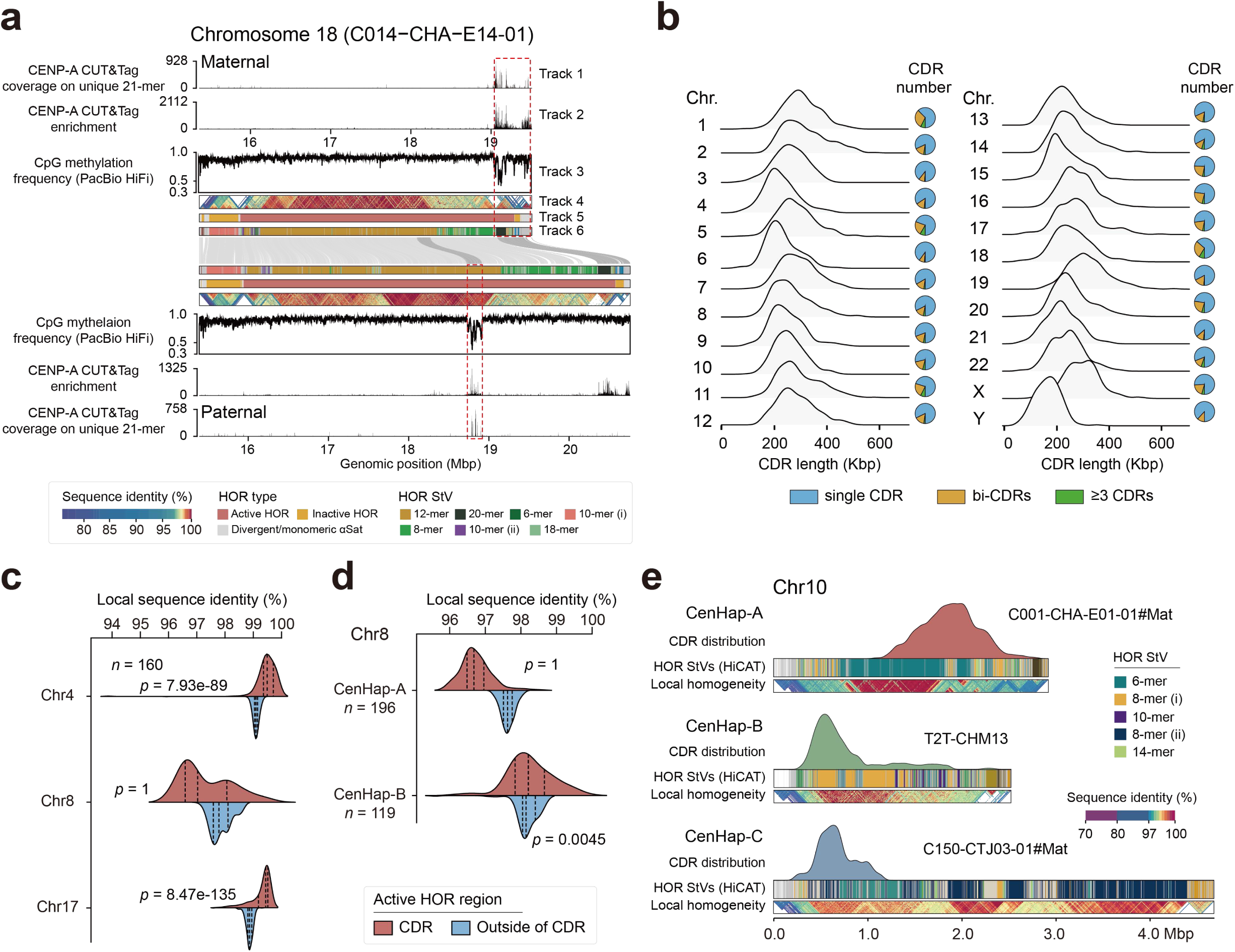
Variations in human centromeric CDR size and positioning. **a**, Centromeric sequence and epigenetic patterns of two haplotypes in C014-CHA-E14-01. Track 1: CENP-A CUT&Tag read coverage restricted to inter-haplotype unique 21-mers; Track 2: CENP-A CUT&Tag enrichment, calculated as CENP-A CUT&Tag read coverage normalized to the IgG control; Track 3: 5mC methylation frequency; Track 4: sequence similarity plot; Track 5: αSat family annotation; Track6: HOR StV annotation. The central links demonstrate alignment between the two haplotypes. The apparent secondary CENP-A peak at the distal end of the paternal haplotype in the standard mapping (Track 2) is absent from the unique-21-mer-anchored track (Track 1), confirming it as a short-read mapping artifact arising from αSat sequence homogeneity rather than a secondary kinetochore site. **b**, For each chromosome, the left plot shows the distribution of CDR lengths, and the right pie chart illustrates the number of CDRs. **c**, Violin plots depict the local similarity of CDR and non-CDR active HOR regions of three chromosomes. One-tail student’s *t* test is performed for each chromosome. **d**, Violin plots show local similarity of CDR and non-CDR active HOR regions of two different CenHaps of chromosome 8. **e**, CDR location distribution for each CenHap of chromosome 10 is displayed in 3 tracks. Track 1: CDR distribution after lifting individuals of a CenHap to the corresponding reference genome; Track 2: HOR StVs annotation; Track 3: StainedGlass sequence identity plot.

CDR sizes varied substantially across chromosomes, with median lengths ranging from ∼160 Kbp for CENY to ∼326 Kbp for CEN19 (**Fig. 6b**). They displayed a stronger correlation with HOR array lengths (Spearman correlation *r* = 0.54, *p* = 0.0076), than the overall chromosome lengths (*r* = 0.26, *p* = 0.22; **Supplementary Fig. 43**). While most chromosomes harbored a single CDR, we detected additional CDRs (located >80 Kbp from the primary CDR) on 1,482 chromosomes (16.8%), specifically on chromosomes 1 and 18 (**Fig. 6b**). The recurrence of bi-or multi-CDR configurations, as evidenced by individuals from several haplogroups displaying bi-CDRs on the Y chromosome (**Supplementary Fig. 44**), implies this may represent an underappreciated yet common feature of human centromere biology.

As expected, almost all CDRs (99.94%) mapped to active HOR arrays, with local sequence identity within CDRs significantly higher than the whole active HOR regions (*p* = 9.12×10^-97^, one-tailed Student’s *t* test), highlighting the preference of kinetochore assembly on the youngest, most recently expanded and homogenized repeats (e.g., CEN04 and CEN17 in **Fig. 6c**; **Supplementary Fig. 45**). However, we also identified exceptions that violated this pattern. In CEN08, contrasting with CenHap-B where CDRs localized to recently expanded regions, 94.4% of CenHap-A centromeres positioned their CDRs outside these expansion zones, near the active array edge (**Fig. 6d**; **Supplementary Fig. 46**). Further exceptions included three centromeres from chromosomes 6 and two from chromosome 18, whose CDRs localized outside of active HOR regions (**Supplementary Fig. 47**), suggesting non-active HOR αSat segments may have the potential to serve as kinetochore assembly sites, although additional experimental evidence will be required to establish this possibility.

To characterize CDR positioning dynamics across individuals, we aligned CDR coordinates based on centromere synteny maps (**Methods**), revealing that ten chromosomes exhibited unimodal CDR distributions while others displayed bimodal or multimodal peaks, implying divergence of functional centromere localization (**Supplementary Fig. 48**). This dynamic distribution landscape was further confirmed by the CDR positionings from HPRCy1 and HGSVC3 using lymphoblastoid cell lines (LCLs; **Supplementary Fig. 49**; **Methods**). Interestingly, in CEN03, most centromeres had CDRs to the left of the megabase-scale HSat1A array, but several displayed CDRs to the right, suggesting the relatively loose constraints on kinetochore positioning (**Supplementary Fig. 48**). Moreover, we observed strong associations between CenHap categories and CDR distributions. In CEN10, for instance, lifting CDRs against their respective CenHap references yielded a single peak per CenHap, with CenHap-A CDRs clustering around the right boundary of long canonical 6-mer HOR arrays, while CenHap-B haplotypes showing CDRs near the p-arm, a region dominated by 8-mer StVs (**Fig. 6e**). These findings revealed how structural haplotype variation directly influences the epigenetic specification of functional centromeres.

### Centromeric recombination and single-base *de novo* mutation

Centromere regions have long been recognized as recombination deserts^54,55^, yet the precise dynamics of genetic exchange and linkage patterns across these highly repetitive territories have remained elusive, due to the historical absence of complete centromeric sequences and reliable variant calling from short reads^56,57^. Leveraging our comprehensive collection of gapless centromere assemblies and high-confidence graph-decomposed variants^24^, we systematically characterized recombination patterns across centromeres, by measuring genetic linkage among centromeres and their flanking regions (**Fig. 7a**).

**Figure 7.**
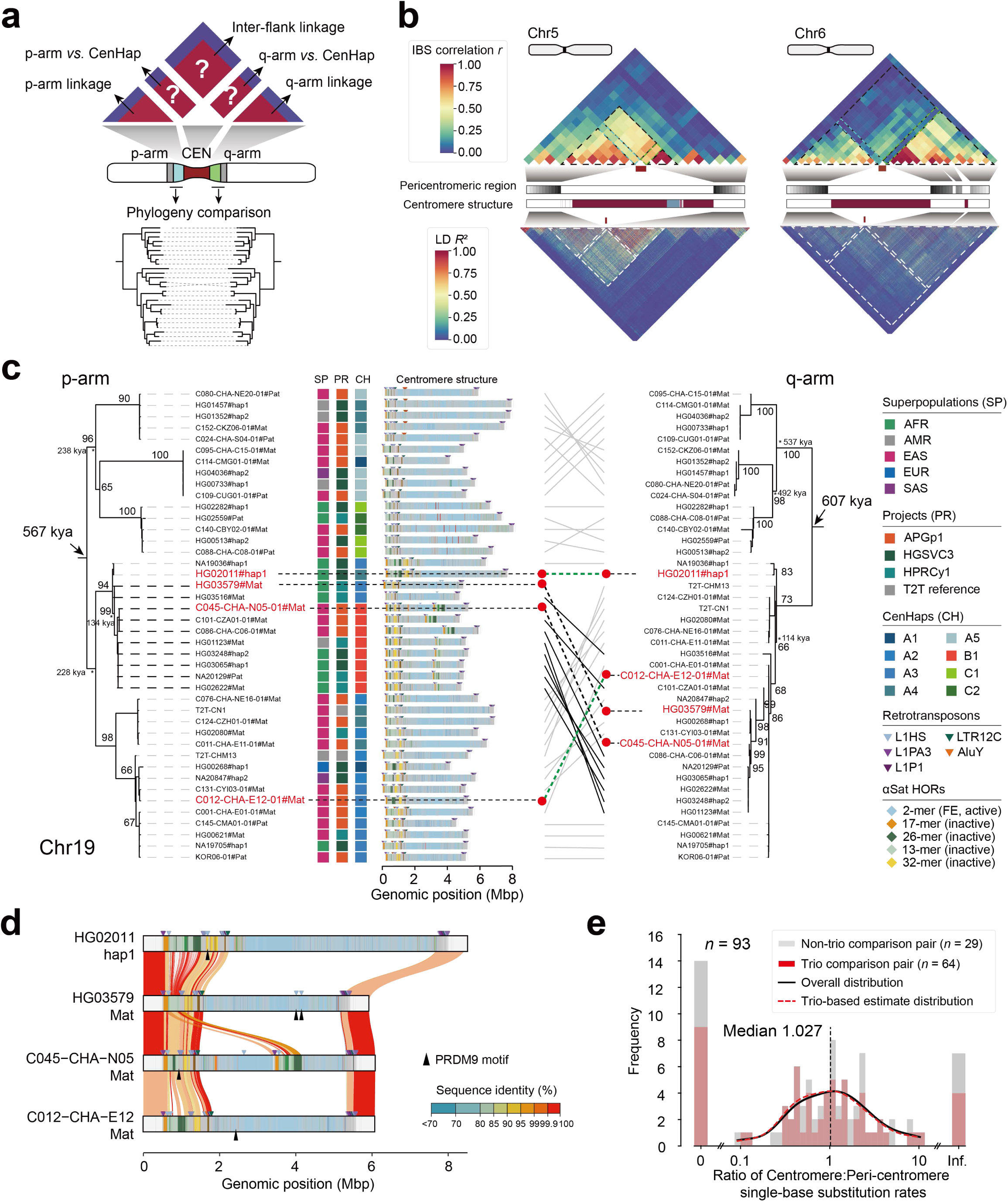
Centromeric recombination and *de novo* single-base substitution. **a**, The strategy to investigate pericentromeric genetic linkage pattern and phylogeny of flanking unique sequences from both p-and q-arms. The upper panel displays a correlation heatmap divided into five sections: p-arm linkage, q-arm linkage, inter-flank linkage, p-arm *versus* CenHap, and q-arm *versus* CenHap. For the first three sections, pairwise IBS scores were computed for each 100-Kbp window across all samples, and the resulting inter-sample IBS vectors were correlated between every pair of windows to generate a symmetric matrix reflecting population-wide linkage patterns across genomic positions. For q-arm *versus* CenHap and p-arm *versus* CenHap correlations, pairwise IBS distances from 100-Kbp windows were correlated with pairwise synteny distances quantified by HORSCAN. **b**, Distinct pericentromeric genetic linkage architectures exemplified by CEN05 and CEN06. Each centromere profile contains two defined subpanels: upper, inter-window IBS correlation heatmap; lower, SNP-level pairwise linkage disequilibrium (LD, R²). CEN05 exhibits strong linkage within and between pericentric flanking regions. In contrast, CEN06 displays attenuated inter-flank linkage, with sharp LD decay at the boundary between pericentromeric region and centromere. **c**, Maximum-likelihood phylogenetic trees for the p-and q-arms of CEN19 show discordant topologies, highlighted by red lines. Nodes with 60%-100% bootstrap support are indicated numerically. **d**, A comparative alignment of CEN19 sequence, including 500 Kbp pericentromeric region, visualized using SVbyEye^69^, reveals discordance in p-and q-arm sequence identity and shows the presence of PRDM9 motifs that colocalize with L1HS elements within the centromere. **e**, Quantitative analysis of single-nucleotide substitution rates across centromeric and pericentromeric regions, based on 93 centromere pairs.

We first calculated the correlations of pairwise identity-by-state (IBS) scores in 100-Kbp windowed flanking regions of active αSat arrays to quantify pericentromeric genetic linkage (**Fig. 7a**; **Methods**). As expected, we observed high linkage in both p-arm (mean Pearson’s correlation coefficient *r* = 0.61, ∼628.4 Kbp) and q-arm (mean *r* = 0.68, ∼560.6 Kbp) pericentromeric regions for euchromosomes (**Supplementary Fig. 50**). Chromosome X showed ever more extensive linked intervals, reaching ∼1 Mbp on the p-arm (mean *r* = 0.89) and ∼1.4 Mbp on the q-arm (mean *r* = 0.67), consistent with its unique evolutionary history and reduced effective population size. However, beneath this overall pattern of recombination suppression, we uncovered a striking asymmetry and chromosome-specific variations that challenge simplistic models of uniform centromeric recombination deserts. Marked asymmetry characterized the genomic span of linked regions across centromeres: CEN03, CEN04, CEN05 and CEN22 exhibited longer p-arm flanks (>300 Kbp) than q-arm counterparts, while CEN06, CEN07, CEN18, CEN20 and CENX showed the opposite pattern (**Fig. 7b**; **Supplementary Figs. 50-52**).This asymmetry may reflect differential recombination suppression, non-central kinetochore positioning, or historical directional gene conversion, though the underlying mechanism remains unresolved.

Given severe recombination repression across active HOR arrays, we hypothesized strong co-inheritance between CenHaps and flanking genetic variations^57^. We calculated correlation between pairwise genetic distances of centromeres (synteny scores in αSat HOR arrays) and pericentromeric genetic distance (IBS) to profile centromere-proximal variant associations (**Fig. 7a**; **Methods**). Seven chromosomes exhibited significant correlations (mean *r* > 0.3, *p* < 0.05) with at least one flank (**Supplementary Fig. 50**), while centromeres with weak correlations were primarily classified by length-driven CenHaps (12/16; e.g., CEN07 and CEN11; **Supplementary Fig. 30**). For chromosomes with HOR StV-driven CenHaps, phylogenetic trees constructed from flanking sequences showed largely congruent topologies for p-and q-arms (**Supplementary Fig. 50**).

To quantify p-arm and q-arm tree concordance, we assessed three complementary metrics: normalized Mutual Clustering Information (MCI), quartet agreement, and Spearman correlation of pairwise identity-by-state distances. As a positive control, chromosome Y, where the absence of recombination forces p-and q-arm sequences to share a single genealogy, yielded the highest scores across all three metrics (MCI = 0.602, quartet agreement = 1.000, IBS *ρ* = 0.816, Mantel *p* < 1×10⁻⁴). Most chromosomes showed robust inter-flank correlation, however, 7 chromosomes fell below the genome-wide mean for all three metrics including CEN01, CEN02, CEN03, CEN06, CEN07, CEN13 and CEN19, suggesting localized disruption of linkage patterns (**Fig. 7b**; **Supplementary Fig. 50**). Among these, the signal on chromosome 19 is attributable to a single clade conflict (**Fig.7c**; **Supplementary Fig. 53**).

Specifically, 49 CEN19 haplotypes, including 36 CenHap-A and 13 CenHap-B carries, formed a distinct monophyletic clade in the p-arm-flank phylogeny that conflicted with the q-arm topology (**Fig. 7c**). Sequence identity analysis confirmed this incongruence, supporting ancient recombination events that decoupled centromeric and flanking haplotypes and suggesting that even within recombination-suppressed regions, rare but consequential genetic exchange can reshape evolutionary trajectories (**Fig. 7d**). Importantly, we identified two high-density binding sites of PRDM9 in CEN19, a key protein in meiosis that defines recombination hotspots^58,59^, that precisely colocalized with truncated L1 elements bearing poly(A) tails (**Fig. 7d**; **Supplementary Fig. 54**). We speculate that L1HS insertion in CEN19 may provide potential PRDM9 binding sites, which may have mediated non-allelic homologous recombination. However, precise mapping of recombination breakpoints within centromeres remains challenging due to repetitive sequence expansion and contraction.

Centromeric regions have long evaded precise single-base mutation rate measurements due to their dramatic structural dynamics. To address this challenge, we established a stringent analytical framework by selecting 93 centromere pairs with >99% identity for reliable substitution rate estimation at single-base resolution (**Supplementary Table 20**). Paralogous αSat pairs between centromeres arising from segmental conversion, local expansion and other structural rearrangements, were excluded to ensure true orthologous relationships (**Supplementary Note 4**). For each centromere pair, we compared substitution frequencies between active αSat HOR arrays and flanking pericentromeric regions to test whether centromeres exhibit evaluated substitution rates than chromosome arms. Contrary to longstanding assumptions of centromeric hypermutation, we discovered that substitution frequencies within αSat HOR arrays were of the same order of magnitude as those in adjacent pericentromeric sequences across all chromosomes (**Fig. 7e**). The distribution of centromere-to-pericentromere substitution ratios centered on unity with a median of 1.027, showing no statistically significant deviation from equal mutation rates (*p* = 0.70, Wilcoxon signed-rank test). These findings were specially confirmed by 64 parental-offspring inter-generational centromere pairs alone (**Fig. 7e**). It should be noted that substitution rates varied over 100-fold across chromosomes. Notably, 14 centromere pairs exhibited a complete absence of substitutions (**Supplementary Table 20**). CDRs seemed to have a lower substitution rate than non-CDR regions among 20 analyzed centromere pairs, in which 16 harbored exactly zero single-base substitutions within the CDRs. However, these estimates are based on a limited number of paired centromeres and relatively short CDR sequence lengths, and should therefore be interpreted with caution (**Supplementary Table 21**). Collectively, these results fundamentally challenge the long-held notion that centromeric sequences evolve under relaxed mutational constraints, instead revealing that centromeres are subject to the same baseline single-base substitution rates as the broader genome.

## Discussion

This study marks a pivotal advance in human centromere genomics and genetics, delivering the most comprehensive atlas of complete centromere sequences and their multidimensional variation landscapes reported to date with over 8,000 gapless assemblies. By developing a reference-free annotation pipeline, we refined the repertoire of human αSat sequences and identified 110 previously uncharacterized HOR arrays. Complementing this,we employed a collinearity-aware algorithm that integrates monomer-level alignments with HOR structural features, enabling us to resolve the extraordinary structural plasticity of human centromeres among population scales, with massive large-scale inversions, complex interchromosomal exchanges, and rapid HOR turnover events. Additionally, our data reveal widespread population differences in centromeric architectural variation, encompassing sequence size, inversion profile, CenHap, and kinetochore assembly positioning (as represented by CDRs). A critical open question emerging from these observations is whether such stratification arises primarily from genetic drift or is shaped by natural selection^60^. Distinguishing these will require pedigree-based telomere-to-telomere assemblies that capture *de novo* centromeric changes, together with direct functional assays of kinetochore specification and segregation fidelity. Collectively, our findings illuminate the multi-level structural variations that underpin centromere architectural diversity.

Conceptually, the discovery of extensive interchromosomal HOR sharing networks, particularly active exchanges among acrocentric chromosomes, recasts centromeres as dynamic genomic territories engaged in ongoing sequence exchange rather than isolated repetitive islands. While prior work suggested severely suppressed recombination across centromeres^55^, we provide evidence for both ancient recombination events (evidenced by topological discordance in CEN19 flanks coincident with PRDM9 binding motifs) and contemporary inter-chromosal HOR transfers, including a phylogenetically-supported transfer from CEN13 to CEN21 primarily observed in EAS superpopulation. Furthermore, approximately 16.8% of chromosomes harbor multiple potential CDRs, with some separated by over 1 Mbp, and specific CenHaps directly influence the site of kinetochore positioning during chromosome segregation. This reveals a level of functional flexibility never before documented at the population scale. Combined with the ability of centromeres to tolerate massive structural rearrangements while preserving conserved function, this plasticity in kinetochore positioning points to an unexpected robustness of centromeres that may serve as an evolutionary buffer, allowing populations to explore diverse centromeric architectures while safeguarding essential chromosome segregation machinery. These findings reposition human centromeres not as static structural elements, but as dynamic evolutionary playgrounds where sequence, structure, and epigenetic specification intersect to shape both evolutionary trajectories and clinical phenotypes. However, fully deciphering the functional and evolutionary consequences of centromeric architectural diversity remains challenging due to current limitations in both centromere-resolved structural variant detection and epigenomic profiling of repetitive satellite arrays. Comparisons with an independent population-level centromere study^61^ further illustrate that different analytical frameworks can provide complementary perspectives on centromere diversity. Although the two studies quantify HOR diversity, haplotype classification, kinetochore-associated regions, and evolutionary dynamics using distinct definitions and approaches, comparative analyses reveal both shared patterns and framework-dependent differences that inform interpretation of human centromere variation (**Supplementary Note 5**). Future advances integrating centromere-aware structural variant frameworks, pangenomic assemblies from multiple cohorts, and long-read epigenomic technologies such as DiMeLo-seq^62^, will be essential for systematically resolving centromeric rearrangements and their associated chromatin landscapes across diverse human populations.

Compared to structural dynamism, their single-nucleotide substitution rates remain relatively constrained, which we found to be comparable to those of flanking pericentromeric regions. This dual-track evolutionary model, in which structural innovation proceeds alongside baseline point-mutational constraint, reconciles decades of conflicting observations and establishes a new framework for understanding how centromeres balance functional conservation with architectural diversity. However, it is important to note the relative substitution rates we estimated have a broad range of variation (>100-fold) across chromosomes, which requires more pedigree-based assemblies to quantify accurate intergenerational substitution rates^63^. Additionally, our estimate quantifies orthologous single-base substitution after excluding structural rearrangements and non-orthologous paralogues, and is therefore complementary to concurrent estimates of total array divergence and *de novo* variation that additionally capture gene conversion and unequal exchange^61^; jointly, these metrics describe the distinct point-mutational and concerted-evolution components of centromere turnover.

Importantly, the establishment of reference ranges for human parental variation in centromeric sequences and configuration provides a quantitative framework for assessing potential risks of meiotic stability and centromere-associated diseases. Emerging evidence has linked centromeric variation to cancer and human disorders (e.g., birth defects, fibrosis)^21–23^. The documented transmission of ultra-large inversions (particularly pericentric inversions) through meiosis, despite classical expectations of sterility, reveals unexpected tolerance in human reproduction, allowing occasional transmission of such large rearrangements, albeit with increased risk of pregnancy loss or infertility, thereby maintaining them at low population frequencies^64–66^. Similarly, individuals with extreme centromeric size asymmetries between homologous centromeres raises critical questions about threshold effects in chromosomal segregation errors that may contribute to age-related aneuploidy and trisomy disorder risk in offspring^21,67^. The observation of pronounced homologous centromere size outliers underscores the need for future studies to determine whether such extreme variation influences centromere function, chromosome segregation fidelity, or reproductive outcomes. With the increasing recognition of centromeric variation’s role in disease, incorporating centromeric profiles into personalized medicine alongside conventional genetic risk factors holds substantial promise.

## Acknowledgements

The authors thank Jing Liu (Zhejiang University), Sirui Ye (Zhejiang University) and Lingjuan Xie (Shanghai Academy of Agricultural Science) for assistance in data analysis. We thank Dan Yang, Sanhua Fang and Yingying Huang from the Core Facilities, Zhejiang University School of Medicine for their technical support. We thank all APG members for suggestions. This study was funded, in part, by the National Key R&D Program of China (2025YFC3410300) to D.W., Xiaofei Yang, Y.M., B.S., Y.H. and Xiangyu Yang, National Key R&D Program of China (2024YFA1802500), the funding from International Institutes of Medicine of Zhejiang University (KY2022-098), Basic Research Center Program (32388102), the New Cornerstone Science Foundation through the XPLORER PRIZE and the K.C.Wong Education Foundation to G.Z., the National Natural Science Foundation of China (32422019), the Natural Science Foundation of Shaanxi Province (2024JC-JCQN-28), the National Key R&D Program of China (2022YFC3400300), the Fundamental Research Funds for the Central Universities (xzy012024088), and the Scientific Research Program of Shaanxi Provincial Department of Education (23JK0290) to Xiaofei Yang.

## Author contributions

G.Z., D.W. and Xiaofei Yang conceived and designed this study. G.Z. and K.Y. supervised this study. D.W., Y.S., Q.C. and Q.N. assembled and validated the centromere sequences. Y.S., K.F. and Q.C. annotated centromeric TEs and genes. Y.S. performed satellite clustering and HOR inference. S.W., Xiaofei Yang and Y.S. analyzed centromere synteny. D.W., F.Z. and Y.S. investigated centromeric inversions. D.Y., Xiangyu Yang, Y.Y. and Y.H. constructed iPSCs and conducted their culture. D.Y. and Y.Y. performed CENP-A CUT&Tag experiments and sequencing. L.N. analyzed the CUT&Tag and PacBio HiFi methylation data. Y.S., A.L. and S.W. analyzed centromere haplotypes. Y.M., K.Y. and B.S. provided suggestions on data analysis. Y.S., D.W., S.W., L.N., G.Z. and Xiaofei Yang wrote the manuscript, with input from all authors.

## Methods

### Assembly of human centromere sequences

Haplotype-level phased assemblies from 160 individuals in APGp1 were constructed using an integrated strategy based on high-depth PacBio HiFi and Oxford Nanopore Technologies (ONT) ultra-long reads, assembled via hifiasm^70^ and Verkko^71^ in both trio and Hi-C modes^24^. To comprehensively capture human centromeric diversity across global populations, we additionally included the phased, diploid genome assemblies of four published reference-level genomes T2T-CHM13^8^, T2T-CN1^11^, Q100-HG002 ^14,15^(https://github.com/marbl/HG002), and YAO^72^, and assemblies from HPRCy1^25^ (*n* = 43, excluding HG002 and individuals overlapping with HGSVC3) and HGSVC3^17^ (*n* = 65). For the assemblies from HPRCy1 and HGSVC3, contigs were scaffolded onto chromosomes using RagTag^73^.

### Evaluating centromere assembly quality

Based on alignments to the CHM13 pericentromeric regions (±5 Mbp), we first delineated the approximate boundaries of each centromeric region. We then validated centromere assemblies from APGp1 using the following complementary approaches. First, PacBio HiFi reads and Oxford Nanopore Technology (ONT) reads are phased with parental specific *K-*mers from NGS reads by secphase and Canu, respectively. These phased reads were subsequently aligned to corresponding haplotype-resolved assembly using minimap2^74^ (v2.26-r1175) and Winnowmap2^75^ (v2.03). Read mapping coverage uniformity across centromeric regions is evaluated by using GCI^26^ and NucFreq^27^. Depth profiles from GCI were generated with plot_depth.py (https://github.com/yeeus/GCI/tree/main/utility/plot_depth.py), and NucFreq profiles were visualized using NucPlot.py (https://github.com/mrvollger/NucFreq) after further filtration with “*samtools-F 2308*”. GCI scores for centromeric regions were computed using GCIscore.py. Candidate assembly issues reported by GCI were intersected using BEDTools^76^ (v2.31.1) with zero-depth regions derived from stringently filtered PacBio HiFi and ONT alignments. For NucFreq, we defined potential error regions as high-heterozygosity loci with at least 10% support for the second most frequent allele. Second, we applied the Flagger^25^ (v0.3.2) pipeline to detect potentially problematic regions (erroneous, duplicated, or collapsed). These regions were intersected with centromeric coordinates using BEDTools(v2.31.1) ^76^. Third, we assessed concordance between the assemblies and PacBio HiFi reads using VerityMap^29^ (v2.1.2), which identifies discordant *K*-mers and flags them for correction. Similarly, assembly concordance with ONT data was evaluated using GAVISUNK^28^, which detects concordant single-unique nucleotide *K*-mers (SUNKs) between the assembly and the long reads. Final assembly issue regions supported by at least two validation methods were merged using *bedtools multiinter*. For certain downstream analyses, gapless centromere sequences with no candidate issues were recruited. Additionally, we used Merqury^77^ (v1.4.1) to calculate assembly quality value (QV) in centromeric regions under default settings, based on a hybrid *K*-mer database (*k* = 21) constructed from both whole-genome NGS and PacBio HiFi reads.

### Annotating centromere satellite arrays

Human peri/centromeric satellite sequences, including αSat, HSat1, HSat2, HSat3, βSat, and γSat, were annotated separately. For αSat annotation, a consensus database was generated for each monomeric class from the T2T-CHM13 genome assembly. Identification of αSat elements was performed using megablastn with the parameter of ‘*-evalue 1e-10-task megablast*’. The resulting BED file was processed using a custom script (https://github.com/Asian-Pan-Genome/Centromere), with filtering criteria applied to retain hits with sequence identity ≥ 85% and alignment length > 100 bp. HSat2 and HSat3 annotations were conducted following the pipeline established for the T2T-CHM13 genome (https://github.com/altemose/chm13_hsat). Repeat elements were initially identified using RepeatMasker (v4.1.2) with the Dfam database (v3.3) under the following settings: sensitive mode (*-s*), species tag for human (*-species human*), and the NCBI BLAST-derived search engine RMBlast (*-e ncbi*). HSat1, βSat, and γSat regions were determined from RepeatMasker track using the following classification scheme: all “SAR” annotations were designated as HSat1A, “HSATI” annotations as HSat1B, “BSAT|Beta|LSAU-BSAT” as βSat, and “GSAT” annotations as γSat. Centromeric satellite annotation tracks, including those for ribosomal DNA (rDNA), were merged using satellite-specific distance thresholds: 10 Kbp for αSat and 5 Kbp for HSat1, HSat2, HSat3, βSat, and γSat. To precisely define pericentromeric and centromeric regions, an iterative extension approach was implemented using a custom script (https://github.com/Asian-Pan-Genome/Centromere). Specifically, initial start and end positions were defined based on the αSat region, with 5-Mbp flanking intervals on both sides. In each iteration, continuous satellite blocks were compared with the current pericentromeric region. If a block exceeded 10 Kbp in length or was located within 2 Mbp of the current pericentromeric boundaries, the region was extended to encompass this block. Acrocentric short arms were fully incorporated into the final satellite annotation track. To mitigate errors arising from excessive boundary extension in centromere length comparisons across populations, bar plots were generated for each centromere to facilitate manual validation. To determine the total length of each satellite array per chromosome, we first intersected the satellite annotations with the defined centromeric regions. The cumulative length was calculated as the sum of these merged intervals. To ensure consistency, all assemblies used in this study, comprising APGp1, HPRCy1, HGSVC3, and the external reference-quality genomes (T2T-CHM13, T2T-CN1, Q100-HG002, and YAO), were uniformly processed through the same annotation pipeline.

### Structural arrangements of human satellite arrays

#### Sequence strand switch

We identified and cataloged sequence strand switch points, classifying them based on genomic context: (i) within individual satellite arrays, (ii) between distinct satellite arrays, or (iii) at the boundaries between satellite arrays and the centromeric transition regions (regions of non-satellite DNA occurring near or between satellite DNA arrays in centromeres). Putative switch intervals within active arrays were initially identified using BEDTools (v2.31.1) and subsequently manually verified.

#### Centromere architecture classification

Centromere architecture (CenArch) was defined as the ordered sequence of satellite arrays along the centromeric region of each chromosome. To capture both fine-scale and coarse-scale architectures, satellite arrays were first filtered by size at 10 Kbp thresholds, and only the arrays passing each threshold were retained for ordering. For every haplotype, the retained arrays were then arranged according to their genomic coordinates, and the resulting ordered sequence of array types was used to assign an architecture label to each chromosome. The two sets of labels were subsequently cross-compared and manually curated to resolve threshold-induced splits or merges, yielding a consensus catalogue of 180 centromere architectures. For acrocentric chromosomes (13, 14, 15, 21 and 22), haplotypes lacking rDNA, exhibiting rDNA at the leading edge of the centromere, or not anchored to a telomere were additionally excluded to mitigate subtelomeric assembly artifacts. The resulting organizational profiles were visualized as bar plots for each centromere and manually validated. Large-scale duplications and inversions were called using self-alignment and pairwise dot plots generated with GEPARD^78^(v2.1.0). For CEN04, a word length matching the canonical HOR length (3,429 bp) was used. For CEN01, a 500-bp word length was selected to optimally resolve inversion boundaries.

### Centromeric inversions

#### Centromeric inversion calling

Peri/centromeric regions were defined as the centromeres and their 5-Mbp flanking sequences on both chromosome arms in T2T-CHM13. For each haplotype assembly, raw peri/centromeric inversion calls were generated against T2T-CHM13 using PAV^79^ (v2.4.6) and LSGvar (v1.0.0, https://github.com/Hanjunmin/LSGvar). Non-redundant peri/centromeric inversion calls were derived by integrating results from the two callers using BEDTools^76^ ‘intersect’ (v2.31.1) with the parameter: ‘*-f 0.5-r*’. All peri/centromeric inversions were manually curated by visualizing dotplots generated by mummerplot^80^ (v3.5). Given the low confidence of alignments between satellite arrays, the inversion calls where both flanking breakpoints localized to a single satellite array were not included in downstream analysis (e.g., the 27.6-Mbp HSat3B array in CEN09).

#### Inversion validation

Local alignments of long reads were manually inspected using NucFreq, GCI and GAVISUNK. For the ultra-large pericentric inversions in CEN07 and CEN09, Hi-C chromosomal interaction maps were used for confirming the inverted genomic structures. Clean Hi-C reads from C037-CHA-S17-01 (heterozygous inversion carrier) and C040-CHA-S20-01 (reference genotype) were mapped to the T2T-CHM13 assembly using BWA mem^81^, with primary alignments retained. Long-range interactions (>1 Mbp) were visualized as heatmaps and compared around inversion breakpoints, where strong pericentric interactions indicated the mapping of paired short reads to the two pericentric inversion breakpoints of CEN07. Hi-C data for the CEN09 pericentric inversion (C051-CHA-N11-01, heterozygous inversion carrier; C045-CHA-N05-01, reference genotype) were processed following the same workflow.

As supplementary validation, Bionano optical maps were generated for C037-CHA-S17-01 and C051-CHA-N11-01. High-molecular-weight genomic DNA was isolated from frozen whole blood using the Nanobind CBB Big DNA Kit. DNA was labeled and counterstained via the Direct Label and Stain (DLS) protocol, loaded onto a Saphyr chip and imaged on the Saphyr instrument (Bionano Genomics). For each sample, an in silico reference map was generated from its corresponding diploid genome assembly using the fa2cmap_multi_color.pl script (Bionano HybridScaffold v1.0). This script performed an in silico digestion of the DLE-1 labeling motif (CTTAAG) to produce a reference CMAP file for each assembly. Subsequently, the align_bnx_to_cmap.py script (Bionano Pipeline v1.0) was used to align raw molecules to their respective in silico reference CMAPs. This alignment process utilized the Bionano RefAligner (v1.0) with a Signal-to-Noise Ratio (SNR) filter of 2 (--snrFilter 2) and optimized parameters for DLE-1 labelin on the Saphyr platform (optArguments_nonhaplotype_DLE1_saphyr_human.xml). Resulting alignments were loaded into Bionano Access for manual inspection and visualization via the “Anchor to Molecules” alignment view.

### αSat monomer clustering and HOR inference

#### Global monomer clustering algorithm

To address potential reference bias in HOR identification across multi-ancestry genomes, which will lead to the missing identification of population or individual specific monomer and HOR structures, we implemented a novel workflow for global monomer clustering (see details in **Supplementary Note 2**). We extracted ∼244 million monomers from assemblies in APGp1, HPRCy1, HGSVC3, and four reference-level complete assemblies (T2T-CHM13, T2T-CN1, Q100-HG002 and YAO). Similar monomer sequences were identified at a 96% identity threshold and clustered by abundance using VSEARCH^82^ (v2.30.0), a rapid heuristic-based tool for sequence analysis. Due to the greedy nature of the clustering algorithm, we further refined clusters by applying the Louvain method for community detection. The accuracy of monomer clustering was evaluated by comparison with curated HOR annotations in the T2T-CHM13 assembly. Each resulting monomer cluster was assigned a unique numeric ID. For subsequent HOR inference, monomer sequences from each haplotype assembly were merged into continuous blocks (maximum interval ≤ 10 Kbp) using BEDTools (v2.31.1) and encoded as strings of cluster IDs.

#### *De novo* HOR inference

We employed two complementary approaches for HOR identification: the graph-based HORmon^44^and the hierarchical tandem repeat mining (HTRM)-based HiCAT^45^, with details provided in **Supplementary Note 2**. To ensure robust HOR calls, distinct filtering criteria were applied to the outputs of HORmon and HiCAT, reflecting their different methodological principles. This yielded a final set of HOR structural variants (StVs), which were categorized into five classes based on population frequency: core (>90%), dispensable (>5%), low-frequency (>1%), rare (1-1%), and singleton (*n* = 1). To group HOR StVs into structurally homogeneous arrays, we clustered them based on the edit distance between their monomer ID strings. This approach quantifies organizational differences, including insertions, deletions, and substitutions of monomers, the grouping of StVs with highly similar monomer compositions. The resulting clusters represent distinct HOR arrays and were further rigorously evaluated against those previously reported in the T2T-CHM13 assembly.

#### HOR inter-chromosome sharing

Pangenomic profiling of HOR StVs enabled population-level analysis of inter-chromosomal sharing patterns. An HOR StV was defined as “shared” if it occurred on more than two chromosomes. A shared HOR array was designated as any array containing at least one shared StV. To ensure reliability, sharing events located within regions with potential assembly artifacts or incomplete centromeres were excluded. For CEN01/05/19/16, CEN13/21, and CEN14/22 groups, we identified the longest shared segments that contained consistent sequence information among dominant shared HOR StVs to infer their evolutionary history. According to the layered expansion model^9^, the same StV may display differential sequence divergence across layers, with core layers being more homogeneous and flanking layers more divergent. To perform a spatially unbiased sampling, we extracted shared segment sequences from three regions of the active arrays: Region 1 (p-arm flanking regions), covering 0-200 Kbp from the active array start; Region 2 (central regions), spanning the midpoint ± 100 Kbp; and Region 3 (q-arm flanking regions), comprising the terminal 200 Kbp. For chromosome 19, we adjusted the region length to 100 Kbp (**Supplementary Note 3**). From each region, we randomly sampled 1,000 shared sequences per StV. The pooled sequences were aligned using MAFFT^83^ (v7.525), and used to construct maximum-likelihood phylogenies with IQ-TREE^84^ (v2.3.6) with the parameter setting:-m MFP-B 1000 --bnni-T AUTO. This procedure was repeated ten times to assess the robustness of tree topology. Furthermore, we tested the assumptions that ancestral StVs accumulate greater intra-variant polymorphism than recently derived ones, and that closely related StVs share more recent common ancestry and thus exhibit lower sequence divergences. We extracted all shared segment sequences labeled by their derived StVs and chromosomes. After removing duplicates with VSEARCH^82^ (v2.30.0), sequences were aligned with MAFFT^83^. We then quantify pairwise allele mismatches both within and between StVs to characterize sequence divergence.

### Centromere haplotyping and population stratification

#### Pairwise collinearity-aware alignment

We employed HORSCAN (https://github.com/XDwan/HORSCAN), an innovative tool based on multi-level dynamic programming, that enables pairwise, single-satellite-aware collinearity analysis for αSat arrays by integrating satellite clustering categories and HOR structures. This HOR structure-guided alignment principle allows HORSCAN to accurately identify both monomer-level sequence variations (monomer CIGAR) and HOR-level structural differences.

#### Centromere Structural Distance

To quantitatively assess the structurally evolutionary distance between αSat arrays, we designed a two-component composite metric termed the Synteny Distance. This metric is based on the monomer-level alignment output from HORSCAN and is designed to decouple and integrate two distinct modes of divergence: sequence divergence (at the monomer level) and annotation divergence (at the HOR structural level).

The complete computational formula is:

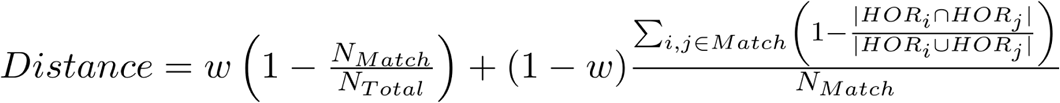

The first component quantifies the monomer sequence divergence rate. This term measures the proportion of non-matching alignments (i. e. mismatches and indels) relative to the total number of aligned monomer pairs. It reflects the raw evolutionary distance at the fundamental DNA sequence (monomer) level, with a high value indicating significant sequence-level differentiation between the two arrays. The second component specifically quantifies structural annotation divergence among sequence-conserved monomers. This term restricts its analysis to the set of sequence-identical monomer pairs. For each such matching pair, it retrieves their respective structural annotations (HOR*_i_* and HOR*_j_* which describe the HOR structures they belong to) and calculates the Jaccard distance between these annotation sets. This metric captures a sophisticated form of structural evolution: a high value signifies that even where monomer sequences are perfectly conserved, the higher-order organization (HOR-level role) of those monomers has diverged.

#### Centromere haplotyping

For each chromosome, we constructed an all-by-all pairwise synteny distance matrix between all haplotype assemblies for the centromeric region. After removing incomplete centromeres, hierarchical clustering was performed using the *pheatmap* R package (v1.0.13, https://github.com/raivokolde/pheatmap) with average linkage, and CenHap groups were defined by cutting the dendrogram at *k* = 2 to 10. On chromosomes displaying distinct hierarchical HOR organization, major CenHaps were assigned based on well-defined structural patterns, particularly the presence of haplotype-specific StVs forming discrete layers. This semi-automated classification was applied uniformly across all chromosomes. It should be noted that HOR annotation based on 96%-similarity monomer clustering lacks discriminative power in highly homologous regions. Since synteny distances in these cases are largely shaped by structural variation, such as alignment gaps and array sizes, rather than nucleotide-level diversity, the resulting CenHap groupings may not fully represent phylogenetic relationships. These classifications should be therefore treated with appropriate caution in downstream analyses. Sequence similarity heatmaps, reflecting local homogenization, were produced with ModDotPlot^68^ (v.0.8.5) under a static mode. The visualization was configured with a custom sequential color palette and non-uniform breakpoints for percent identity mapping to enhance visual discrimination of homology levels.

#### Alignment-free validation of CenHap classification using *k*-mer profiles

To independently evaluate CenHap assignments, we performed an alignment-free *k*-mer-based analysis using raw sequence composition profiles. 31-mer sequences were extracted directly from core αSat arrays across all assemblies without relying on monomer clustering, sequence alignment, or HOR-based structural variant annotations. To enrich population-informative sequence signals, ubiquitous *k*-mers and singleton *k*-mers were removed, retaining *k*-mers with cohort frequencies between 2% and 25%. Pairwise Bray–Curtis dissimilarity matrices were generated based on normalized k-mer abundance profiles, and sample relationships were visualized using Principal Coordinates Analysis (PCoA). The resulting *k*-mer-based clustering patterns were compared with independently defined CenHap classifications to assess the concordance between sequence-composition variation and CenHap assignments.

CenHap frequencies for each chromosome were compared across superpopulations. Population codes illustrated in Figs. 5d and 5h are defined as follows: ACB: African Caribbean in Barbados; ASW: Americans of African Ancestry in Southwest US; BEB: Bengali in Bangladesh; CLM: Colombian in Medellin, Colombia; CHA-C: Chinese Han in Central China; CHA-E: Chinese Han in Eastern China; CHN-N: Chinese from Northern China (Han, Manchu and Mongol); CHA-NE: Chinese Han from Northeastern China; CHN-NW: Chinese from Northwestern China (Hui, Kazak and Uyghur); CHA-S: Chinese Han from Southern China; CHN-SW: Chinese from Southwestern China (Bai, Bouyei, Miao, Tujia, Yi and Zhuang); CZA: Chinese Tibetan; ESN: Esan in Nigeria; EUR-NW: Utah residents (CEPH) with Northern and Western European Ancestry, British in England and Scotland; EUR-S: Iberian Population in Spain, Toscani in Italia; FIN: Finnish in Finland; GWD: Gambian in Western Division, The Gambia; IND: Gujarati Indian in Houston, TX, Indian Telugu in the UK; PJL: Punjabi in Lahore, Pakistan; JPT: Japanese in Tokyo, Japan; KHV: Kinh in Ho Chi Minh City, Vietnam; KOR: Korean in South Korea; LWK: Luhya in Webuye, Kenya; MSL: Mende in Sierra Leone; MXL: Mexican Ancestry in Los Angeles, California; PEL: Peruvian in Lima, Peru; PUR: Puerto Rican in Puerto Rico; STU: Sri Lankan Tamil in the UK; YRI: Yoruba in Ibadan, Nigeria.

#### Phylogenetic inference and CenHap divergence dating

To resolve the phylogenetic relationships and divergence times of major CenHaps on chromosomes 4, 5, 8, 10, and 17, we first evaluated the topological consistency between p-arm and q-arm phylogenies. Independent phylogenies were inferred using two datasets: (1) graph-based SNPs from highly linked regions, filtered with parameters “*--set-missing-var-ids @:# --geno 0.1 --maf 0.02 --snps-only*”, and (2) 20-Kbp flanking sequences extracted from unique alignment blocks in pericentromeric regions, identified by whole-chromosome alignment to T2T-CHM13 using MUMmer^85^ (v4.0.1). Multiple sequence alignments were generated with MAFFT^83^ (v7.525), and maximum-likelihood phylogenies were reconstructed using IQ-TREE^84^ with an auto-detected model in 1000 bootstrap replicates (v2.3.6). For divergence time estimation, two representative haplotypes were selected from each major phylogenetic lineage, with chimpanzee and bonobo as outgroups^44^. Node ages were estimated under a molecular clock model using BEAST^86^ (v2.7.7), applied to branch lengths from the 20-Kbp regions, using human-chimpanzee and chimpanzee-bonobo divergence times of 6.2 and 1.6 million years, respectively. For each alignment block, independent MCMC runs were performed and convergence was assessed by inspecting trace plots and effective sample sizes. After discarding 10% burn-in, posterior tree samples from independent runs were combined and summarized using TreeAnnotator in BEAST (v2.7.7) suit. The maximum clade credibility tree was selected as the target tree, and node ages were summarized as posterior medians with 95% highest posterior density intervals. Nodes with posterior probabilities below 0.5 were not annotated. Node ages are reported as posterior median estimates, and uncertainty is represented by 95% Highest Posterior Density (HPD) intervals.

### Epigenetic analysis of human centromeres

#### PBMC isolation and iPSC reprogramming

Peripheral blood mononuclear cells (PBMCs) were isolated from anticoagulated whole blood samples of five individuals (C014-CHA-E14-01, C020-CHA-E20-01, C050-CHA-N10-01, C080-CHA-NE20-01 and C102-CZA02-01) using Ficoll density-gradient centrifugation following the manufacturer’s protocol (Sanguine Biosciences). Diluted blood was layered onto Ficoll and centrifuged at 1,000 × g for 30 minutes with the brake off. The mononuclear layer was collected, washed twice with PBS, and immediately used for reprogramming. PBMCs were reprogrammed using the Epi5™ Episomal iPSC Reprogramming Kit (Thermo Fisher Scientific). Cells were electroporated with episomal plasmids on the Neon™ Transfection System using the program: 1550 V, 10 ms, three pulses. After electroporation, cells were maintained in N2B27 medium supplemented with bFGF, then transitioned to Essential 8™ Medium according to the manufacturer’s instructions. iPSC colonies typically emerged between days 15 and 21.

#### CUT&Tag library preparation and sequencing

Chromatin profiling was performed using the Hyperactive Universal CUT&Tag Assay Kit for Illumina Pro (Vazyme TD904) according to the manufacturer’s instructions. Approximately 10,000 iPSCs were harvested per reaction and gently bound to Concanavalin A–coated magnetic beads (ConA Beads Pro) to immobilize intact nuclei. Cells were permeabilized with 0.05% digitonin and incubated with a mouse monoclonal anti-CENP-A antibody (Enzo Life Sciences, ADI-KAM-CC006-E) at 4°C overnight, followed by a 1-hour incubation with secondary antibody at room temperature. The pA/G–Tn5 transposome was then added to mediate tethered tagmentation near antibody-bound chromatin regions for 1 hour at 37°C in the presence of magnesium ions. Following enzymatic fragmentation, DNA was released using DNA Extract Beads Pro and purified with VAHTS DNA Clean Beads. Indexed sequencing libraries were generated using the TruePrep Index Kit V4 (Vazyme TD204–TD207) by PCR amplification for 12-14 cycles, ensuring sufficient yield while avoiding over-amplification. Library size distribution (∼200–800 bp) and concentration were verified by Agilent 2100 Bioanalyzer and Qubit dsDNA HS Assay, respectively. All qualified libraries (20 μL final volume per sample) were pooled equimolarly and sequenced on Illumina Novaseq 6000 platform (paired-end 150 bp). IgG control libraries were prepared in parallel to evaluate background signals and assess antibody specificity.

#### CUT&Tag data analysis

CUT&Tag sequencing data from the five PBMC-derived iPSC samples were analyzed. Low-quality bases and adaptors were firstly removed via fastp^87^ (v0.23.4). Subsequently, the filtered reads were aligned to the corresponding diploid assembly of each sample via BWA mem^81^ (v0.7.18). Only primary alignments were retained with “*samtools-F 2308*”. Finally, normalized CENP-A enrichment was obtained using bamcompare from deepTools^88^ (v3.5.6) with the command of “bamCompare-b1 {chip.bam}-b2 {igG.bam}-o {log2ratio.bedgraph} --binSize 1000 --numberOfProcessors 40 --operation ratio --outFileFormat bedgraph”. To soften potential mapping biases in highly repetitive centromeric regions, we applied a unique-*k*-mer-assisted mapping strategy. Briefly, we used meryl(v1.4.1) to identify unique 21-mers and obtain their genomic positions on the diploid assembly. We then retained only CENP-A and IgG CUT&Tag reads that overlapped these unique 21-mer positions using BEDTools ‘intersect’ (v2.31.1). Finally, we re-calculated the CENP-A enrichment using bamcompare from deepTools (v3.5.6) on the filtered BAM files, following the same parameter as described above.

#### CDR analysis

5mC methylation signals were called from the mapped PacBio HiFi reads with 5mC base modification values using pb-CpG-tools (v1.4.0) (https://github.com/PacificBiosciences/pb-CpG-tools). CDR-Finder^89^(https://github.com/arozanski97/CDR-Finder) was used to identify draft CDR regions with the parameters of window_size = 5000 and len_filter = 20000 and the regions lacking obvious hypomethylation were removed in manual curation. Filtered adjacent CDR regions were merged if they met the following criteria: (1) interval length less than 100 Kbp; (2) both CDR regions length larger than 0.8 time of their interval; (3) summed lengths of the two CDRs larger than 0.6 time of the merged region. We compared the genomic locations of CDRs and CENP-A enrichment peaks to quantify the epigenetic consistency. For each CDR identified in the five samples, we considered it to be consistent with CENP-A peaks if the overlapping region between the two intervals covered more than 50% of the length of either the CDR or the peak. And a centromere was considered epigenetically consistent only if all of its CDRs are consistent with CENP-A peaks.

The associations between mean CDR length and chromosome size or HOR array size were evaluated using Spearman’s rank correlation. For each chromosome, the mean CDR length was calculated and correlated with the corresponding whole chromosome size or HOR array size. Spearman’s correlation coefficients and associated *P* values were calculated using the stat_cor() function (method = “spearman”) implemented in the ggpubr (v0.6.0) package in R (v4.5.1).

To lift CDRs to a specific reference, HORSCAN results were converted to PAF format using a custom script. The PAF file was then transferred to a chain file using paf2chain (v0.1.0, https://zenodo.org/records/8108447). Finally, CrossMap^90^ (v0.2.8) was used to lift CDR regions to the corresponding reference via the chain file.

StainedGlass^91^ was used to generate dot plots and calculate pairwise identity across fixed 5 Kbp windows. To avoid the artificially reduced identity estimates for similar sequences as previously reported^68^, we adjusted the window size to 4.9 Kbp and 10 Kbp for chromosomes 4 and Y, respectively. The similarity for each window is calculated as the mean identity of the two adjacent windows on both sides. Furthermore, the CDR and non-CDR active HOR regions identity was represented by the mean of the similarity of the windows inside them.

#### Comparing CDRs in APGp1 and HGSVC3

To investigate whether CDR location preferences vary across populations, we selected 20 samples from HGSVC3 and 10 samples from HPRCy1 for CDR profiling. ONT reads were aligned using Winnowmap2^75^ (v2.03), and methylation frequencies were calculated with modkit (https://github.com/nanoporetech/modkit). CDR annotation and lift were performed using the same pipeline as described above.

#### Centromeric recombination

To characterize genetic linkages within and between pericentromeric regions of p and q arms, we conducted correlation and linkage disequilibrium (LD) analyses using a pangenome graph-decomposed SNP dataset^24^. For correlation analysis, pericentromeric intervals spanning 2 Mbp from the centromere on autosomes and 5 Mbp on chromosome X were partitioned into non-overlapping 100-Kbp windows, with exclusion of windows overlapping pericentromeric satellite arrays (γsat, βsat, HSat1-3) or complex inversion loci. SNP filtering was performed using PLINK^92^ (v1.9.0-b.7.7)using the parameter: “*--set-missing-var-ids @:# --geno 0.1 --maf 0.02 --snps-only*”. A modified filtering strategy was applied to acrocentrics, CEN01, CEN09, CEN16 and CENX: “*--set-missing-var-ids @:# --maf 0.02 --snps-only*”. Only windows retaining ≥50 SNPs after filtering were included in downstream analyses. Pairwise identity-by-state (IBS) values between haplotype genomes within each window were calculated using PLINK^92^ (v1.9.0-b.7.7). Correlation coefficients (*r*) of IBS profiles among different windows (from a single arm or two arms) were used to generate the final correlation heatmap, spanning two pericentromeres from p and q arms (**Fig. 7a**). For LD analysis, haplotype SNP data were converted into diploid format, with the chromosome X treated as homozygous, and LD values were calculated across the same extended pericentromeric intervals using PLINK^92^.

To explore the genetic association between pericentromeric sequences from two arms and centromere CenHaps, we measured pairwise CenHap distances based on synteny scores of αSat HOR arrays. We then calculated the correlation between the pericentromeric genetic distances (1 - IBS) of each pericentromeric window and centromere CenHap distances to characterize the pericentromere-centromere genetic linkage (**Fig. 7a**).

Inter-flank phylogenetic concordance. For each chromosome, p-arm and q-arm centromere-flanking sequences were independently aligned using MAFFT (v7.525), and maximum-likelihood phylogenies were reconstructed with IQ-TREE (v2.3.6) using the parameters-m MFP-B 1000 --bnni-T AUTO. Specifically, to reduce potential biases arising from differences in p-arm and q-arm linkage length, SNP availability, and repetitive sequence composition, we restricted phylogenetic reconstruction to non-satellite, non-tandem-repeat pericentromeric regions. Candidate regions were selected to maximize phylogenetically informative SNP content while ensuring comparable sequence representation between the two centromere flanks. The final pericentromeric phylogeny for each chromosome is available through the APG portal (https://genome.zju.edu.cn/APG/download/). Concordance between the two phylogenies was quantified using three complementary metrics: normalized Mutual Clustering Information (MCI), which measures global topological similarity while reducing sensitivity to low-support branches; quartet agreement calculated after collapsing branches with bootstrap support <70%, which evaluates local topological consistency; and the Spearman correlation between pairwise identity-by-state (IBS) distance matrices, with statistical significance assessed by a Mantel permutation test (10,000 permutations). Chromosome Y, which lacks meiotic recombination and is therefore expected to share a single genealogy across both flanking regions, served as a positive control for maximal inter-flank concordance. Chromosomes with concordance values below the genome-wide mean for all three metrics were classified as exhibiting inter-arm phylogenetic discordance.

Recombination hotspots were analyzed as previously described, following the pipeline established for investigating recombination between human acrocentric chromosomes^10^. Briefly, each haplotype assembly was scanned for 17 known human PRDM9 binding motifs^58^ using FIMO^93^ with the command “*fimo --verbosity 2 --thresh 1e-4 --oc ${dir} PRDM9_motifs.human.txt ${seq}*”. To minimize false discoveries, only motif matches with a *P* value < 0.3 were kept. For each PRDM9 motif, occurrence counts were quantified in 20-Kbp windows across each haplotype chromosome using BEDTools (v2.31.1).

#### Comparing centromeric and pericentromeric single-base substitution rates

Given the frequent recurrence of large SVs in centromere regions, a high-quality curated centromere panel that rules out false-positive orthology arising from SVs and conversion is essential for single-base substitution analyses. Among all the gapless centromere pairs from assemblies in APGp1, HGSVC3 and HPRCy1, we firstly excluded the centromere pairs with a HORSCAN (v1.0) alignment score less than 0.9, and subsequently a rigorous masking strategy was applied to mitigate the paralogous influence from repeat expansion, contraction, and gene conversion (See **Supplementary Note 4**). Specifically, we masked αSat positions harboring HOR structural variants (detected by HORSCAN) or periodic variation arising from gene conversion, together with their 10 flanking monomers on each side, to prevent these mutations from perturbing the alignment of the entire HOR periodic unit. Subsequently, centromere pairs with >20% of their total αSat array length masked were excluded. The remaining pairs underwent manual curation to exclude centromere pairs exhibiting dispersed structural variants or large-scale segmental insertions or deletions (**Supplementary Note 4**). We further employed phylogeny (inferred by IQ-TREE) to ensure the authentic orthology of monomer alignments between centromeres, by removing monomer pairs with abnormal counts of substitutions (edit distance > 5). Finally, 93 structurally and sequentially orthologous centromere pairs were selected for single-base substitution analysis (**Supplementary Table 20**).

Extended pericentromeric regions (10 Mbp flanking each side of the αSat array) of each pair were aligned in 50 Kbp non-overlapping windows using Winnowmap2 (v2.03). Windows that were non-syntenic or exhibited outlier local SNP density (>3 standard deviations above the chromosomal mean) were excluded. Considering the recombination in pericentromeres, we manually curated the pericentromeric intervals of each pair for substitution analysis, by determining the boundaries of variation density leaps at each arm (**Supplementary Note 4**).

The substitution rate for each pair was calculated by dividing the total number of nucleotide mismatches, derived from HORSCAN results for centromere αSat array or CIGAR strings from Winnowmap2 alignments for pericentromeric intervals, by corresponding alignment lengths. Considering the challenge in estimating the divergence time of nearly identical sequences, we compare raw substitution rates between centromeric and pericentromeric regions, instead of estimating the inter-generation or yearly single-base substitution rates. For our paired comparison, we computed the log10-transformed ratio of the core centromere rate to its corresponding pericentromeric rate. The statistical significance of these ratios was assessed against the null hypothesis of a 1:1 ratio using a two-sided, non-parametric Wilcoxon signed-rank test. We further partitioned each αSat array based on the CDR and HOR annotations and calculated substitution rates separately for each subregion.

## Code availability

The codes and custom scripts used in this study are publicly accessible at https://github.com/Asian-Pan-Genome/Centromere. HORSCAN could be assessed at https://github.com/XDwan/HORSCAN.

## Data availability

Centromere annotations for each assembly have been deposited on Github (https://github.com/Asian-Pan-Genome/Centromere), including genomic coordinates of centromeric regions, various satellite arrays, and detailed HOR annotations, and can be browsed online in APG CentroViewer (https://genome.zju.edu.cn/APG/cenview/). High-resolution images of centromere assembly quality evaluation, pericentromeric phylogeny, satellite organization, CENP-A CUT&Tag mapping enrichment plots, and alignments of 93 centromere pairs for substitution-rate analysis are available at Zenodo (https://doi.org/10.5281/zenodo.17647353) or APG portal (https://genome.zju.edu.cn/APG/download/). CENP-A CUT&Tag sequencing data have been submitted to the National Genomic Data Center (NGDC, https://ngdc.cncb.ac.cn/) under HRA014419 through the experiment number HRX2355018 to HRX2355022, and are released with the approval by the Human Genetic Resources Administration of China (registration number 2026BAT00481). Individuals interested in accessing the data should submit a request to the DAC via NGDC.

