## Supplementary Figures for "Multidimensional variation and population stratification across 8000 complete human centromeres"

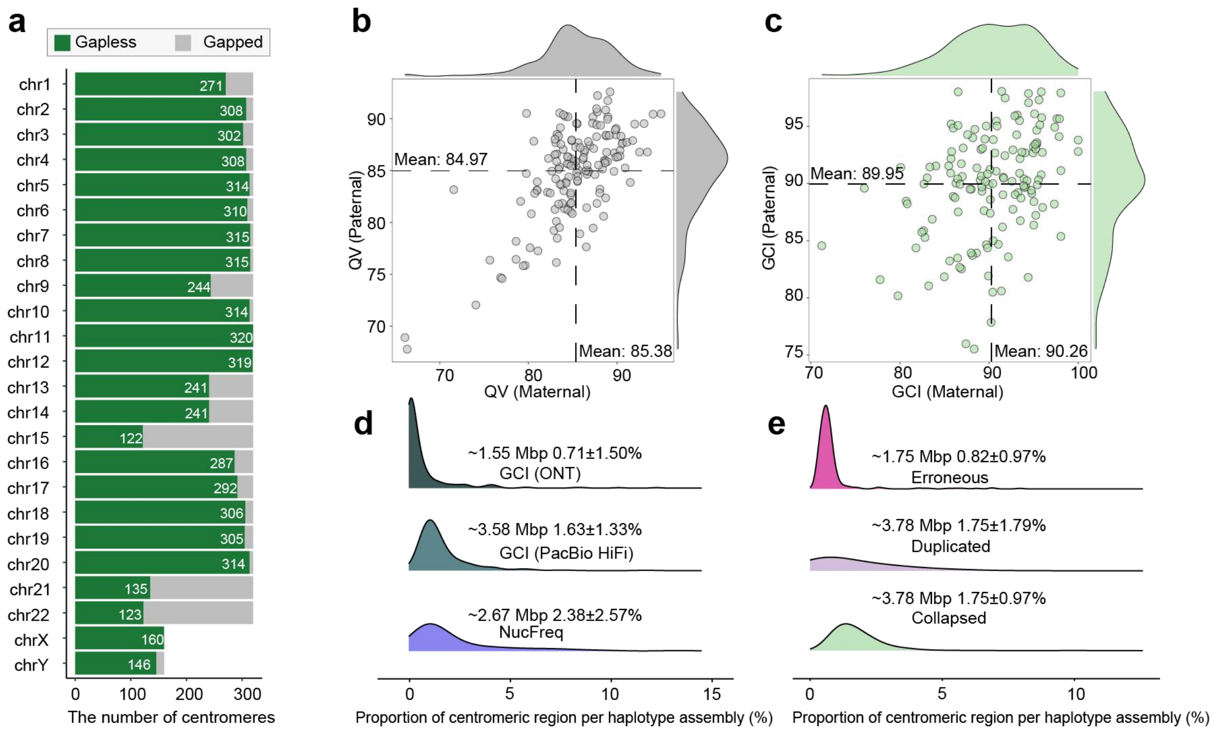


**Supplementary Fig. 1 | Quality assessment of centromere assemblies in APGp1. a**, Centromere assembly completeness per chromosome in APGp1 genomes. Green bars represent the haploid numbers with gap-free assembly across chromosomes. **b**, Quality Values (QVs) of APGp1 centromeres in maternal and paternal haploid assemblies. **c**, Genome Continuity Inspector (GCI) scores of APGp1 centromeres per haploid. **d**, Sizes and percentages of potential assembly issue regions reported by GCI, and Nucfreq evaluation based on PacBio HiFi and ONT read alignments on APGp1 centromeres per haploid. **e**, Sizes and percentages of potential assembly issue regions evaluated by Flagger on APGp1 centromeres per haploid.

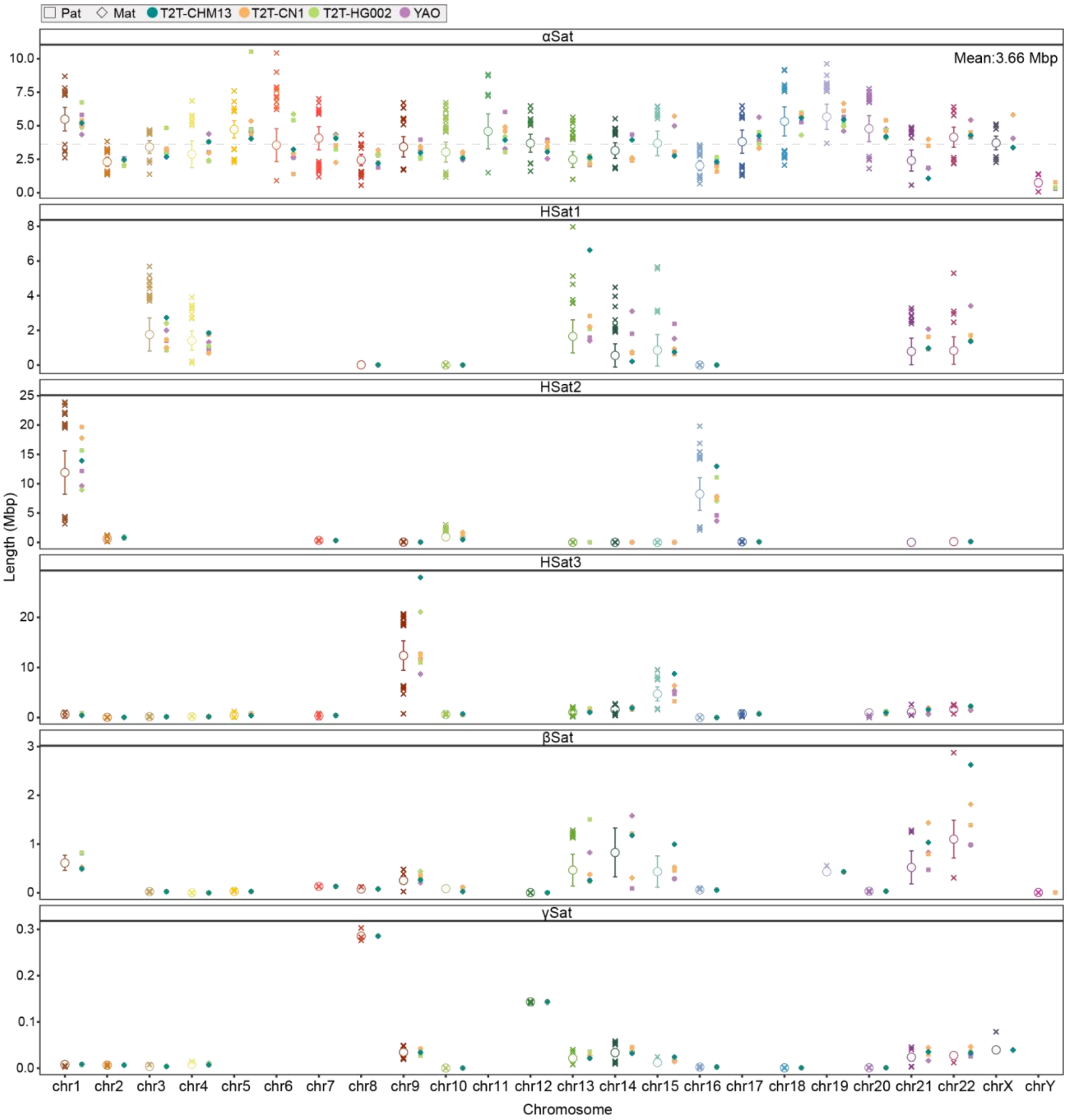


**Supplementary Fig. 2 | Composition and size variation of major centromeric satellite arrays across all chromosomes in APGp1.** Length distributions of αSat, HSat1, HSat2, HSat3, βSat and γSat arrays are plotted per chromosome (left), alongside previously released T2T-level reference assemblies (right), including phased assemblies of T2T-CN1, YAO and Q100-HG002, and haploid assembly T2T-CHM13. Hollow circles represent the mean values, and the bars indicate the mean ± standard deviation (s.d.). Outliers, defined as any values falling outside the range of mean ± 2 s.d. threshold, are denoted by "×". In the αSat panel, the dashed line indicates the mean α-satellite array size per centromere. Caution should be noted that for acrocentric chromosomes (13, 14, 15, 21 and 22), the distributions of satellite array lengths may be partially confounded by incomplete assembly within sub-telomeric region, which contain highly similar satellite arrays (e.g., HSat3, βSat, HSat1) that remain exceptionally challenging to fully resolve.

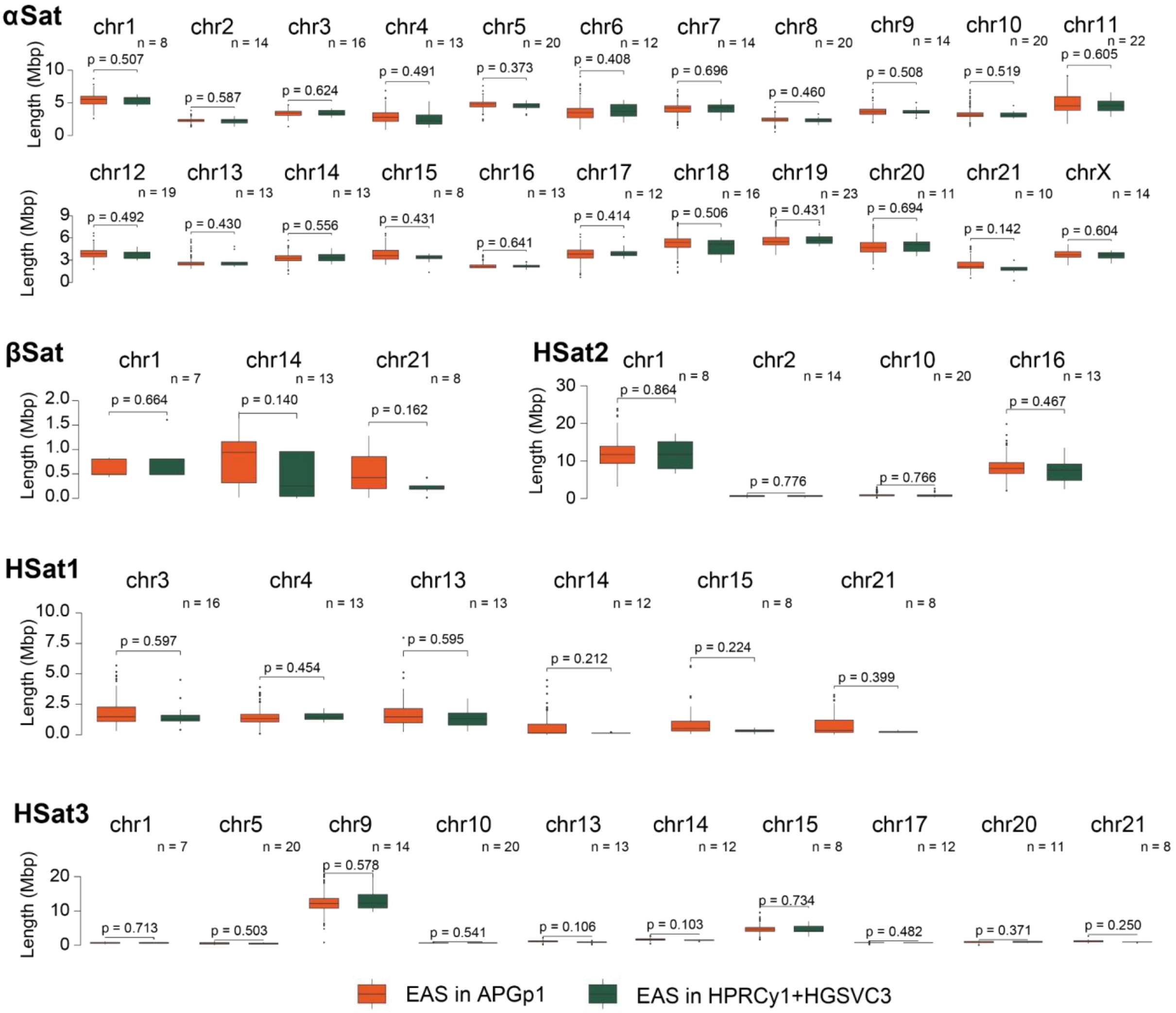


**Supplementary Fig. 3 | Batch effect evaluation on centromeric satellite arrays in the EAS superpopulation between APGp1 and previous projects (HPRCy1 and HGSVC3)**. Comparisons of αSat, HSat1, HSat2 and HSat3 array sizes between EAS individuals from APGp1(orange) and those from HPRCy1 and HGSVC3 (green) are shown. Only haplotypes with completely assembled centromeres on the corresponding chromosome are considered. For each satellite and chromosome, a two-sided Wilcoxon rank-sum test is performed whenever at least three non-APG haplotypes are available. The APG group is randomly down-sampled to match the non-APG size and the test is repeated 10 times; the mean *p*-value is reported above each box pair, and the number of non-APG haplotypes used in the comparison is indicated in the upper-right corner of each panel (*n*). Only chromosomes whose mean satellite array spans at least 500 Kbp in EAS are shown. No significant differences are observed, indicating the absence of technical batch effects. In each boxplot, the center line represents the median, the box limits indicate the upper and lower quartiles (25th and 75th percentiles), and the whiskers extend to 1.5 times the interquartile range from the box.

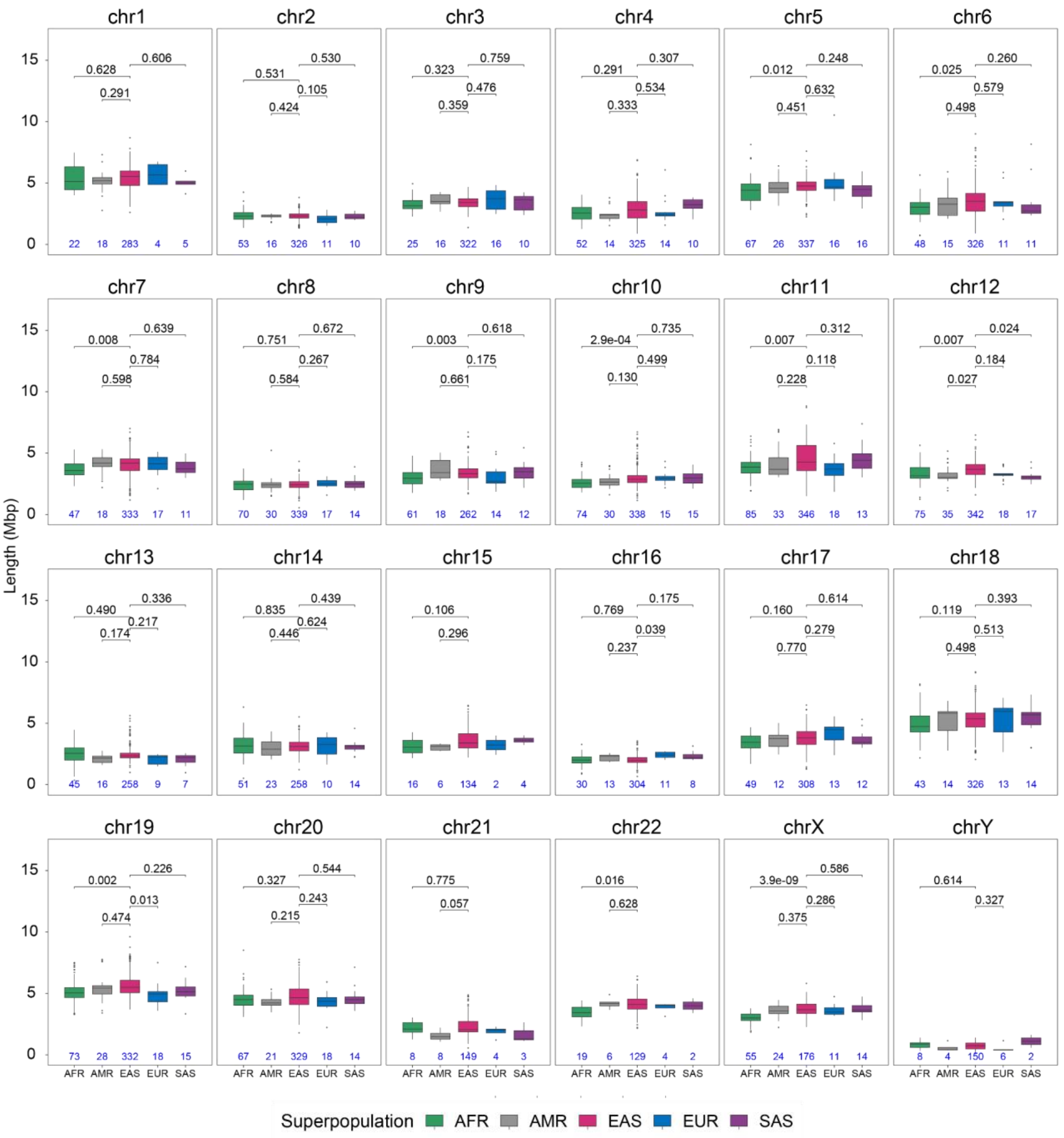


**Supplementary Fig. 4 | αSat array size variation across five global superpopulations.** Boxplots illustrate the size distribution of αSat arrays across five superpopulations. For each chromosome, a two-sided Wilcoxon rank-sum test was performed when at least five non-EAS haplotypes were available. To control for sample size asymmetry, the larger comparative group was randomly downsampled to match the smaller group, and the test was permuted across 10 iterations. The mean p value is reported above each box pair. Total numbers of fully assembled centromeres analyzed per chromosome are indicated at the bottom of each boxplot. In each boxplot, the center line represents the median, the box limits indicate the upper and lower quartiles (25th and 75th percentiles), and the whiskers extend to 1.5 times the interquartile range from the box. AFR, African ancestry; AMR, American ancestry; EAS, East Asian ancestry; EUR, European ancestry; SAS, South Asian ancestry.

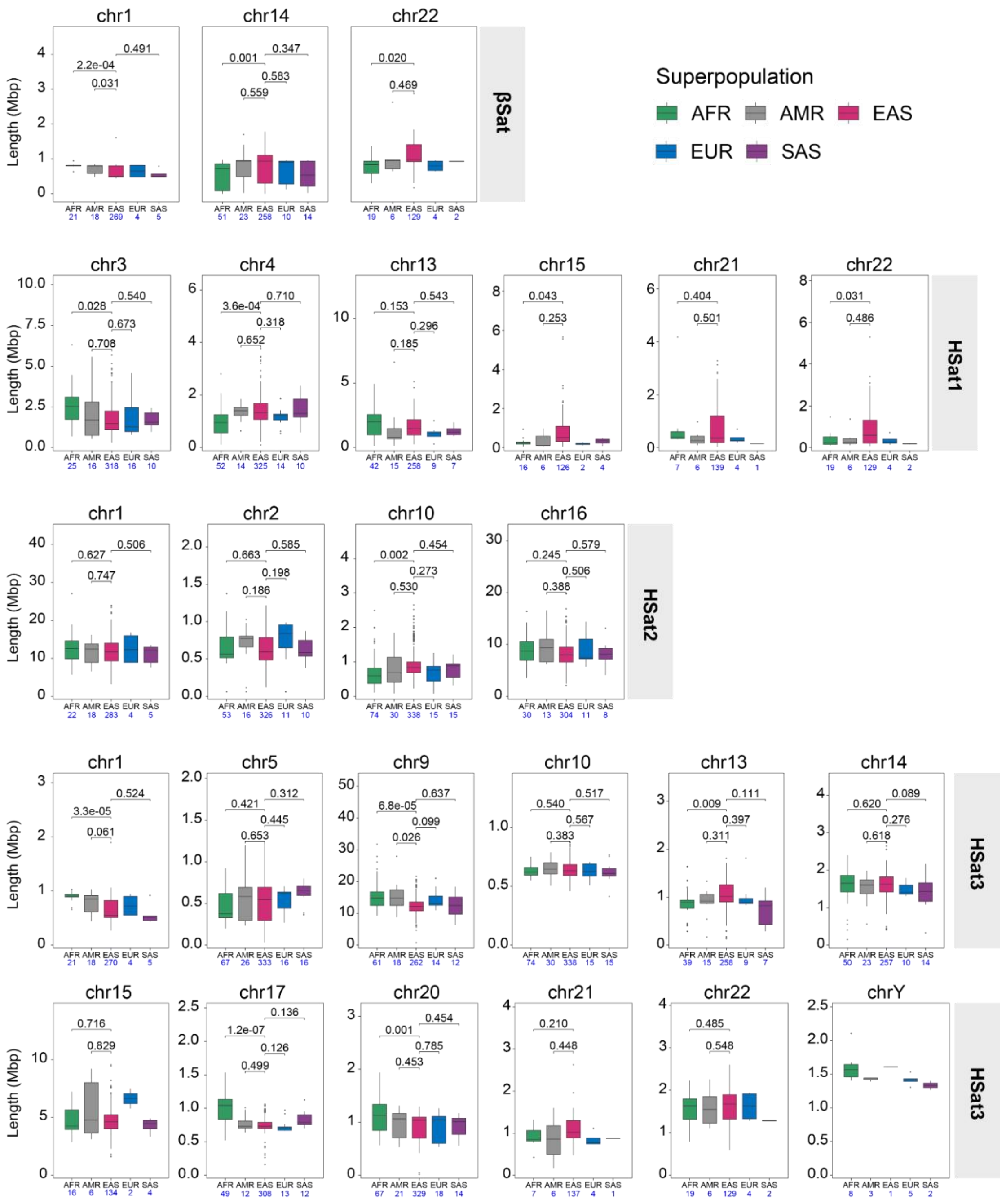

**Supplementary Fig. 5 | HSat and βSat array size variation across five superpopulations.** Boxplots illustrate the size distribution of βSat**,** HSat1, HSat2 and HSat3 arrays across superpopulations. For each chromosome, a two-sided Wilcoxon rank-sum test was performed when at least five non-EAS haplotypes were available. To eliminate sample size asymmetry, the larger group was randomly downsampled to match the smaller group, with the mean *p* value derived from 10 independent permutation runs. Comparisons involving acrocentric centromeres should be interpreted with caution, due to possible incomplete assembly near telomeric regions on these chromosomes. Total numbers of completely assembled centromeres analyzed per chromosome are indicated below each boxplot. In each boxplot, the center line represents the median, the box limits indicate the upper and lower quartiles (25th and 75th percentiles), and the whiskers extend to 1.5 times the interquartile range from the box. AFR, African ancestry; AMR, American ancestry; EAS, East Asian ancestry; EUR, European ancestry; SAS, South Asian ancestry,.

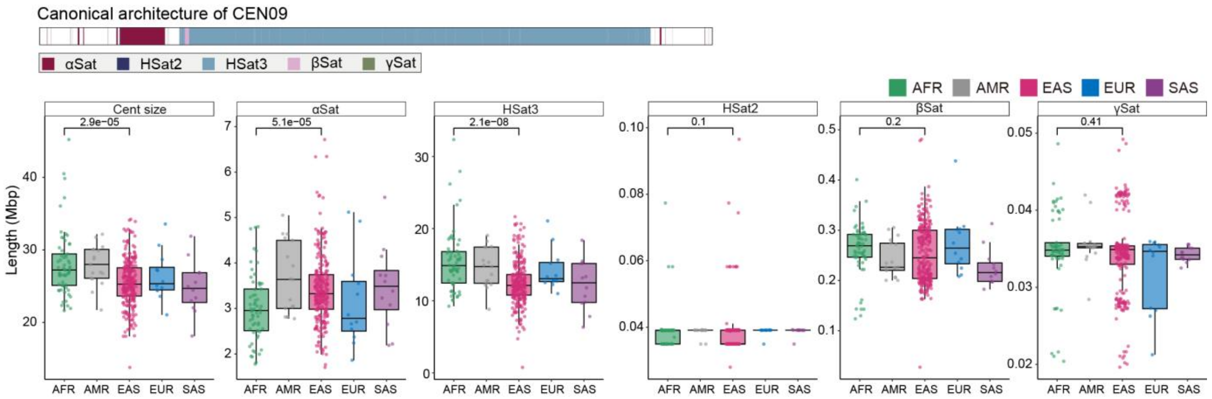


**Supplementary Fig. 6 | Size variation in HSat3 arrays of CEN09.** CEN09 exhibits significantly smaller size in EAS than that in AFR ($p = 2.9 \times 10^{-5}$, Wilcoxon rank-sum test), attributing to the variation of HSat3 array sizes.

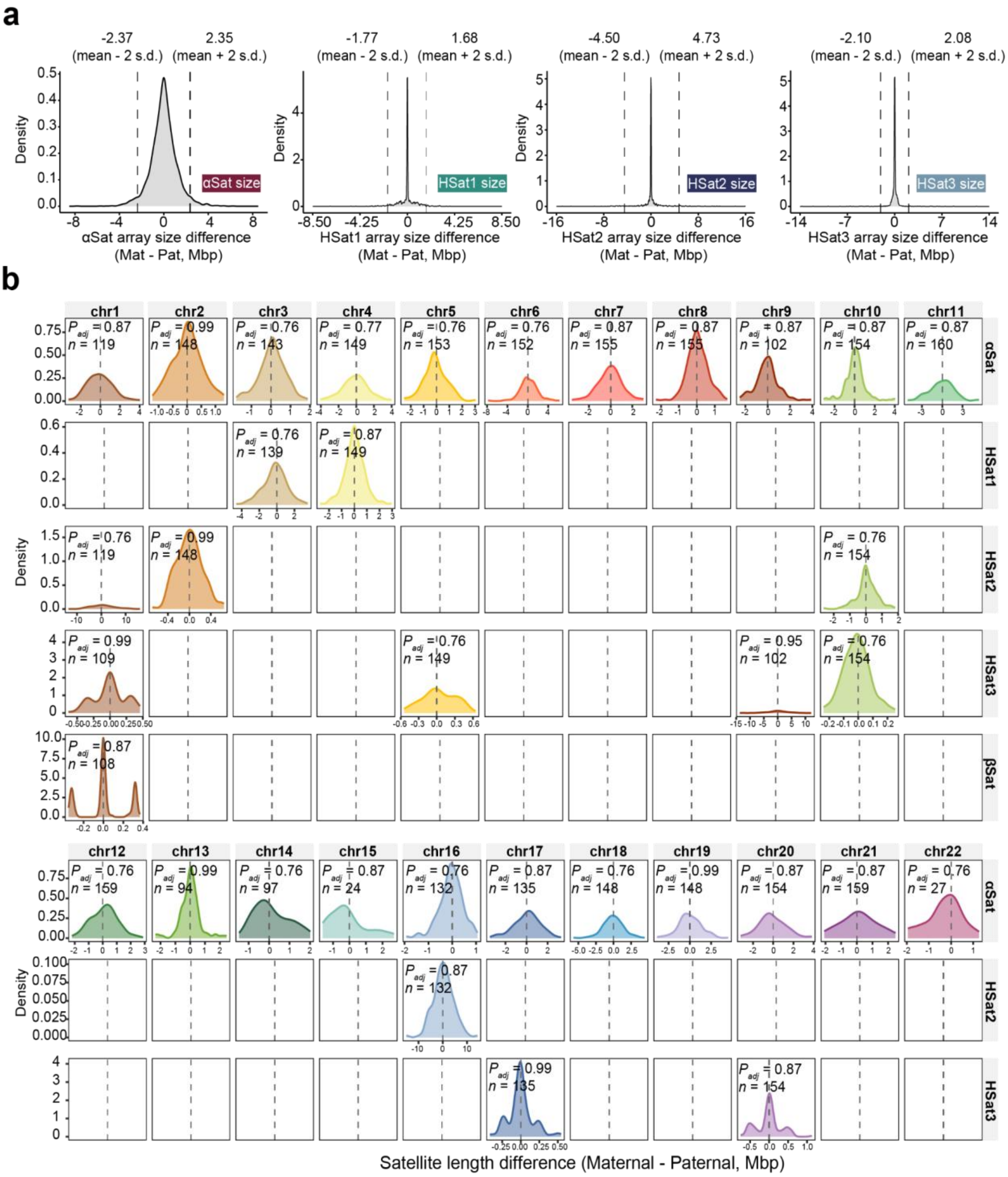

**Supplementary Fig. 7 | Satellite array size differences between maternal and paternal haplotypes across autosomes in APGp1.** a, Density plot of the αSat, HSat1, HSat2 and HSat3 array size difference between maternal(Mat) and paternal(Pat) centromere pairs. Dashed lines demarcate the mean ± 2 s.d. (standard deviations) interval. b, Density distribution of the maternal-paternal length difference ($\Delta = L_{\mathrm{Mat}} - L_{\mathrm{Pat}}$, Mbp) across each chromosome × satellite combination. Rows correspond to satellite types (αSat, HSat1, HSat2, HSat3, and βSat), and columns to autosomes. Only combinations whose mean array length exceeded 500 Kbp were analyzed. The five acrocentric chromosomes (chromosomes 13, 14, 15, 21 and 22) were restricted to αSat, due to assembly limitation within short arms. Each chromosome × satellite combination was tested for a directional Mat/Pat bias using a two-sided one-sample Student's t-test of Δ against $\mu = 0$, with multiple-testing correction across the 35 retained combinations using the Benjamini–Hochberg procedure. The BH-adjusted $p$ value and number of complete Mat/Pat pairs per combination are shown in the top-left corner of each panel. The dashed vertical lines indicate Δ = 0.

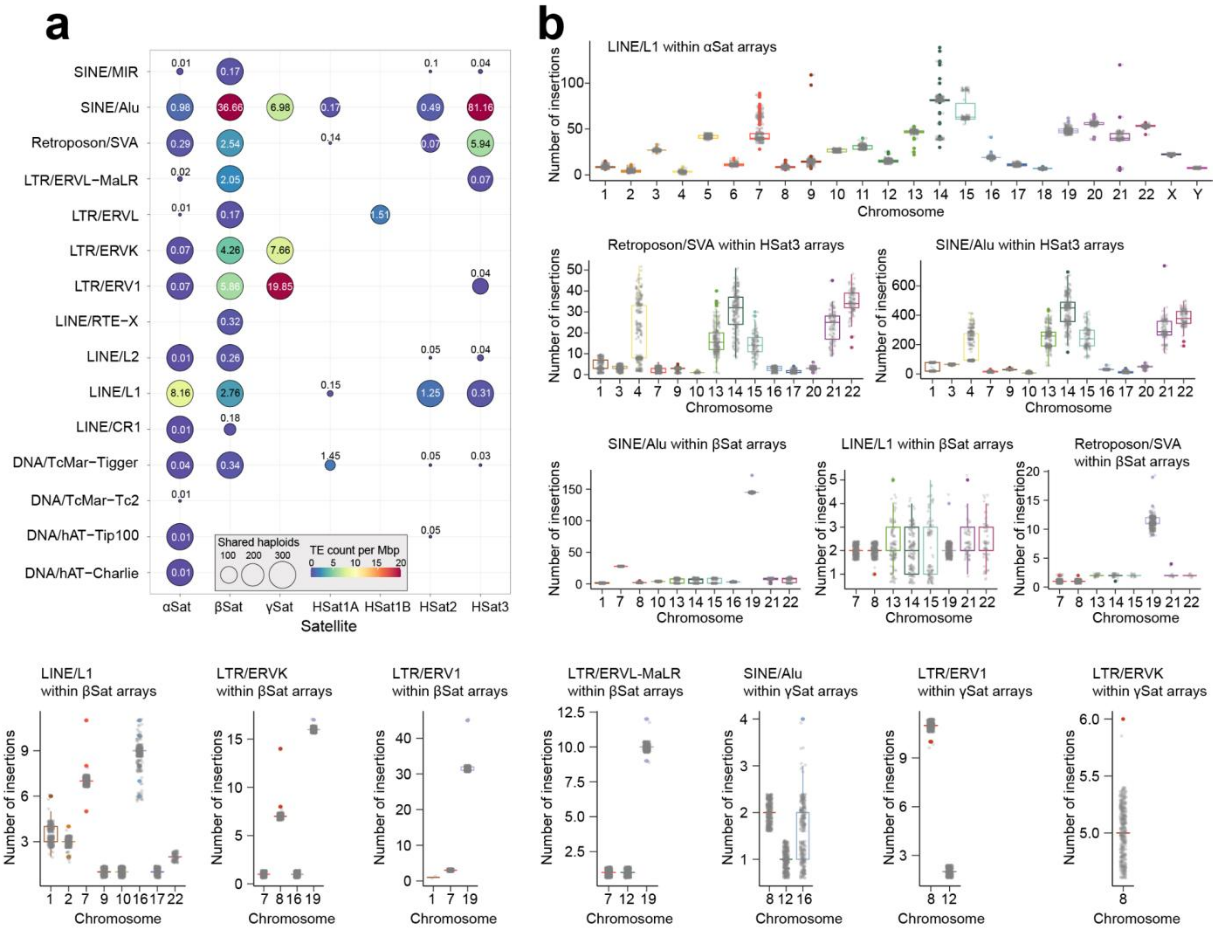


**Supplementary Fig. 8 | Transposable element profiles in human centromeric satellite arrays. a**, Counts of TE insertions by types within different satellite arrays. Circle sizes indicate the numbers of haploid assemblies carrying the insertions. Colors and numbers within the circle represent the centromeric TE counts per megabase within each satellite family. **b**, Distribution of TE insertion counts per chromosome. While the overall number of TE insertions is conserved across the population, variable patterns are observed, for example, LINE/L1 in CEN07 (αSat arrays) and CEN16 (HSat2 arrays), and Alu/SVA in CEN04 (HSat3 arrays). Note that TE insertion counts in acrocentric centromeres should be interpreted with caution, considering the potentially incomplete assembly near telomeric regions.

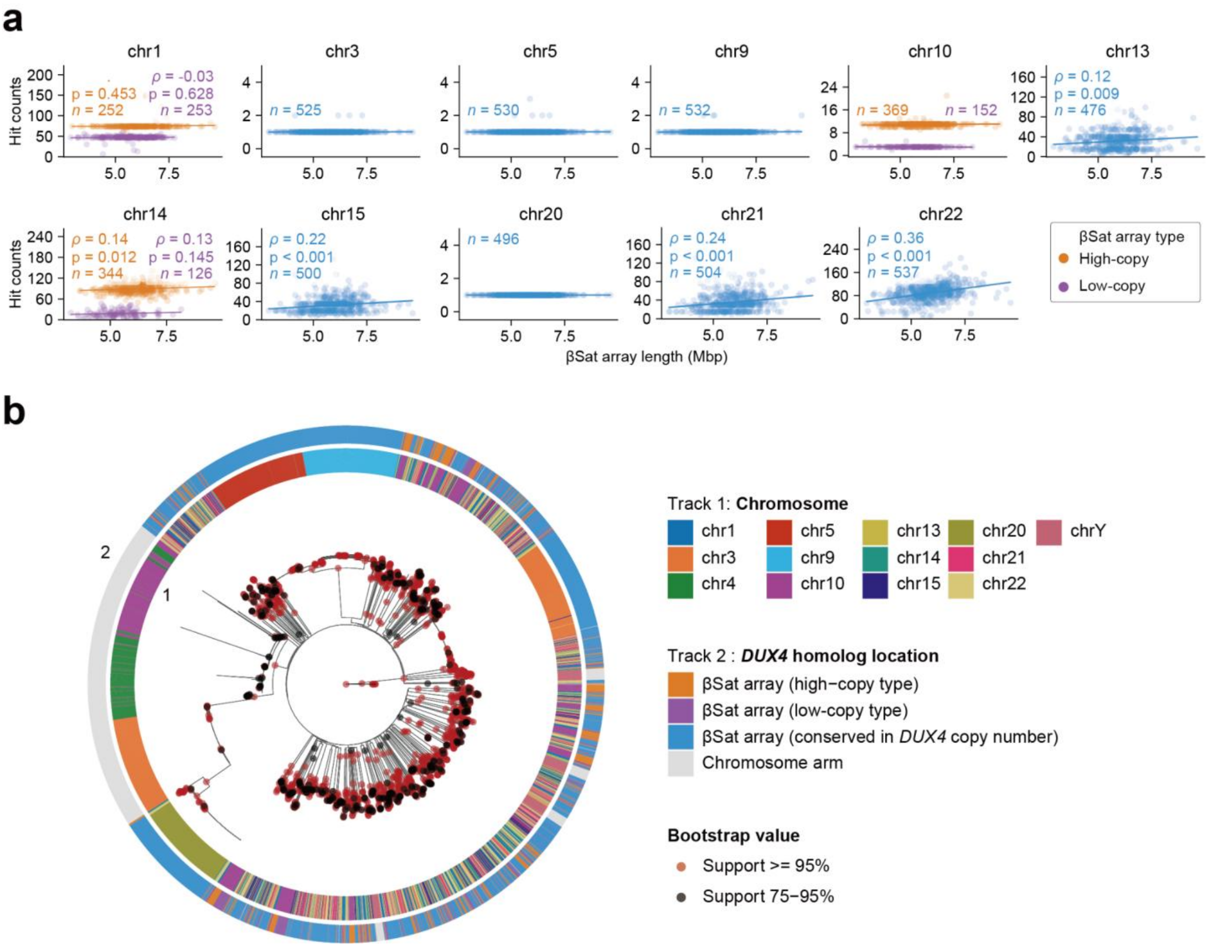


**Supplementary Fig. 9 | Copy number variation and phylogenetic relationships of *DUX4* homologs in human βsat arrays. a**, Scatter plots showing the relationship between βSat region length and *DUX4* Coding sequence hit count across different chromosomes. Spearman correlation coefficients (*ρ*) and *p*-values (derived from a two-sided Spearman's rank correlation test) are displayed. *DUX4* copy numbers on acrocentric chromosomes (*e. g.* 13, 15, 21 and 22) show significantly positive correlations with βSat region lengths. Some chromosomes (e. g. 1, 10 and 14) exhibit discrete, multi-layered copy number states. In contrast, other chromosomes (*e. g.* 3, 5, 9 and 20) are almost fixed with low copy number. **b**, A phylogenetic tree of *DUX4* homologs in βSat arrays and functional subtelomeric *DUX4* orthologs from chromosomes 4 and 10. Node bootstrap support is indicated by red (>= 95%) and grey (75-95%). Annotation tracks from most to outermost, represent: (1) chromosome (color-coded), locations of *DUX4* homologs. For example, a sequence associated with an orange inner ring (chr3) and a grey outer ring (Ortholog) represents a *DUX4* ortholog from chromosome 3. The phylogeny shows that homologs from stable, low-copy regions (*e. g.* chromosomes 3, 9 and 20) cluster with functional orthologs (*e. g.* chromosomes 4 and 10) into conserved clades. In contrast, homologs from the high-copy, actively expanding arrays (*e. g.* chromosomes 13, 14, 15 and 21) formed complex, rapidly evolving clades.

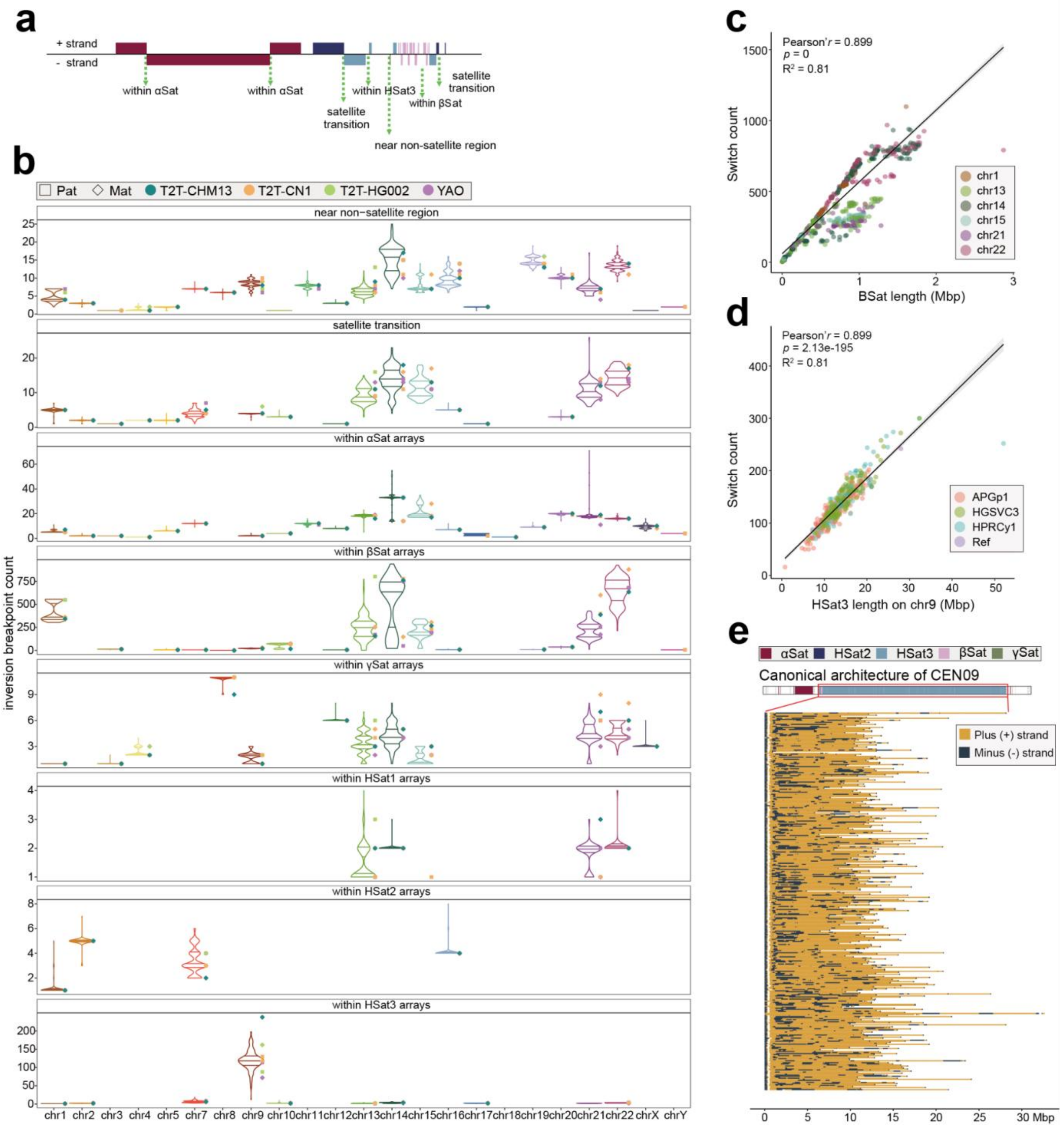


**Supplementary Fig. 10 | Sequence strand switch breakpoints in human centromeric satellite arrays. a**, Schematic of sequence strand switches as a proxy for rearrangement breakpoints. **b**, The number of strand switch breakpoints within and between satellite arrays across haploid assemblies.In each violin plot, the solid lines denote the 25th percentile, median, and 75th percentile, respectively. **c** and **d**, Correlation between strand switch breakpoint counts and array length in (**c**) βSat and (**d**) HSat3 arrays in CEN09. **e**, Strand switches in HSat3 arrays of CEN09 are randomly distributed rather than localized to hotspots.

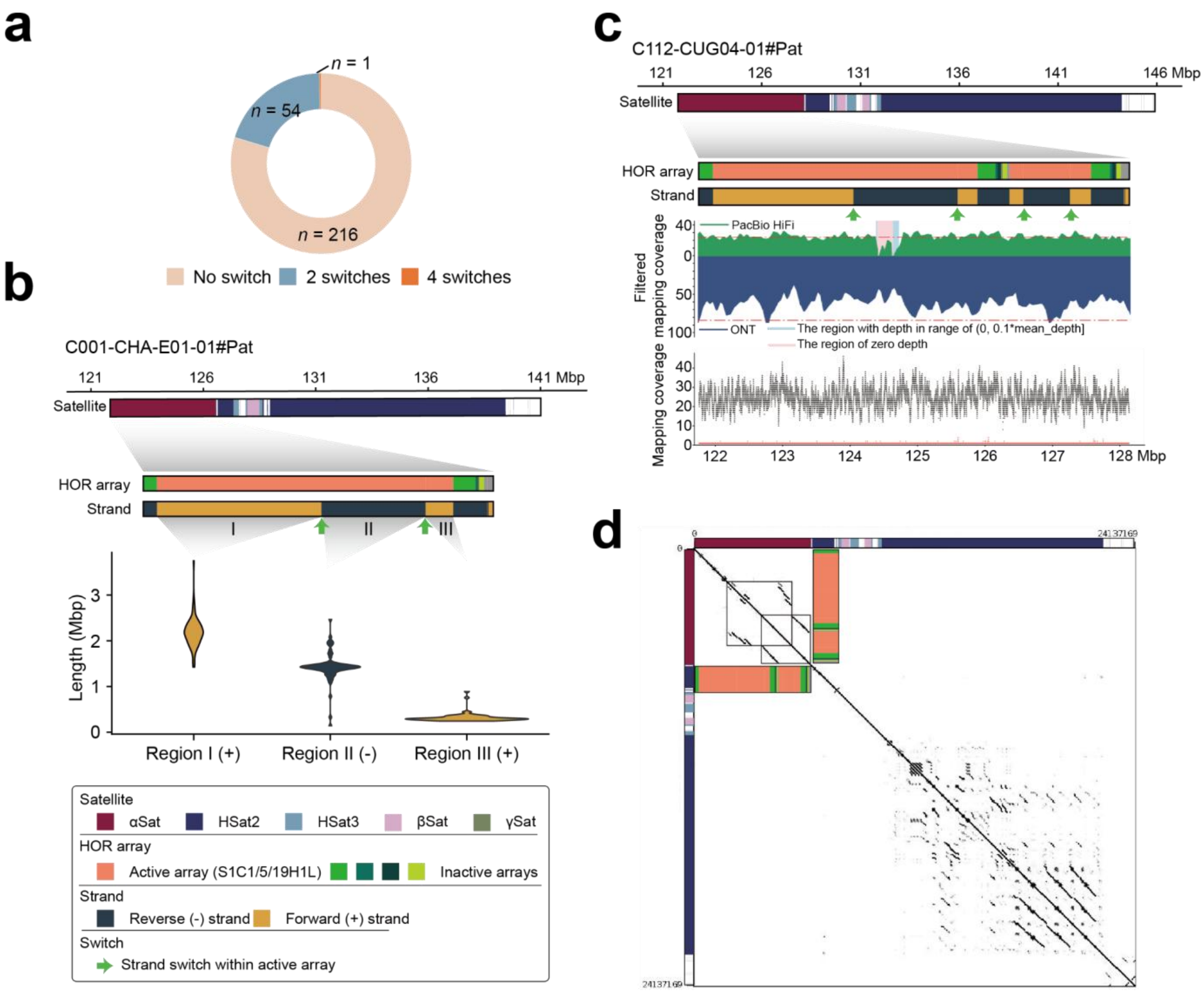


**Supplementary Fig. 11 | Polymorphic megabase-scale strand inversion inside the active HOR arrays of CEN01. a**, Distribution of haplotype assemblies by the number of strand switches (0, 2, or 4) detected inside the active HOR arrays of CEN01. **b**, Examples of two strand switches within the active arrays of CEN01 (e. g. C001-CHA-E01-01#Pat), accompanied by a violin plot comparing the genomic lengths of three segments within this region. **c**, A single haplotype assembly (i. e. C112-CUG04-01#Pat) showing four strand switches within the active array. The assembly was validated by GCI and NucFreq, with junction loci displaying even read coverage and no notable increase of heterozygous sites. **d**, Self-alignment dotplot of CEN01 (C112-CUG04-01#Pat, word size = 1,000), revealing large duplications that span both the inverted active HOR array and adjacent inactive HOR regions.

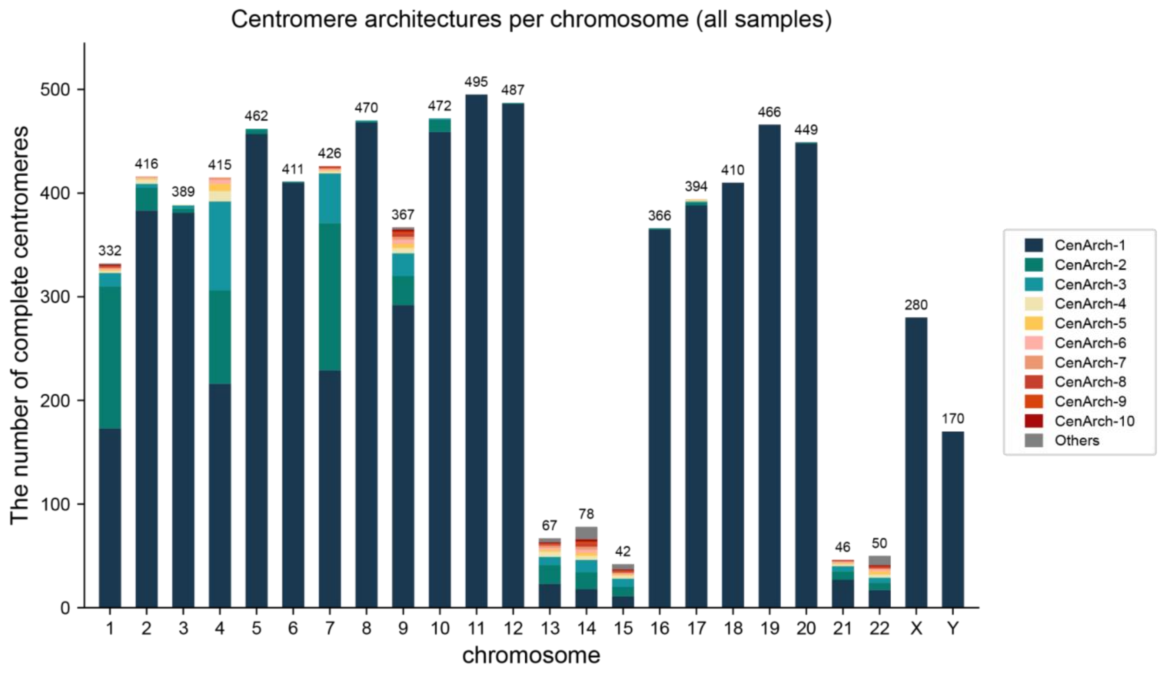


**Supplementary Fig. 12 | Distribution of centromere architectures (CenArchs) across chromosomes in all samples.** Colours and architecture indices denote the chromosome-specific centromere architectures, ordered by decreasing haplotype frequency, with the total number of complete centromeres indicated above each bar. For acrocentric chromosomes (13, 14, 15, 21 and 22), beyond the gapless criterion, we additionally removed any centromere lacking annotated telomere or rDNA to avoid misclassification due to incomplete assembly of peri-telomeric satellite sequences.

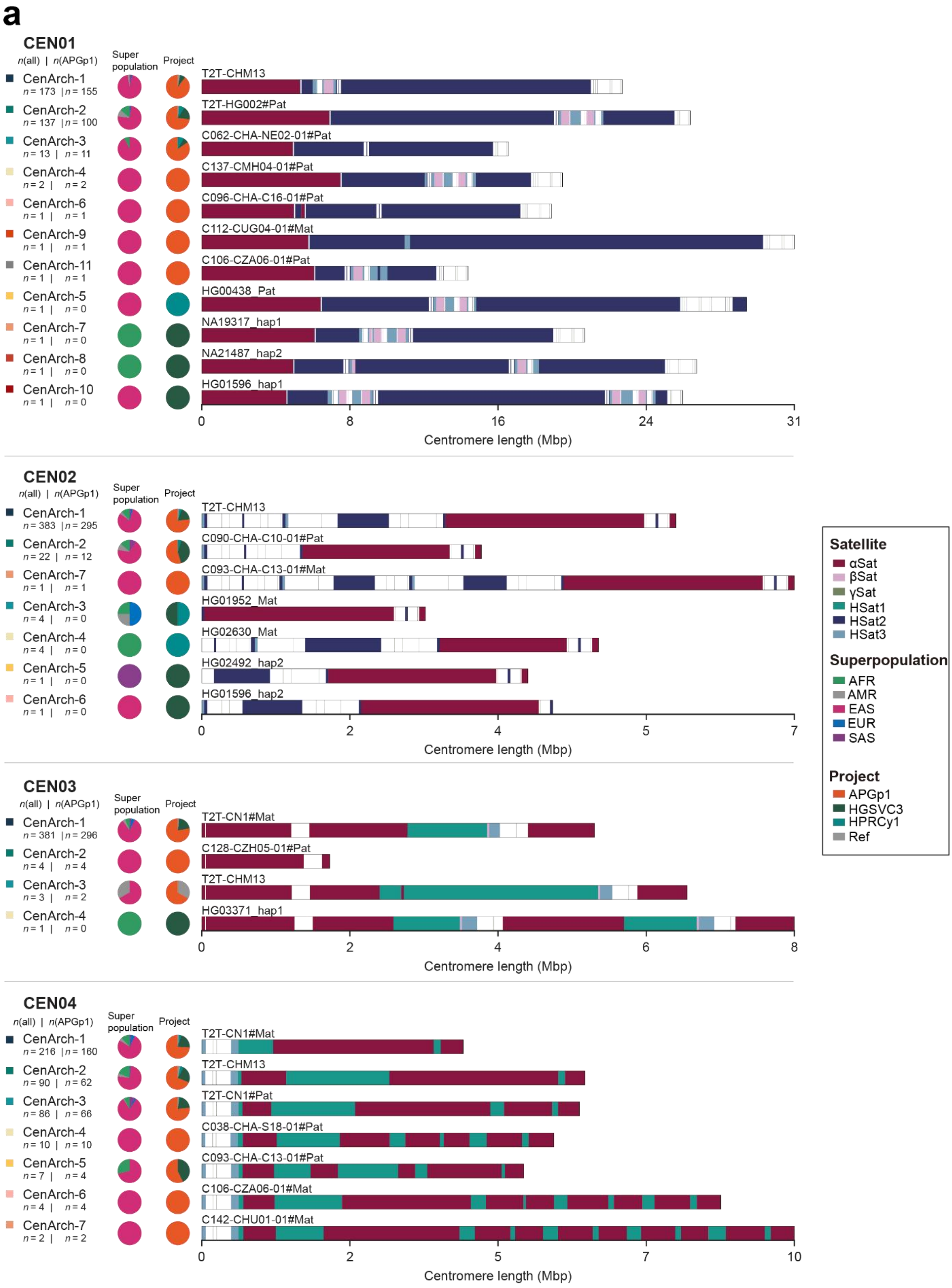
a
CEN01
n(all) | n(APGp1)
Super population
Project
CenArch-1
n = 173 | n = 155
CenArch-2
n = 137 | n = 100
CenArch-3
n = 13 | n = 11
CenArch-4
n = 2 | n = 2
CenArch-6
n = 1 | n = 1
CenArch-9
n = 1 | n = 1
CenArch-11
n = 1 | n = 1
CenArch-5
n = 1 | n = 0
CenArch-7
n = 1 | n = 0
CenArch-8
n = 1 | n = 0
CenArch-10
n = 1 | n = 0
T2T-CHM13
T2T-HG002#Pat
C062-CHA-NE02-01#Pat
C137-CMH04-01#Pat
C096-CHA-C16-01#Pat
C112-CUG04-01#Mat
C106-CZA06-01#Pat
HG00438_Pat
NA19317_hap1
NA21487_hap2
HG01596_hap1
0
8
16
24
31
Centromere length (Mbp)
CEN02
n(all) | n(APGp1)
Super population
Project
CenArch-1
n = 383 | n = 295
CenArch-2
n = 22 | n = 12
CenArch-7
n = 1 | n = 1
CenArch-3
n = 4 | n = 0
CenArch-4
n = 4 | n = 0
CenArch-5
n = 1 | n = 0
CenArch-6
n = 1 | n = 0
T2T-CHM13
C090-CHA-C10-01#Pat
C093-CHA-C13-01#Mat
HG01952_Mat
HG02630_Mat
HG02492_hap2
HG01596_hap2
0
2
4
5
7
Centromere length (Mbp)
Satellite
αSat
βSat
γSat
HSat1
HSat2
HSat3
Superpopulation
AFR
AMR
EAS
EUR
SAS
Project
APGp1
HGSVC3
HPRCy1
Ref
CEN03
n(all) | n(APGp1)
Super population
Project
CenArch-1
n = 381 | n = 296
CenArch-2
n = 4 | n = 4
CenArch-3
n = 3 | n = 2
CenArch-4
n = 1 | n = 0
T2T-CN1#Mat
C128-CZH05-01#Pat
T2T-CHM13
HG03371_hap1
0
2
4
6
8
Centromere length (Mbp)
CEN04
n(all) | n(APGp1)
Super population
Project
CenArch-1
n = 216 | n = 160
CenArch-2
n = 90 | n = 62
CenArch-3
n = 86 | n = 66
CenArch-4
n = 10 | n = 10
CenArch-5
n = 7 | n = 4
CenArch-6
n = 4 | n = 4
CenArch-7
n = 2 | n = 2
T2T-CN1#Mat
T2T-CHM13
T2T-CN1#Pat
C038-CHA-S18-01#Pat
C093-CHA-C13-01#Pat
C106-CZA06-01#Mat
C142-CHU01-01#Mat
0
2
5
7
10
Centromere length (Mbp)

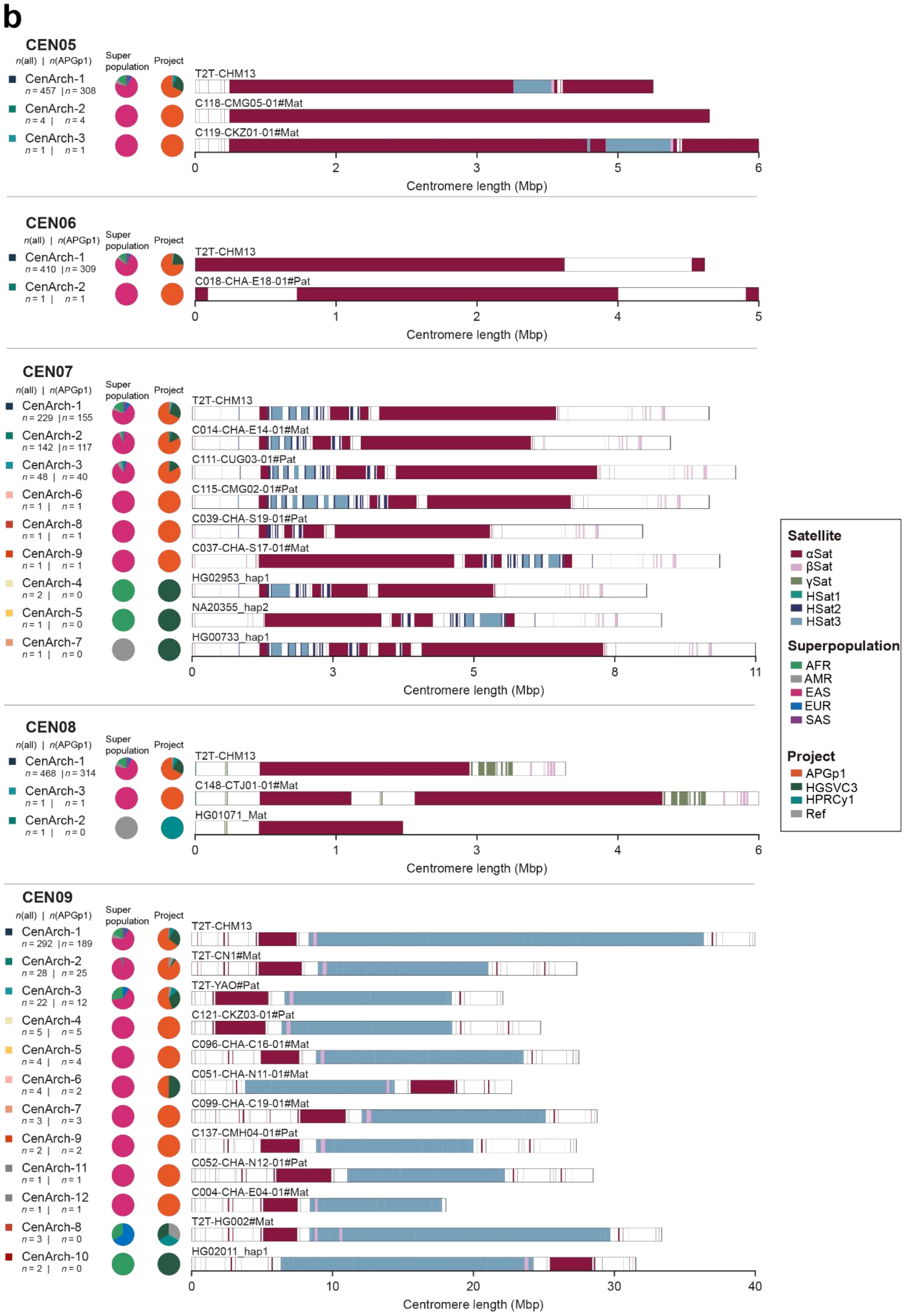

b
CEN05
n(all) | n(APGp1)
Super population
Project
CenArch-1 n = 457 | n = 308
CenArch-2 n = 4 | n = 4
CenArch-3 n = 1 | n = 1
T2T-CHM13
C118-CMG05-01#Mat
C119-CKZ01-01#Mat
Centromere length (Mbp)
CEN06
CenArch-1 n = 410 | n = 309
CenArch-2 n = 1 | n = 1
T2T-CHM13
C018-CHA-E18-01#Pat
CEN07
CenArch-1 n = 229 | n = 155
CenArch-2 n = 142 | n = 117
CenArch-3 n = 48 | n = 40
CenArch-6 n = 1 | n = 1
CenArch-8 n = 1 | n = 1
CenArch-9 n = 1 | n = 1
CenArch-4 n = 2 | n = 0
CenArch-5 n = 1 | n = 0
CenArch-7 n = 1 | n = 0
T2T-CHM13
C014-CHA-E14-01#Mat
C111-CUG03-01#Pat
C115-CMG02-01#Pat
C039-CHA-S19-01#Pat
C037-CHA-S17-01#Mat
HG02953_hap1
NA20355_hap2
HG00733_hap1
CEN08
CenArch-1 n = 468 | n = 314
CenArch-3 n = 1 | n = 1
CenArch-2 n = 1 | n = 0
T2T-CHM13
C148-CTJ01-01#Mat
HG01071_Mat
CEN09
CenArch-1 n = 292 | n = 189
CenArch-2 n = 28 | n = 25
CenArch-3 n = 22 | n = 12
CenArch-4 n = 5 | n = 5
CenArch-5 n = 4 | n = 4
CenArch-6 n = 4 | n = 2
CenArch-7 n = 3 | n = 3
CenArch-9 n = 2 | n = 2
CenArch-11 n = 1 | n = 1
CenArch-12 n = 1 | n = 1
CenArch-8 n = 3 | n = 0
CenArch-10 n = 2 | n = 0
T2T-CHM13
T2T-CN1#Mat
T2T-YAO#Pat
C121-CKZ03-01#Pat
C096-CHA-C16-01#Mat
C051-CHA-N11-01#Mat
C099-CHA-C19-01#Mat
C137-CMH04-01#Pat
C052-CHA-N12-01#Pat
C004-CHA-E04-01#Mat
T2T-HG002#Mat
HG02011_hap1
Satellite
αSat
βSat
γSat
HSat1
HSat2
HSat3
Superpopulation
AFR
AMR
EAS
EUR
SAS
Project
APGp1
HGSVC3
HPRCy1
Ref

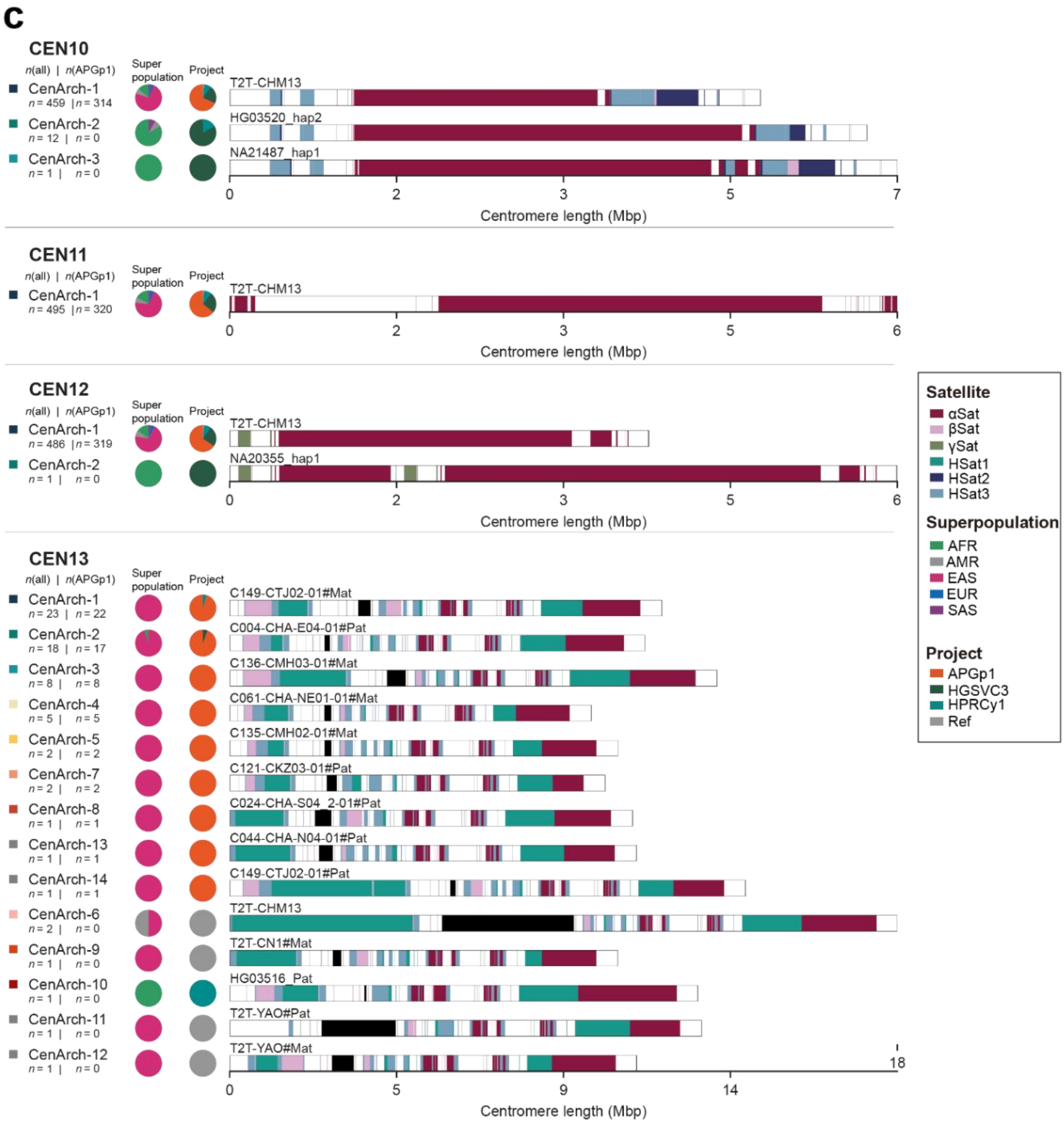
c
CEN10
n(all) | n(APGp1)
Super population
Project
CenArch-1
n = 459 | n = 314
CenArch-2
n = 12 | n = 0
CenArch-3
n = 1 | n = 0
T2T-CHM13
HG03520_hap2
NA21487_hap1
0
2
3
5
7
Centromere length (Mbp)
CEN11
n(all) | n(APGp1)
Super population
Project
CenArch-1
n = 495 | n = 320
T2T-CHM13
0
2
3
5
6
Centromere length (Mbp)
CEN12
n(all) | n(APGp1)
Super population
Project
CenArch-1
n = 486 | n = 319
CenArch-2
n = 1 | n = 0
T2T-CHM13
NA20355_hap1
0
2
3
5
6
Centromere length (Mbp)
CEN13
n(all) | n(APGp1)
Super population
Project
CenArch-1
n = 23 | n = 22
CenArch-2
n = 18 | n = 17
CenArch-3
n = 8 | n = 8
CenArch-4
n = 5 | n = 5
CenArch-5
n = 2 | n = 2
CenArch-7
n = 2 | n = 2
CenArch-8
n = 1 | n = 1
CenArch-13
n = 1 | n = 1
CenArch-14
n = 1 | n = 1
CenArch-6
n = 2 | n = 0
CenArch-9
n = 1 | n = 0
CenArch-10
n = 1 | n = 0
CenArch-11
n = 1 | n = 0
CenArch-12
n = 1 | n = 0
C149-CTJ02-01#Mat
C004-CHA-E04-01#Pat
C136-CMH03-01#Mat
C061-CHA-NE01-01#Mat
C135-CMH02-01#Mat
C121-CKZ03-01#Pat
C024-CHA-S04_2-01#Pat
C044-CHA-N04-01#Pat
C149-CTJ02-01#Pat
T2T-CHM13
T2T-CN1#Mat
HG03516_Pat
T2T-YAO#Pat
T2T-YAO#Mat
18
0
5
9
14
Centromere length (Mbp)
Satellite
αSat
βSat
γSat
HSat1
HSat2
HSat3
Superpopulation
AFR
AMR
EAS
EUR
SAS
Project
APGp1
HGSVC3
HPRCy1
Ref

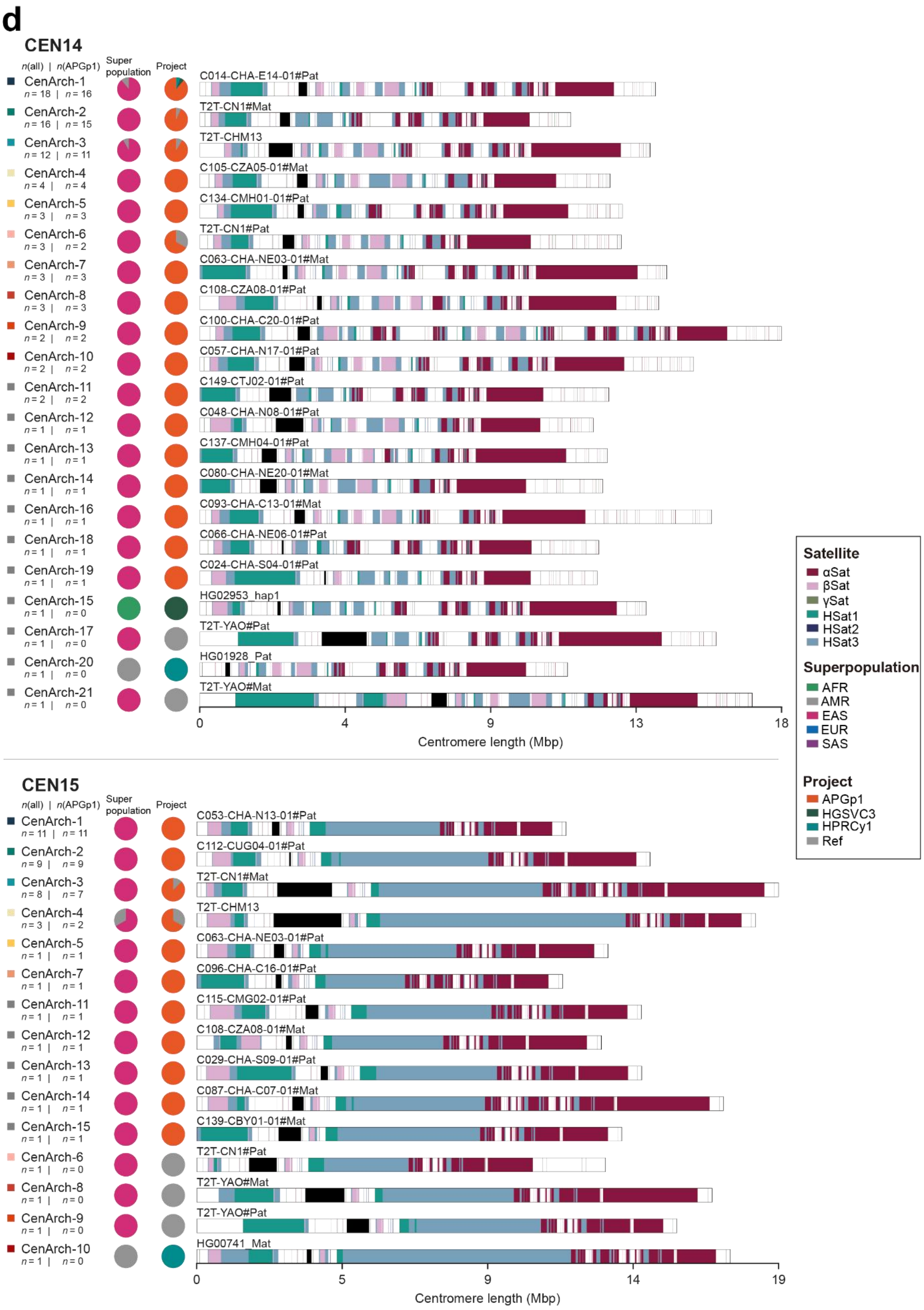
d
CEN14
n(all) | n(APGp1)
Super population
Project
CenArch-1 n = 18 | n = 16
CenArch-2 n = 16 | n = 15
CenArch-3 n = 12 | n = 11
CenArch-4 n = 4 | n = 4
CenArch-5 n = 3 | n = 3
CenArch-6 n = 3 | n = 2
CenArch-7 n = 3 | n = 3
CenArch-8 n = 3 | n = 3
CenArch-9 n = 2 | n = 2
CenArch-10 n = 2 | n = 2
CenArch-11 n = 2 | n = 2
CenArch-12 n = 1 | n = 1
CenArch-13 n = 1 | n = 1
CenArch-14 n = 1 | n = 1
CenArch-16 n = 1 | n = 1
CenArch-18 n = 1 | n = 1
CenArch-19 n = 1 | n = 1
CenArch-15 n = 1 | n = 0
CenArch-17 n = 1 | n = 0
CenArch-20 n = 1 | n = 0
CenArch-21 n = 1 | n = 0
C014-CHA-E14-01#Pat
T2T-CN1#Mat
T2T-CHM13
C105-CZA05-01#Mat
C134-CMH01-01#Pat
T2T-CN1#Pat
C063-CHA-NE03-01#Mat
C108-CZA08-01#Pat
C100-CHA-C20-01#Pat
C057-CHA-N17-01#Pat
C149-CTJ02-01#Pat
C048-CHA-N08-01#Pat
C137-CMH04-01#Pat
C080-CHA-NE20-01#Mat
C093-CHA-C13-01#Mat
C066-CHA-NE06-01#Pat
C024-CHA-S04-01#Pat
HG02953_hap1
T2T-YAO#Pat
HG01928_Pat
T2T-YAO#Mat
0
4
9
13
18
Centromere length (Mbp)
CEN15
n(all) | n(APGp1)
Super population
Project
CenArch-1 n = 11 | n = 11
CenArch-2 n = 9 | n = 9
CenArch-3 n = 8 | n = 7
CenArch-4 n = 3 | n = 2
CenArch-5 n = 1 | n = 1
CenArch-7 n = 1 | n = 1
CenArch-11 n = 1 | n = 1
CenArch-12 n = 1 | n = 1
CenArch-13 n = 1 | n = 1
CenArch-14 n = 1 | n = 1
CenArch-15 n = 1 | n = 1
CenArch-6 n = 1 | n = 0
CenArch-8 n = 1 | n = 0
CenArch-9 n = 1 | n = 0
CenArch-10 n = 1 | n = 0
C053-CHA-N13-01#Pat
C112-CUG04-01#Pat
T2T-CN1#Mat
T2T-CHM13
C063-CHA-NE03-01#Pat
C096-CHA-C16-01#Pat
C115-CMG02-01#Pat
C108-CZA08-01#Mat
C029-CHA-S09-01#Pat
C087-CHA-C07-01#Mat
C139-CBY01-01#Pat
T2T-CN1#Pat
T2T-YAO#Mat
T2T-YAO#Pat
HG00741_Mat
0
5
9
14
19
Centromere length (Mbp)
Satellite
αSat
βSat
γSat
HSat1
HSat2
HSat3
Superpopulation
AFR
AMR
EAS
EUR
SAS
Project
APGp1
HGSVC3
HPRCy1
Ref

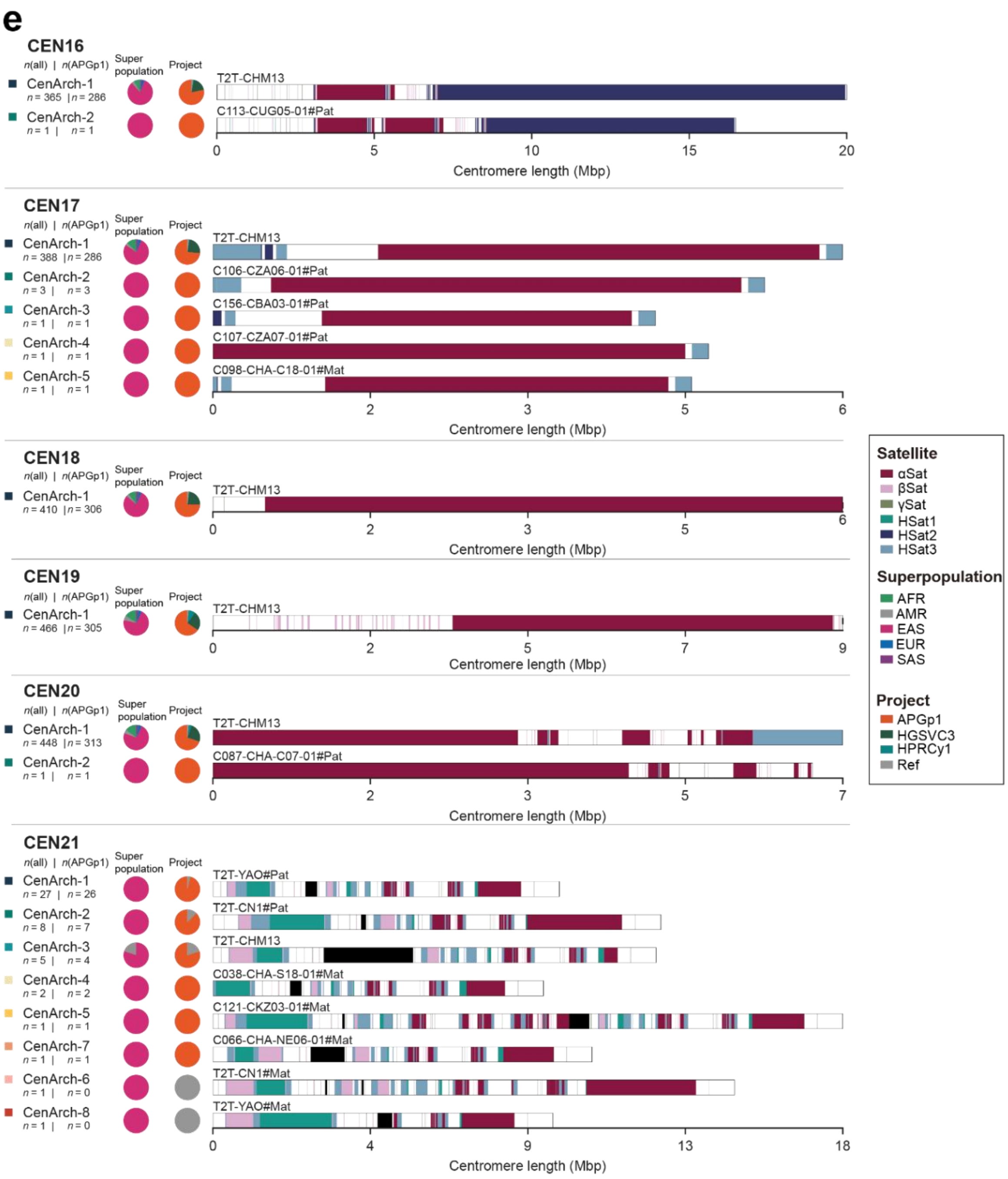

e
CEN16
n(all) | n(APGp1)
Super population
Project
CenArch-1
n = 365 | n = 286
CenArch-2
n = 1 | n = 1
T2T-CHM13
C113-CUG05-01#Pat
0
5
10
15
20
Centromere length (Mbp)
CEN17
n(all) | n(APGp1)
Super population
Project
CenArch-1
n = 388 | n = 286
CenArch-2
n = 3 | n = 3
CenArch-3
n = 1 | n = 1
CenArch-4
n = 1 | n = 1
CenArch-5
n = 1 | n = 1
T2T-CHM13
C106-CZA06-01#Pat
C156-CBA03-01#Pat
C107-CZA07-01#Pat
C098-CHA-C18-01#Mat
0
2
3
5
6
Centromere length (Mbp)
CEN18
n(all) | n(APGp1)
Super population
Project
CenArch-1
n = 410 | n = 306
T2T-CHM13
0
2
3
5
6
Centromere length (Mbp)
CEN19
n(all) | n(APGp1)
Super population
Project
CenArch-1
n = 466 | n = 305
T2T-CHM13
0
2
5
7
9
Centromere length (Mbp)
CEN20
n(all) | n(APGp1)
Super population
Project
CenArch-1
n = 448 | n = 313
CenArch-2
n = 1 | n = 1
T2T-CHM13
C087-CHA-C07-01#Pat
0
2
3
5
7
Centromere length (Mbp)
CEN21
n(all) | n(APGp1)
Super population
Project
CenArch-1
n = 27 | n = 26
CenArch-2
n = 8 | n = 7
CenArch-3
n = 5 | n = 4
CenArch-4
n = 2 | n = 2
CenArch-5
n = 1 | n = 1
CenArch-7
n = 1 | n = 1
CenArch-6
n = 1 | n = 0
CenArch-8
n = 1 | n = 0
T2T-YAO#Pat
T2T-CN1#Pat
T2T-CHM13
C038-CHA-S18-01#Mat
C121-CKZ03-01#Mat
C066-CHA-NE06-01#Mat
T2T-CN1#Mat
T2T-YAO#Mat
0
4
9
13
18
Centromere length (Mbp)
Satellite
αSat
βSat
γSat
HSat1
HSat2
HSat3
Superpopulation
AFR
AMR
EAS
EUR
SAS
Project
APGp1
HGSVC3
HPRCy1
Ref

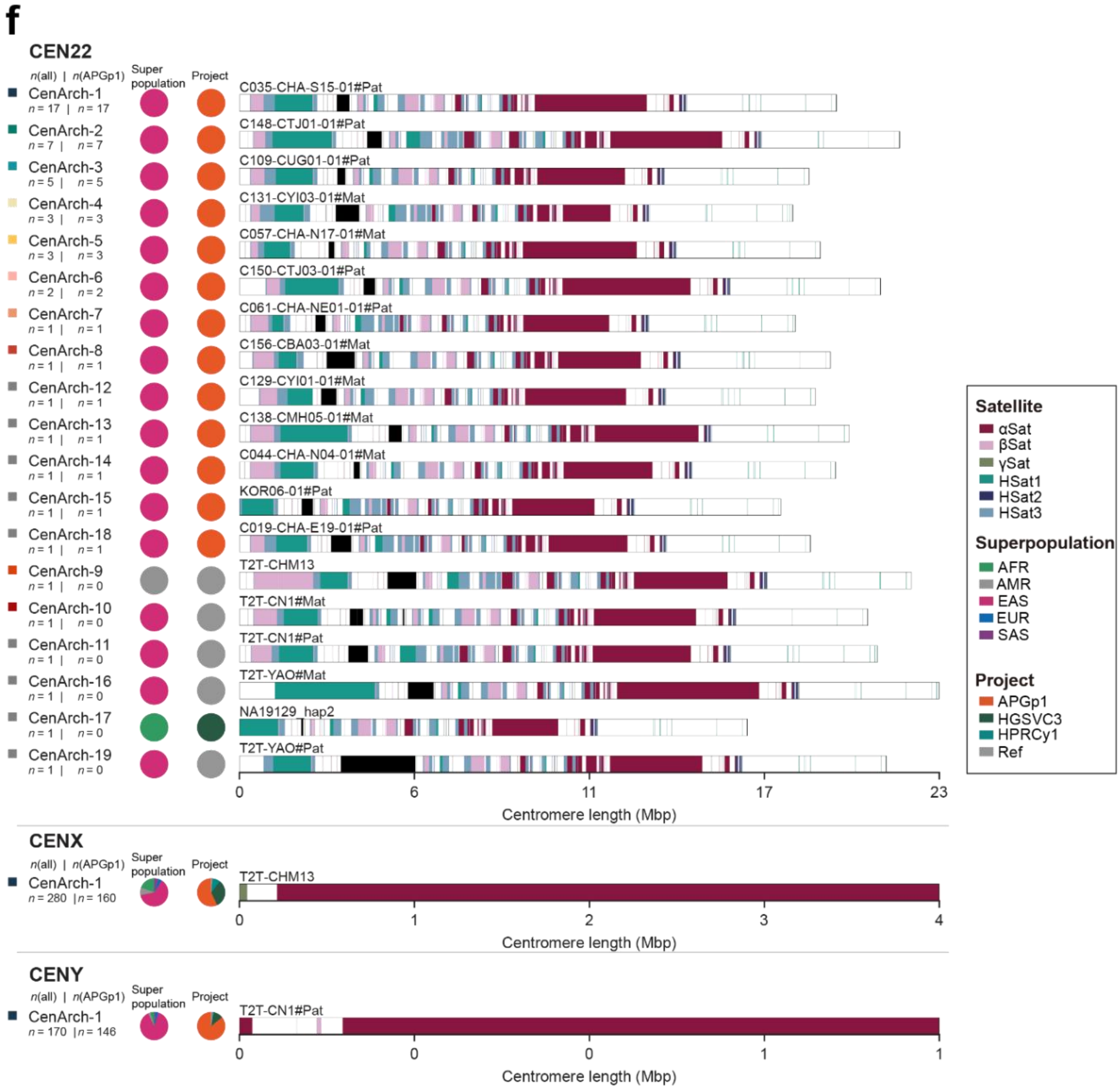


**Supplementary Fig. 13 | Characterization of centromere architectures (CenArchs) based on satellite array organization. a-f**, Centromere architectures of 24 chromosomes. Linear structural tracks map individual architectures ordered by population frequency. Paired numerical matrices to the left denote total haplotype counts across the all samples panel (APGp1, HPRCy1,HGSVC3 and four References) versus the APGp1 cohort, respectively. Twin pie charts track corresponding composition across five global superpopulations and pangenome projects. Architecture indices are assigned in descending order of the number of shared haplotype assemblies (index 1 means most shared), and architectures are ordered for display by project.

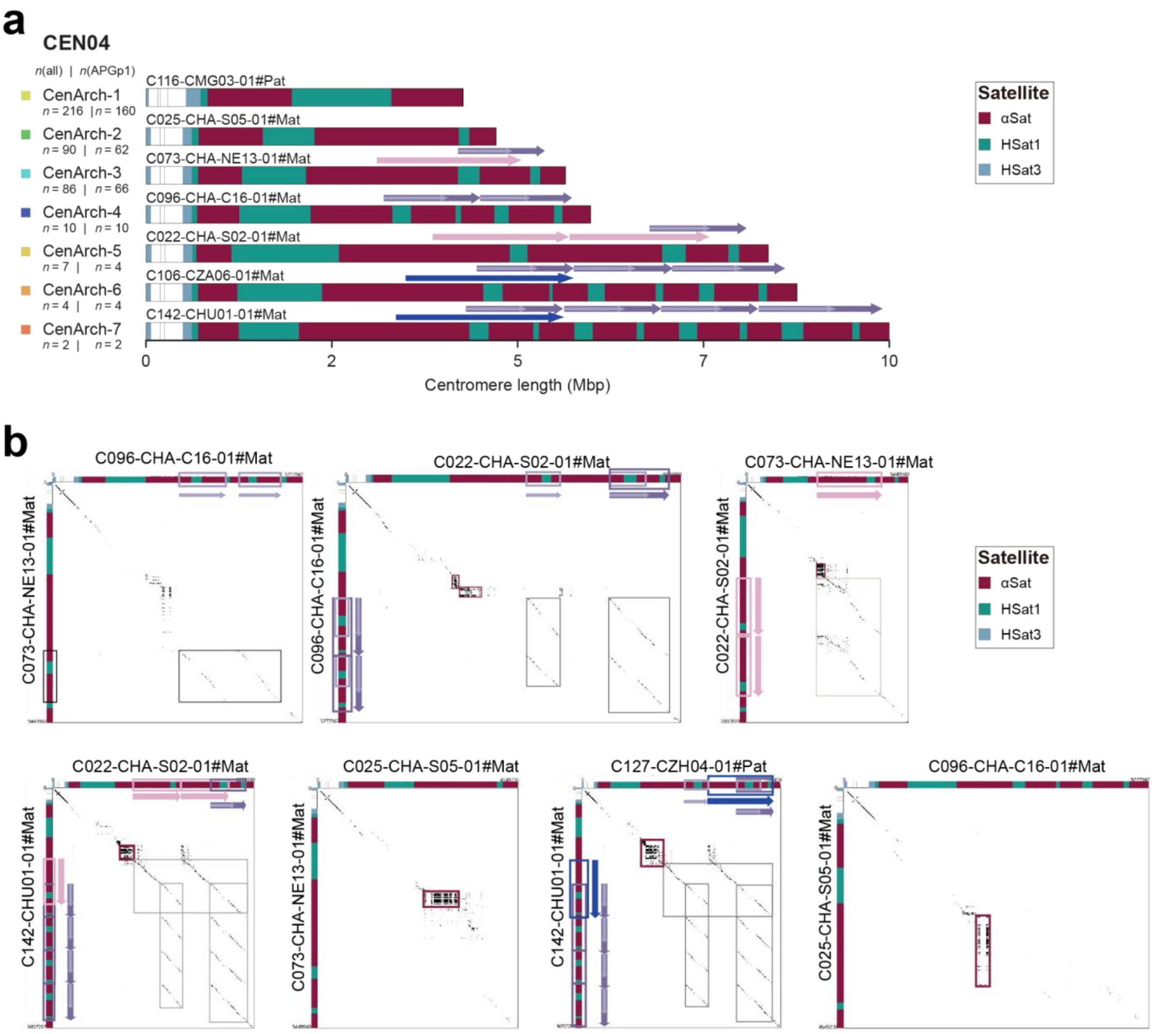


**Supplementary Fig. 14 | Multiple independent large duplication events in CEN04 centromeres. a**, Centromere architectures of CEN04. Linear tracks depict individual CEN04 architectures (CenArch), ordered by population frequency. Paired numerical matrices (left) denote total haplotype counts in the full panel (APGp1, HPRCy1, HGSVC3 and reference assemblies) and in the APGp1 cohort, respectively. Coloured arrows mark large, distinct segmental duplications identified from pairwise alignment dot plots. **b**, Pairwise alignments between representative CEN04 centromere architectures. Dot plots were generated using a word size of 3,429 bp (canonical HOR size in CEN04). Coloured boxes highlight duplicated segments, with colours matching the arrows in **a**. Inconsistency in the boundaries and content of these duplications across individuals indicate that they likely arose from multiple independent events, rather than a single ancestral origin.

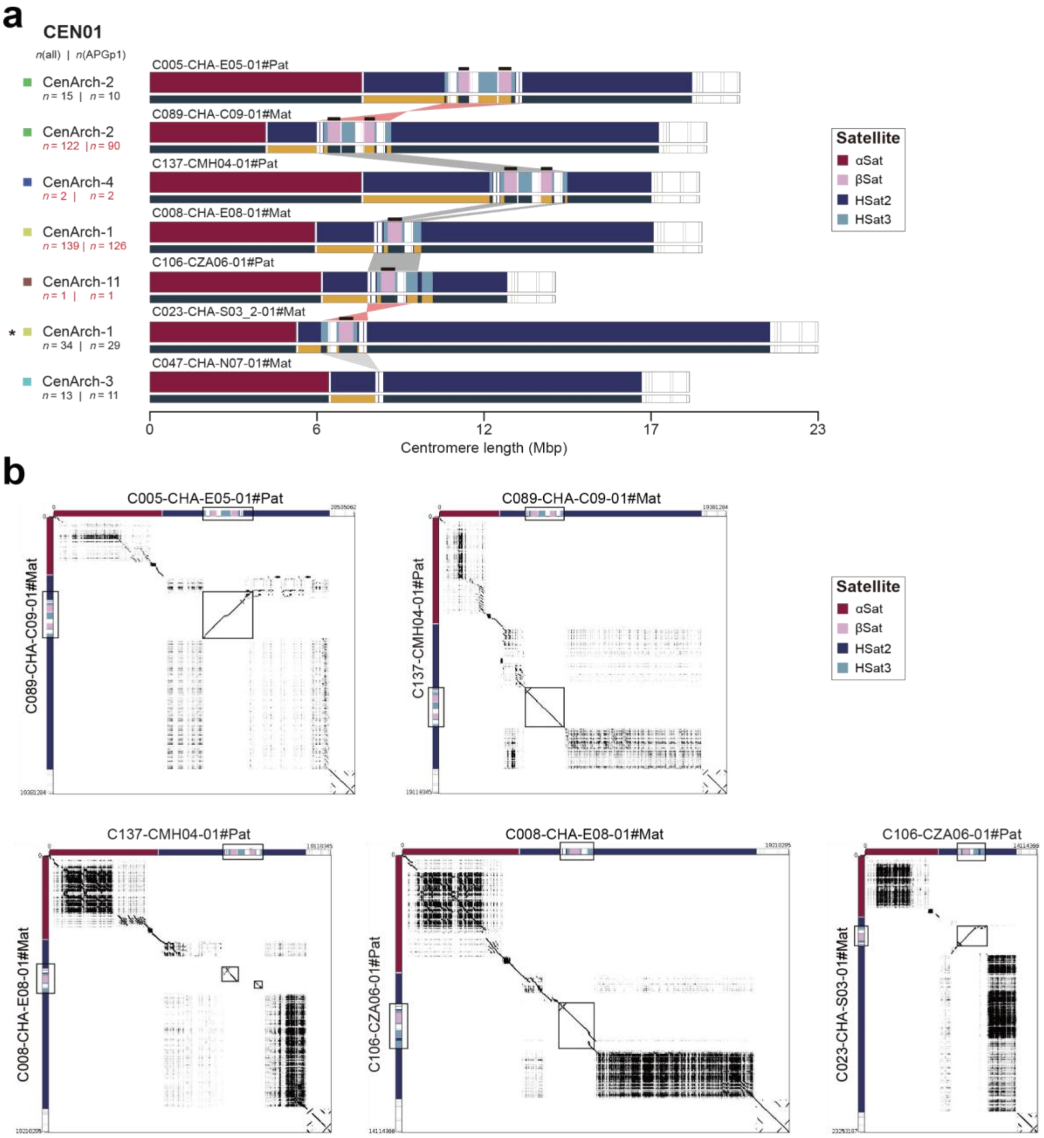


**Supplementary Fig. 15 | Dominant inversion and deletion-mediated βSat copy-number variation shape CEN01 architecture. a,** Linear tracks depicting representative centromere architectures (CenArch) of CEN01. CenArch-1 and CenArch-2 are defined by the presence of one or two β-satellite (βSat) arrays, respectively, with each containing haplotypes both with and without a large inversion relative to T2T-CHM13; one representative example of each combination is shown. CenArch-11 (one βSat array) and CenArch-4 (two βSat arrays) are also displayed; these two architectures were found exclusively with the large inversion. All inversion-bearing architectures are highlighted in red. Paired numerical matrices (left) report the haplotype counts for every CenArch–inversion-status combination in the full dataset (APGp1, HPRCy1, HGSVC3 and reference assemblies; left number) and in the APGp1 cohort (right number). **b**, Pairwise dot-plot alignments of the representative CEN01 architectures. Alignments were computed with a word size of 500 bp to optimally resolve inversion boundaries. Boxes mark inverted segments and alignment regions containing βSat arrays.

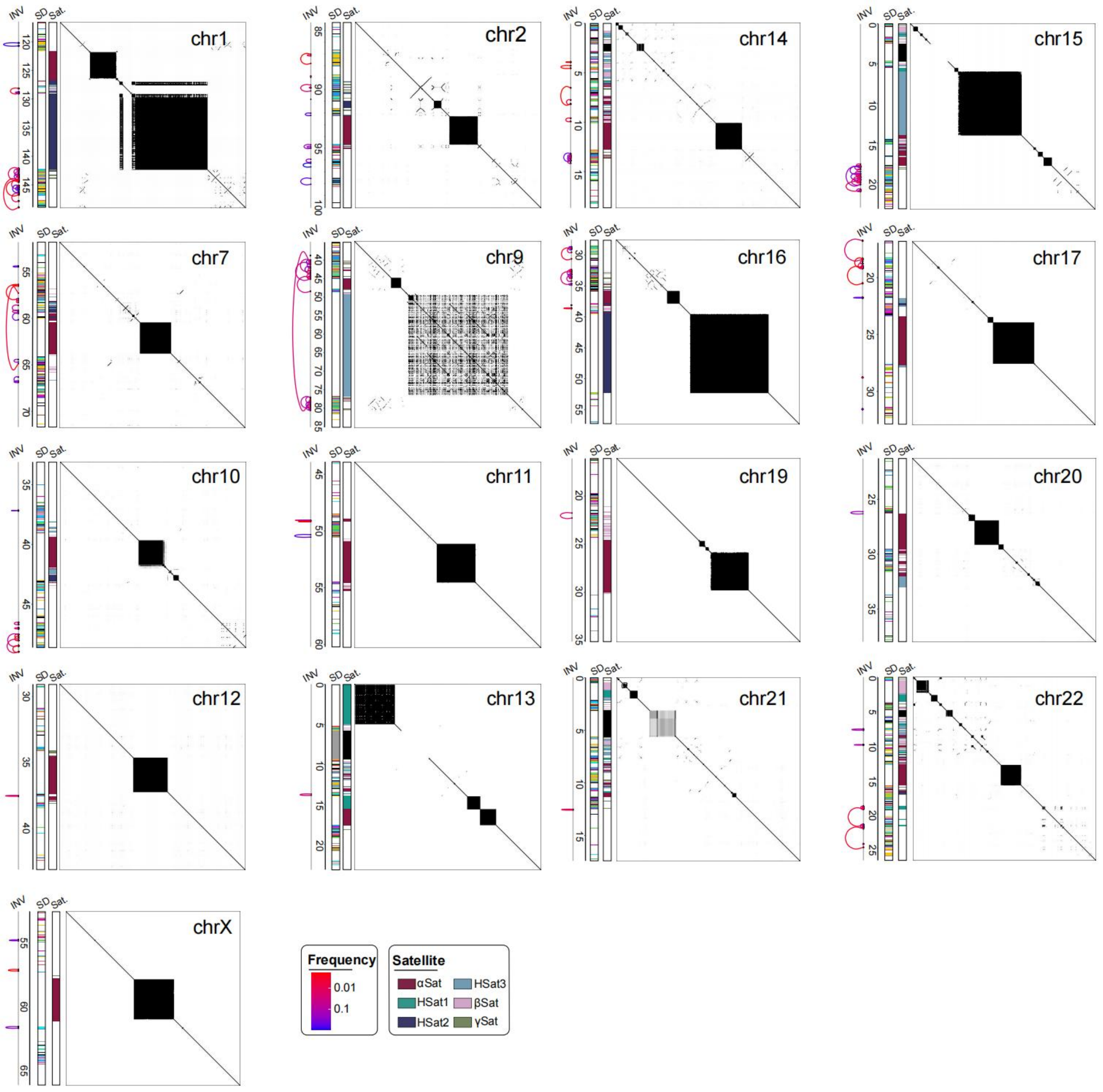


**Supplementary Fig. 16 | Peri/centromeric inversions across human haploid assemblies.** For each chromosome: left, positions of large inversions and their population frequency. Middle, segmental duplication (SD) and satellite (Sat.) annotation. Right, self-alignment dotplots of T2T-CHM13 peri/centromeric sequences.

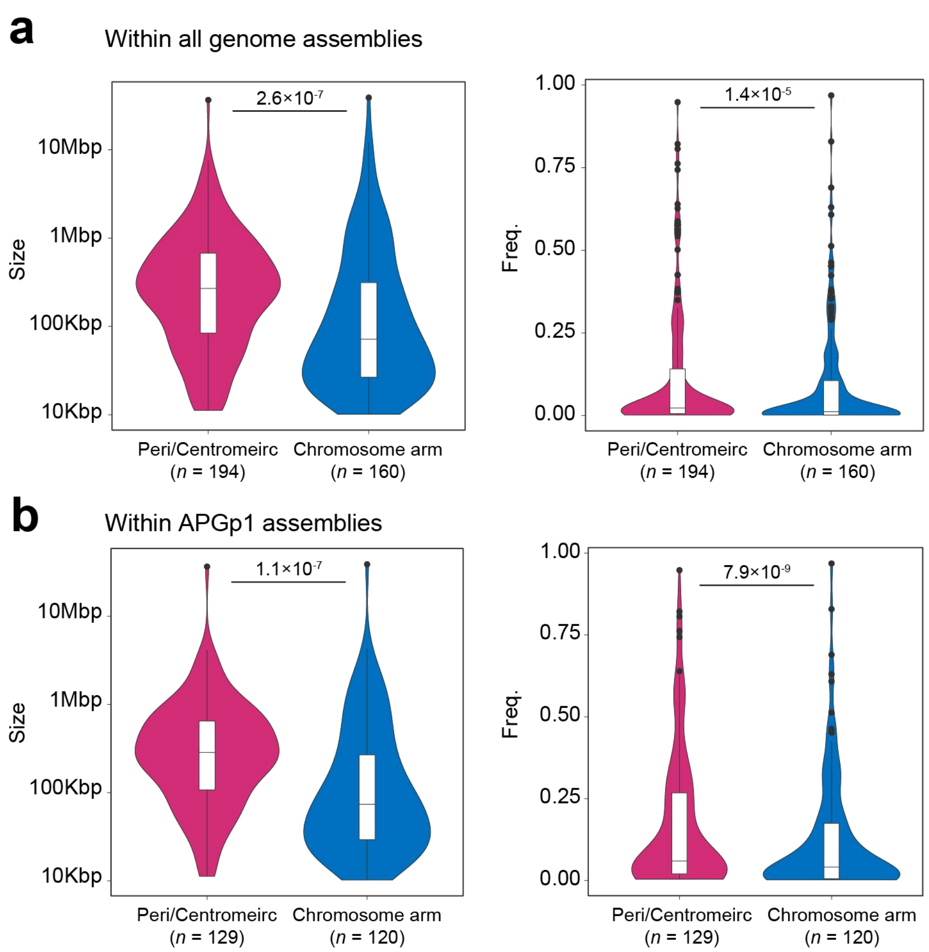


**Supplementary Fig. 17 | Frequency spectrum and sizes of large inversions across chromosome arms and peri/centromeres.** Upper panel (a) shows data for all sampled assemblies, and lower panel (b) shows data for APGp1 assemblies. Compared to the frequency spectrum of large inversions observed at the chromosome arms, the peri/centromeric inversions have higher population frequency and larger genomic size. Statistical significance was determined using a two-sided Wilcoxon rank-sum test.

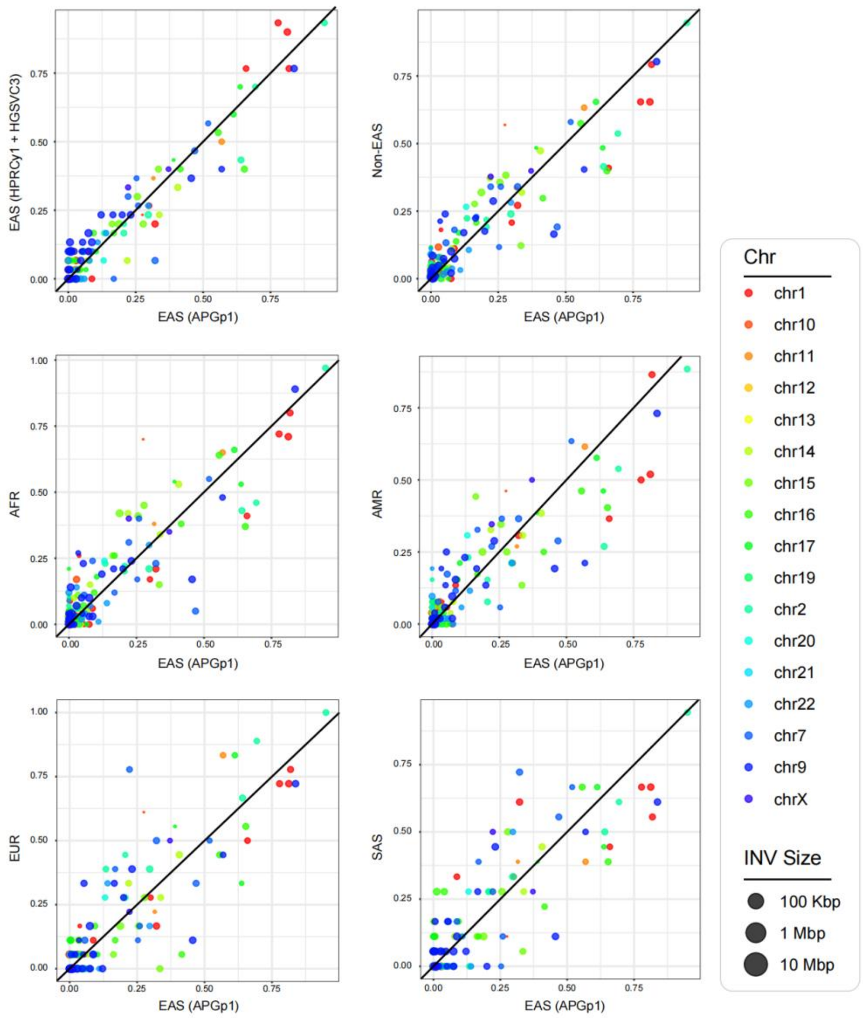


**Supplementary Fig. 18 | Inversion frequency between EAS and other superpopulations.** Comparison of inversion frequencies between EAS superpopulation from APGp1 and other groups: other EAS populations (from HGSVC3 and HPRCy1), non-EAS superpopulations, and other superpopulations (AFR, African; AMR, American; EUR, European; SAS, South Asian). Point sizes represent inversion sizes, and colors indicate chromosomes.

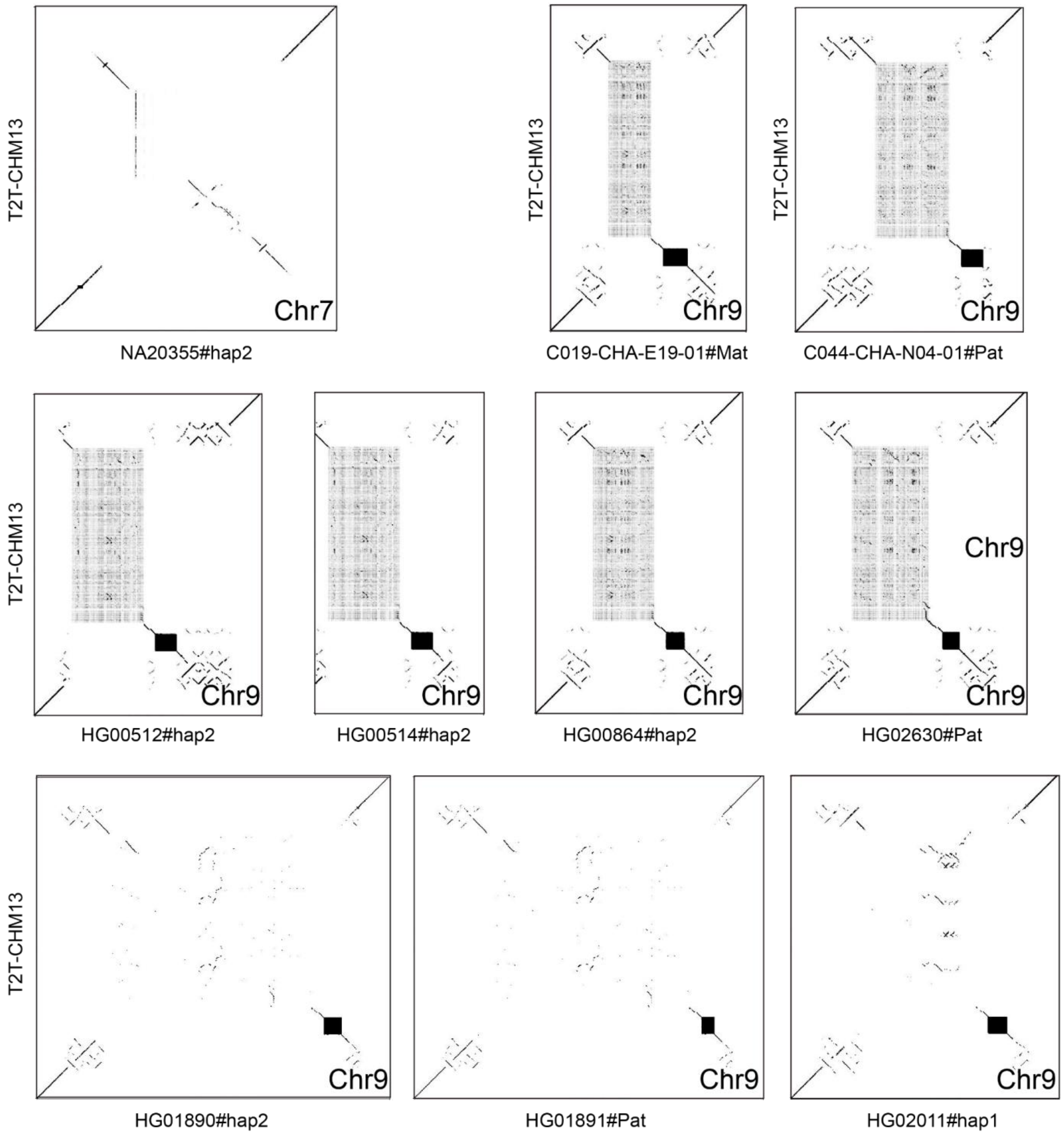


**Supplementary Fig. 19 | Alignment dotplots between the large-inversion carrier haplotype assembly and T2T-CHM13.** The plots were generated by gepard using a word length of 200.

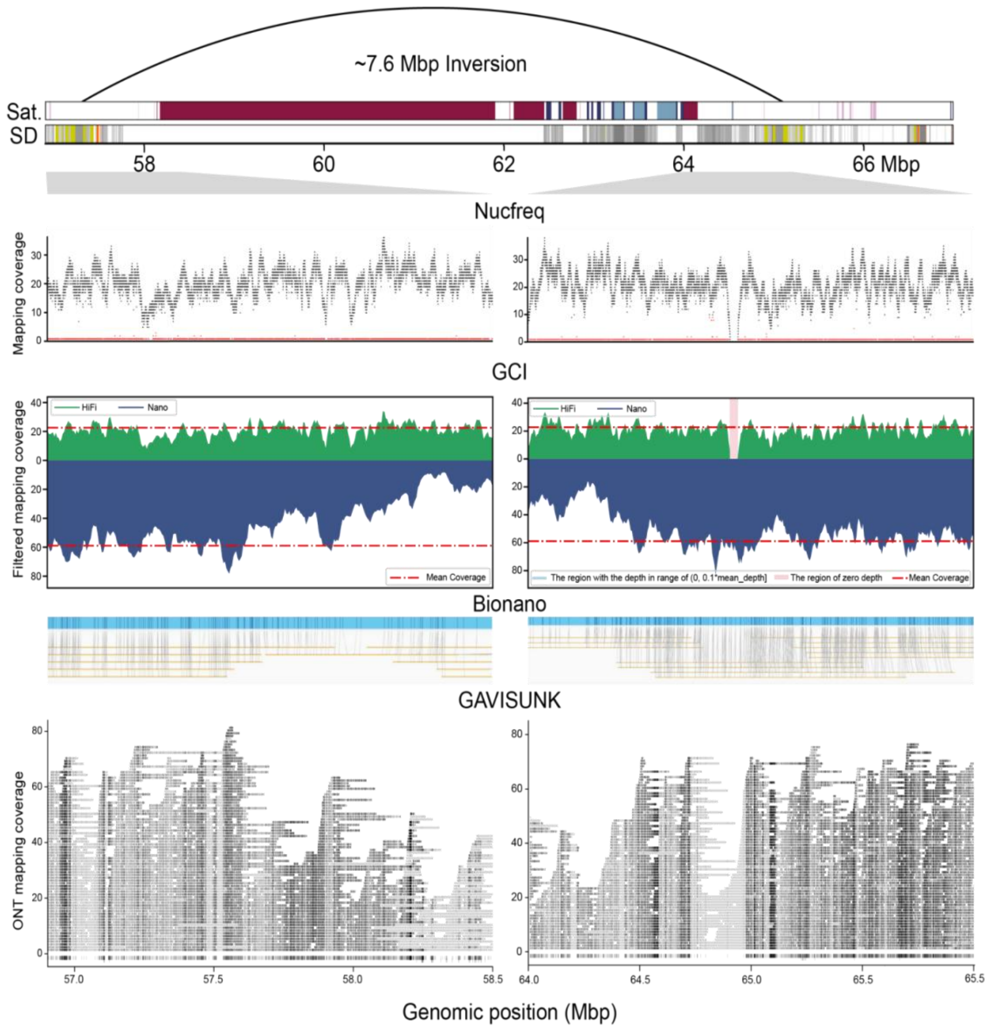


**Supplementary Fig. 20 | Validation of the local assembly around the pericentric inversion breakpoints in CEN07.** The top two tracks show the satellite repeat annotation and segmental duplication (SD) annotation of CEN07 in C037-CHA-S17-01#Mat. The Nucfreq plots demonstrate the PacBio HiFi mapping coverage at both inversion breakpoints. The GCI plots show the PacBio HiFi and ONT read coverage around the inversion breakpoints with stringent alignment filtration. The Bionano plots illustrate correspondence between markers at inversion breakpoints and several ONT reads across the breakpoints. The GAVISUNK plots show the singly unique nucleotide *k*-mers (SUNKs) at the inversion breakpoints and the concordant SUNK in the ONT reads.

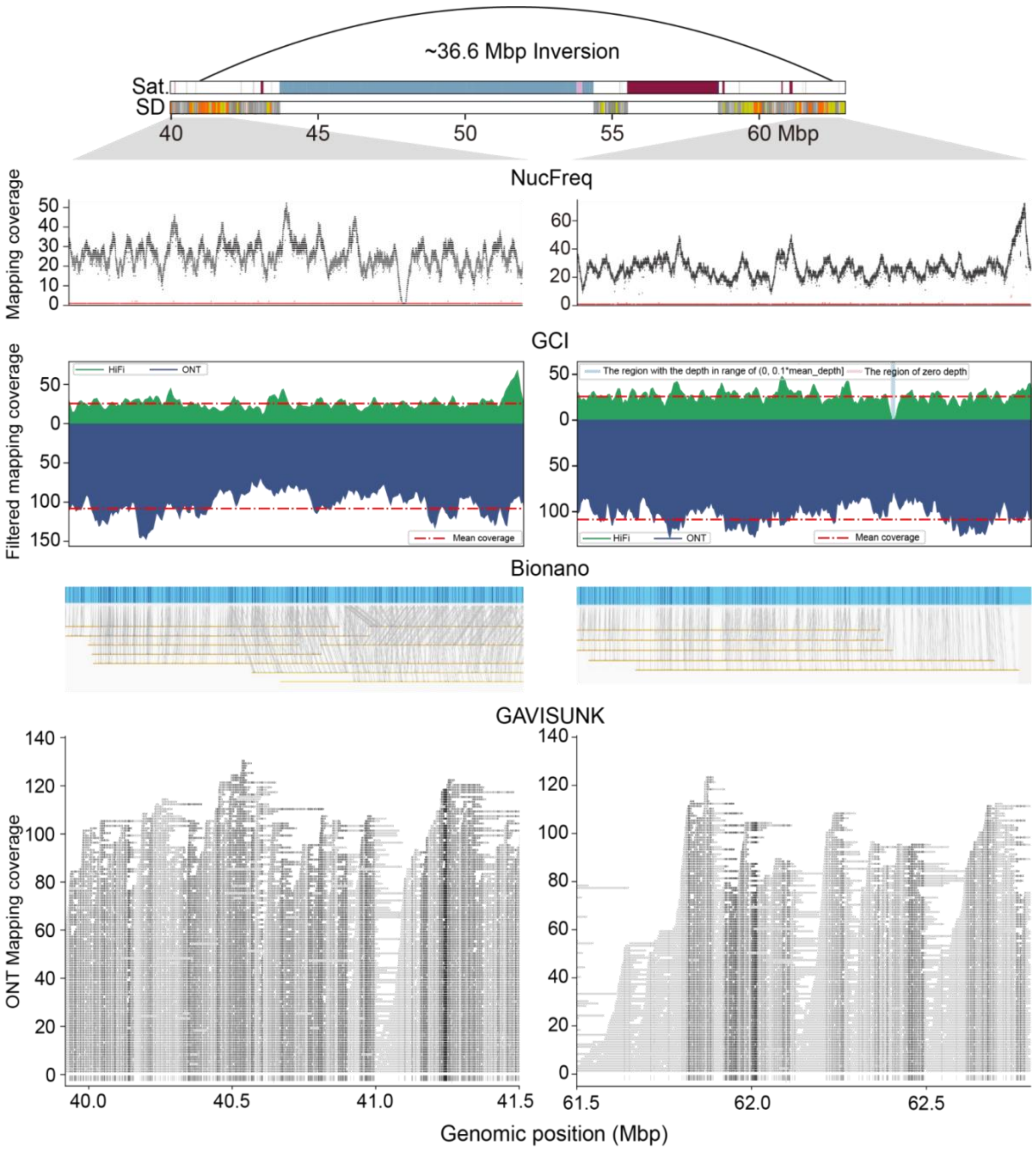


**Supplementary Fig. 21 | Validation of the local assembly around the pericentric inversion breakpoints in CEN09.** The top two tracks show the satellite repeat annotation and segmental duplication (SD) annotation of CEN09 in C051-CHA-N11-01#Mat.The Nucfreq plots demonstrate the PacBio HiFi mapping coverage at both inversion breakpoints. The GCI shows the PacBio HiFi and ONT read coverage around the inversion breakpoints with stringent alignment filtration. The Bionano plots illustrate consistency between markers at the inversion breakpoints and several ONT reads across the breakpoints. The GAVISUNK plots show the SUNKs at the inversion breakpoints and the concordant SUNKs on ONT reads.

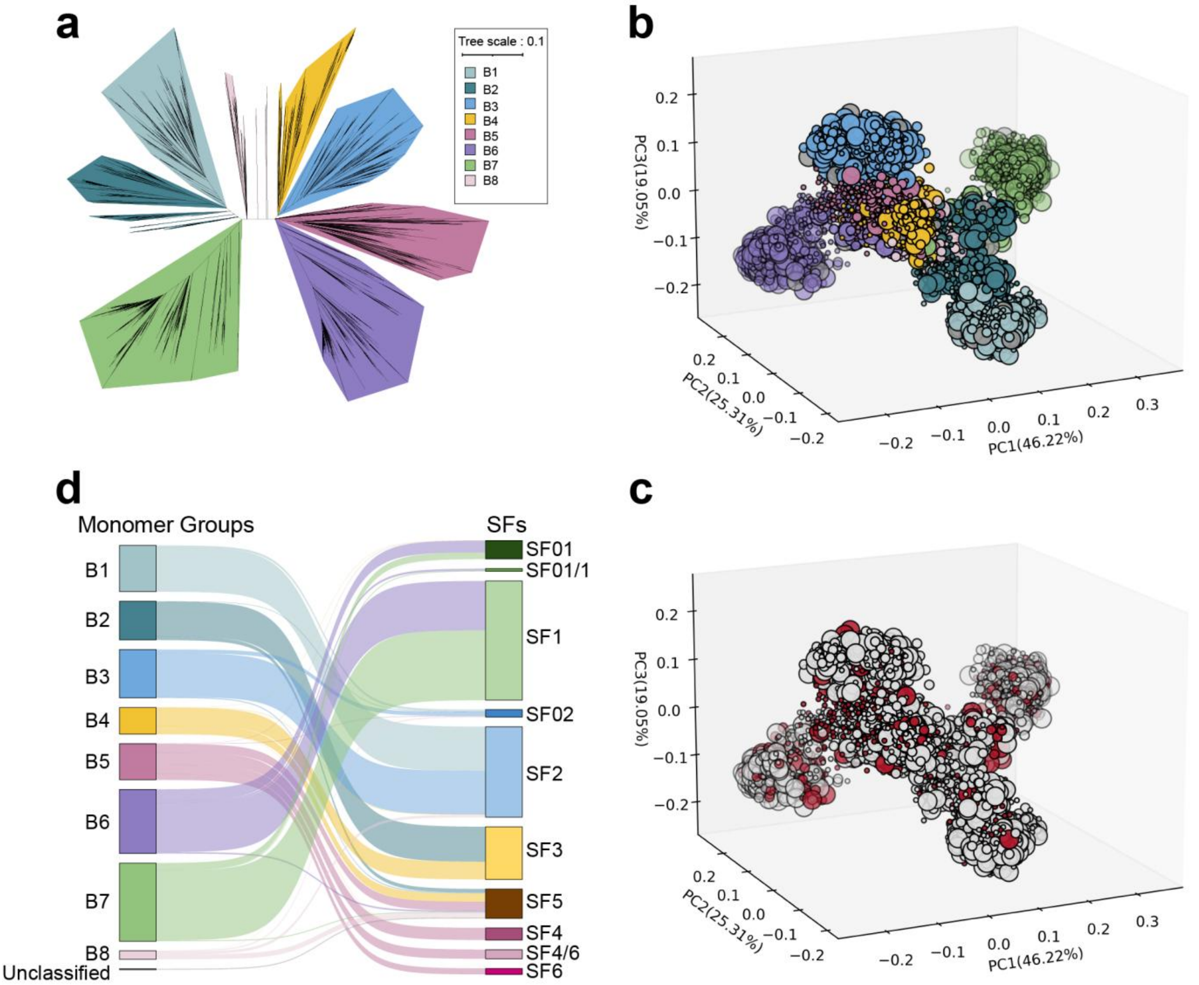


**Supplementary Fig. 22 | Phylogenetic classification of global human αSat HOR monomers. a**, A maximum-likelihood phylogenetic tree of representative αSat sequences from 3,722 HOR-organized monomer clusters, revealing eight distinct phylogenetic groups (B1-B8). **b**, Three-dimensional Principal Component Analysis (3D PCA) plot of 3,722 monomer clusters, colored by eight phylogenetic groups. **c**, 3D PCA plot highlighting the clusters (red) absent to the T2T-CHM13 reference assembly in red. **d**, Concordance between the eight phylogenetically defined groups and the established monomer suprachromosomal family (SF) category.

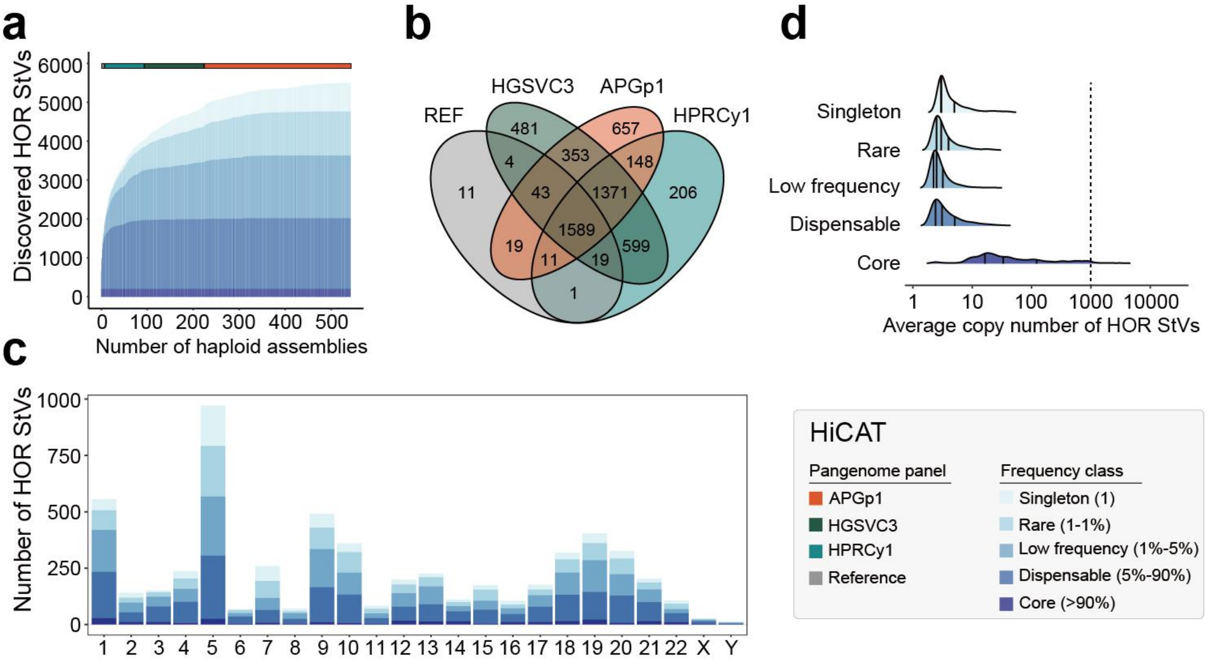


**Supplementary Fig. 23 | Distribution and abundance of HiCAT-based HOR StVs. a**, Growth curve of HiCAT-based HOR StVswith increasing numbers of haploid assemblies. Assemblies from APGp1, HPRCy1, HGSVC3, and reference-grade assemblies are included. HOR StVs are classified into five categories based on population frequency: core (>90%), dispensable (>5% & <=90%), low-frequency (>1% & <=5%), rare (>1 & <=1%), and singleton (*n* = 1). Reference-level assemblies include T2T-CHM13, T2T-CN1, Q100-HG002, and YAO. **b**, Venn diagram of HOR StVs identified in the APGp1, HPRCy1, HGSVC3, and reference panels, showing 657 StVs uniquely identified in APGp1. **c**, Distribution of HOR StVs categorized by population allele frequency across all chromosomes. **d**, Mean copy number of HOR StVs within each allele frequency category.

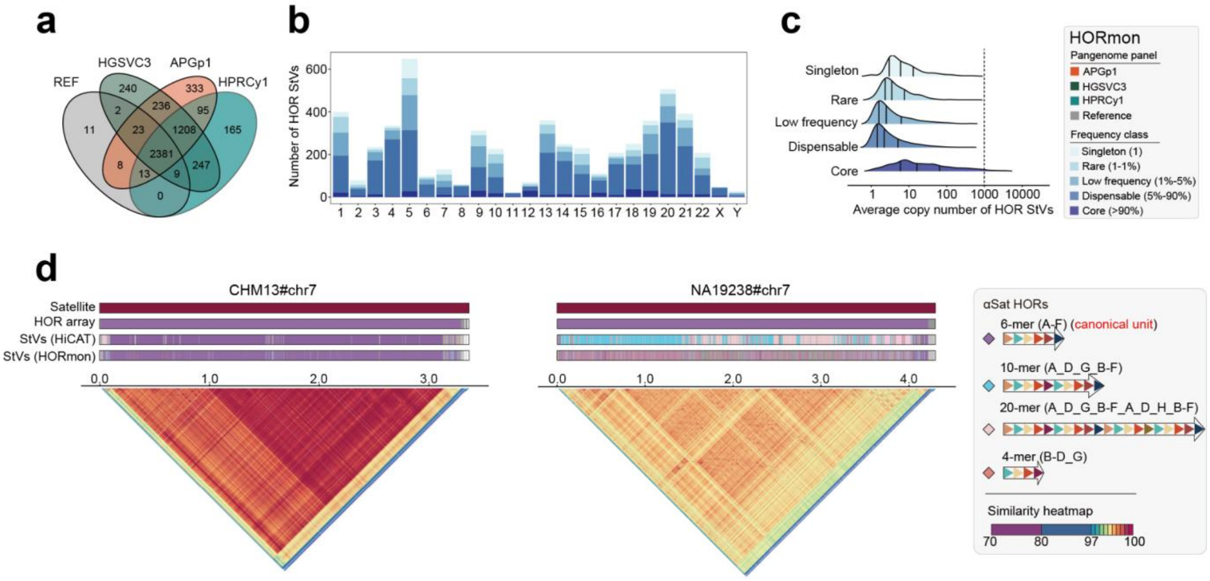

**Supplementary Fig. 24 | Distribution and abundance of HOR StVs across population allele frequencies identified by HORmon. a**, Venn diagram of HOR StVs in APGp1, HPRCy1, HGSVC3, and reference panels, showing 333 StVs uniquely identified in APGp1. **b**, Distribution of HOR StVs categorized by population allele frequency across all chromosomes. **c**, Mean copy number of HOR StVs within each allele frequency category. **d**, Individual-specific expansion of a 4-mer satellite variant in active arrays of CEN07. A 4-mer StV nested within the canonical 6-mer HOR array underwent a massive (>1,300 repeats) expansion in an AFR assembly (NA19238#hap2). Major HOR StVs and monomers are denoted by letters and colors to reflect distinct structural features.

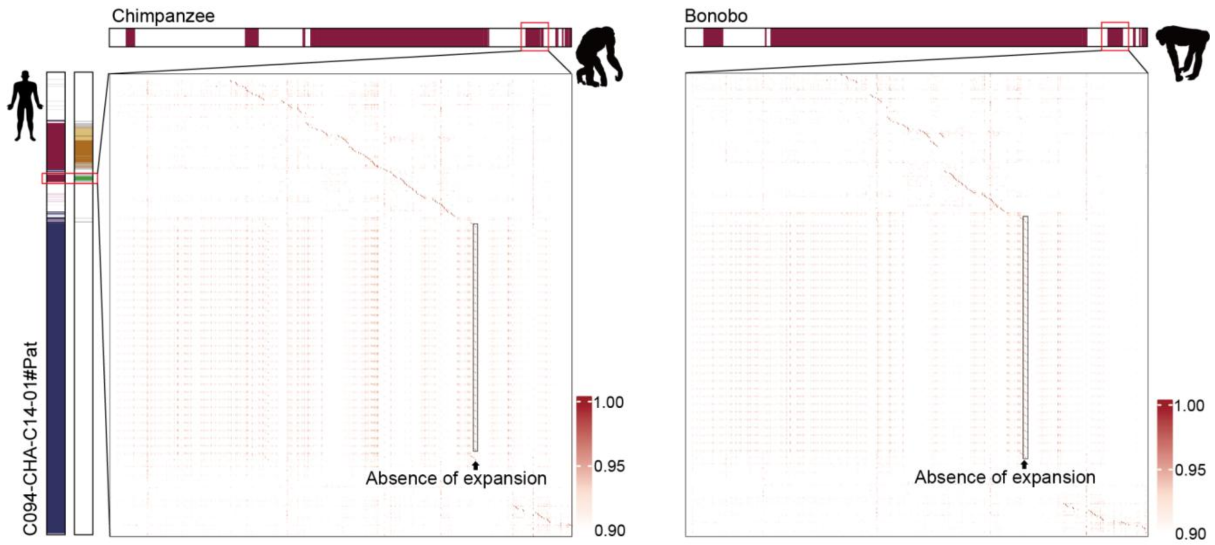


**Supplementary Fig. 25 | Sequence alignments of centromeric sequences around a newly discovered human HOR array in CEN16, between human and non-human primates (chimpanzee and bonobo).** The heatmaps visualize pairwise identity matrices calculated from monomer-level alignments of syntenic centromeric regions.

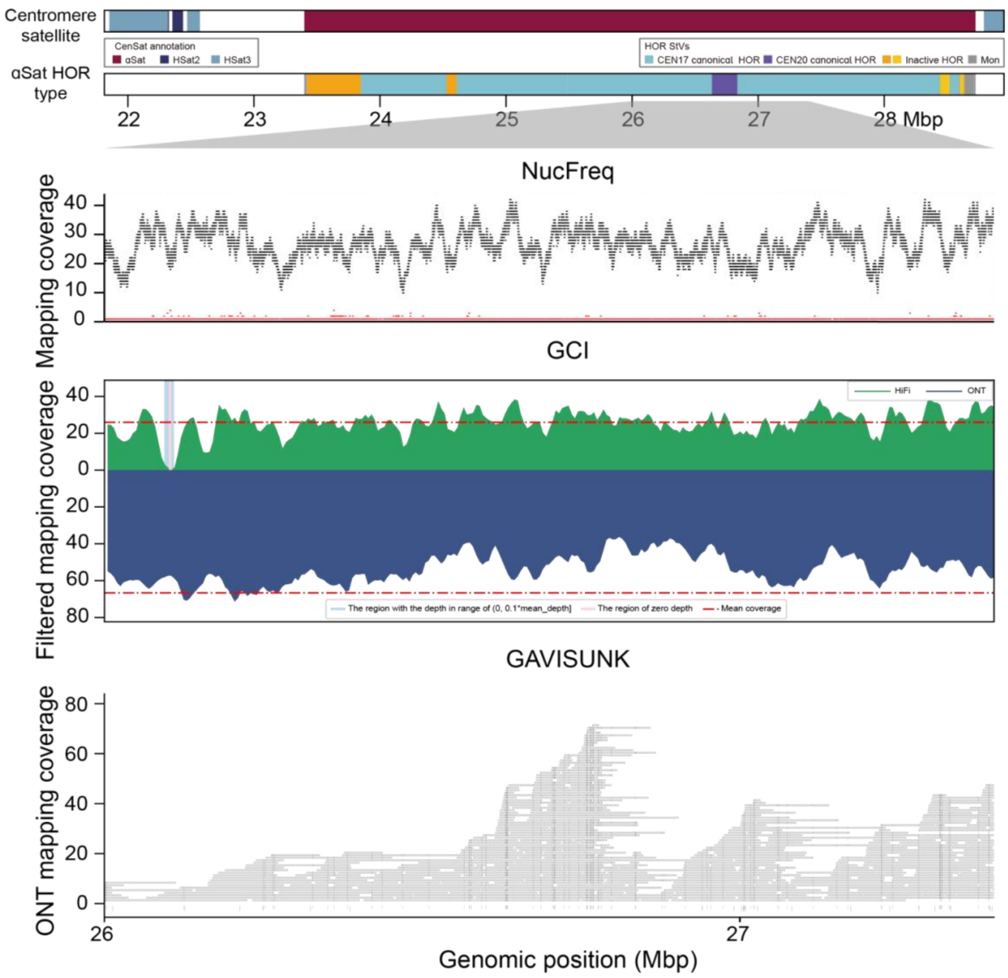


**Supplementary Fig. 26 | Validation of local assembly at a rare interchromosomal translocation of a canonical HOR array from CEN20 to CEN17.** Nucfreq plot demonstrates the PacBio HiFi mapping coverage across both the canonical CEN20 HOR array insertion and its flanking regions in C148-CTJ01-01. GCI plot shows PacBio HiFi and ONT mapping coverage with stringent alignment filtration and GAVISUNK plots show the SUNKs across the region and the concordant SUNKs in ONT reads.

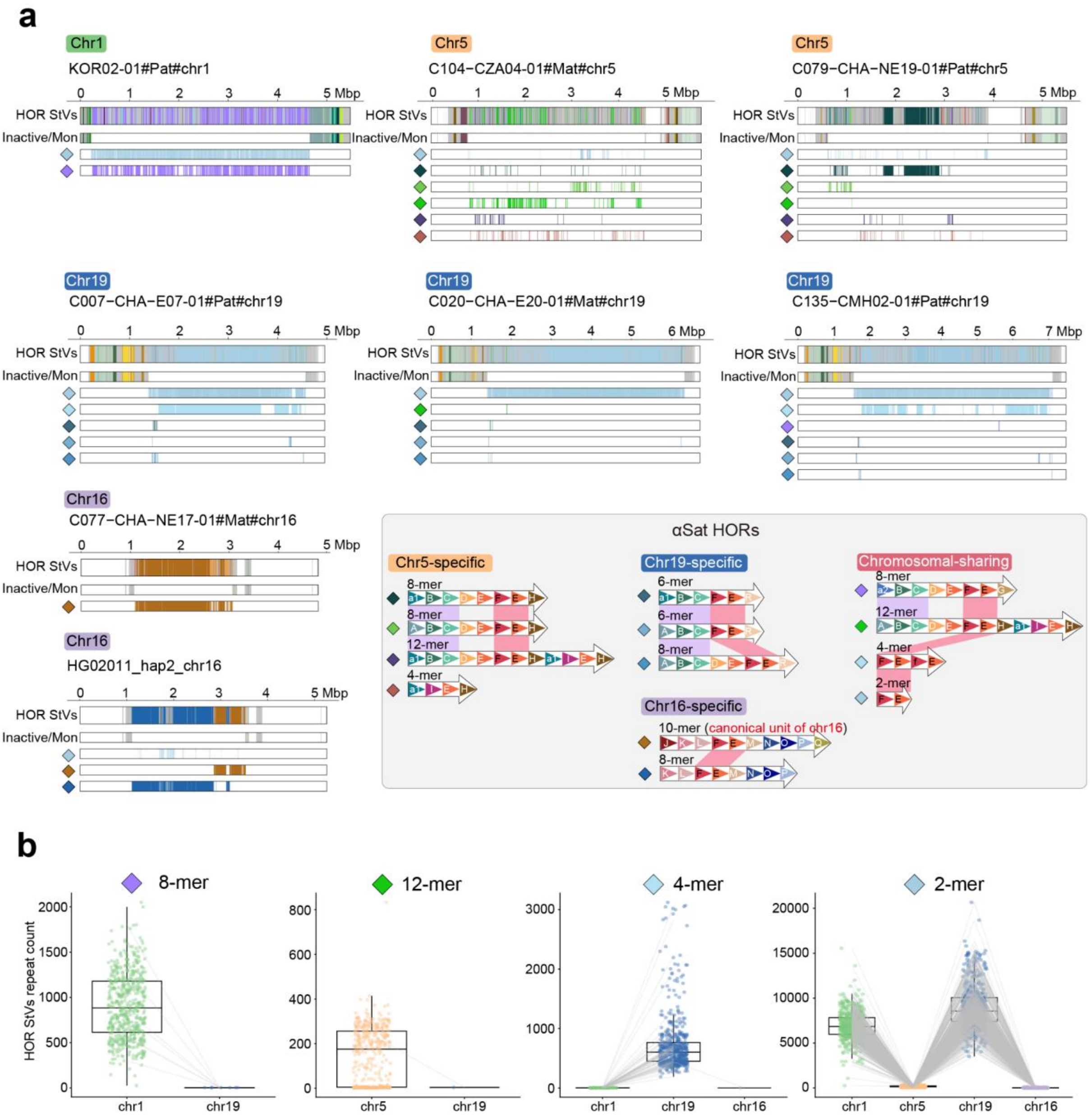


**Supplementary Fig. 27 | Divergence and chromosome-specific amplification of shared HOR StVs between CEN01/05/19/16. a**, Architecture of major HOR StVs on chromosomes 1, 5, 16 and 19. Representative examples are shown for each chromosome, with different StVs displayed in separate tracks. Distinct colors and letters denote specific monomer compositions and structures of HORs. Shared segments are highlighted by purple lines aligning the A-B-C monomers common to chr1, 5 and 19, and red lines aligning the F-E monomers universal to all four chromosomes. **b**, Copy number variation of the inter-chromosomal shared StVs across haplotypes, revealing divergent, chromosome-specific amplification patterns.

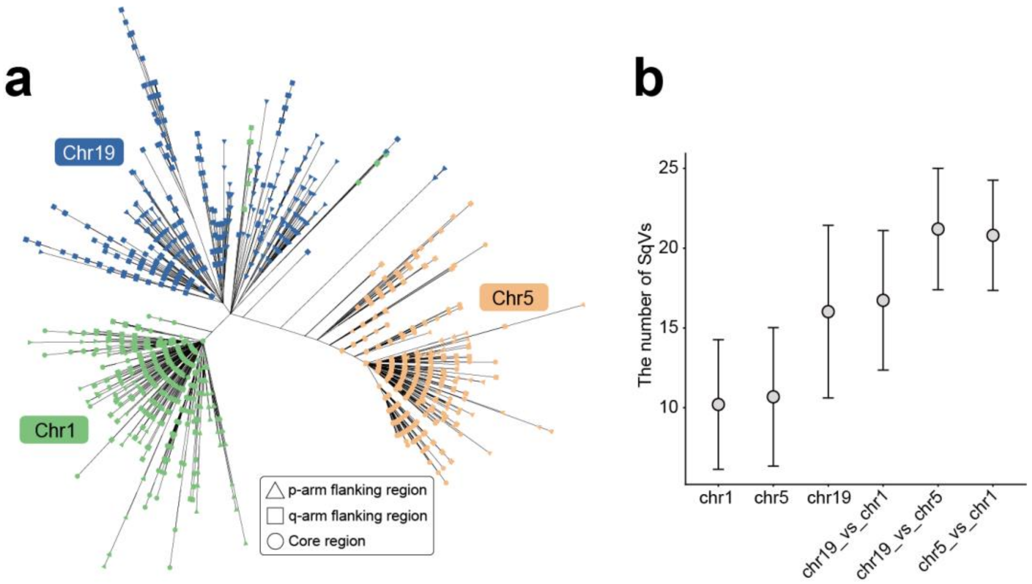


**Supplementary Fig. 28 | Divergence of partially shared HOR sequences (hor_A-C) among CEN01/05/19. a**, Phylogeny of partially shared sequence (hor_A-C) among CEN01/05/19. **b**, Pairwise alignments of 1,000 randomly selected sequences per chromosome revealed that chromosome 19 exhibits the highest level of sequence polymorphism.

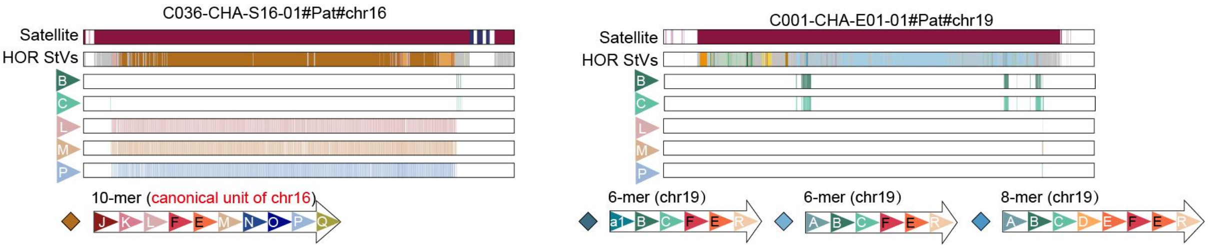


**Supplementary Fig. 29 | Shared monomers in the flanking regions of active arrays specific to CEN19 and CEN16.** Flanking regions of CEN19 and CEN16 active arrays bidirectionally incorporate core monomers from each other's HORs (B and C from CEN19 around CEN16; L, M, and P from CEN16 around CEN19). Within the framework of a 'layered expansion' model, this pattern suggests a shared ancestry between CEN16 and CEN19.

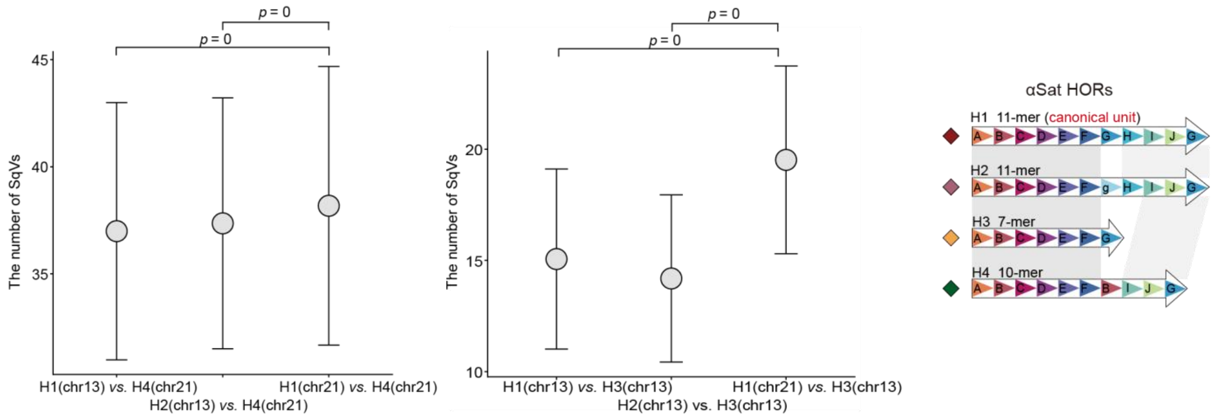


**Supplementary Fig. 30 | Reduced SNP divergence of shared StVs between CEN21 and CEN13.** H1, H2, H3, and H4 denote the dominant HOR StVs in CEN13 and CEN21, represented as follows: H1 corresponds to the canonical 11-mer HOR StV (A–J_G), H2 to an 11-mer HOR variant (A–F_g_H–J_G), H3 to a 7-mer HOR variant (A–G), and H4 to a 10-mer HOR variant (A–F_B_I–J_G). The circle represents the mean value, and the bar indicates the mean ± standard deviation. The CEN21 10-mer HOR sequence (H4) is significantly closer to CEN13 11-mer HOR H1/H2 (SNP divergence: 37.0 ± 5.8) than to the CEN21 11-mer HOR H1 (38.2 ± 6.5), suggesting an inter-chromosomal transfer from CEN13 (Wilcoxon rank-sum test). In contrast, the CEN13 7-mer is most closely related to the CEN13 11-mer, confirming a local, intra-chromosomal origin.

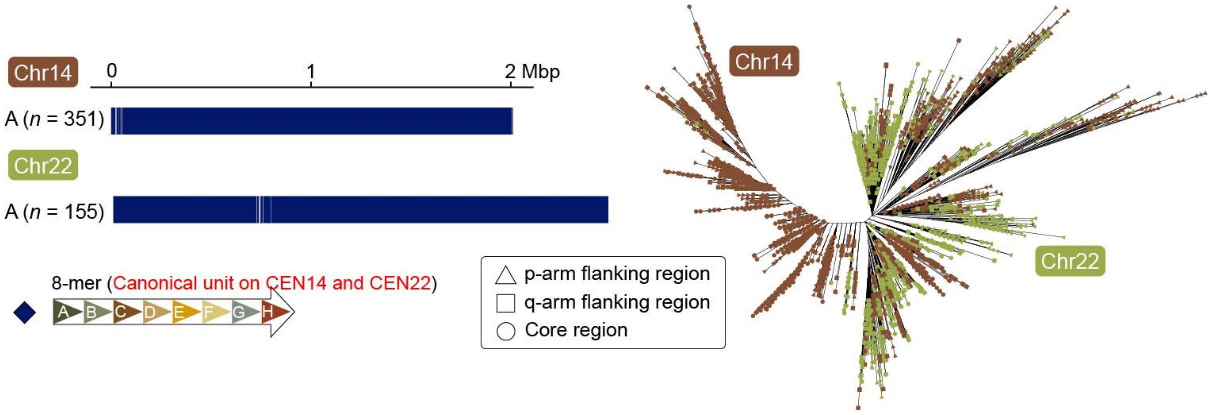


**Supplementary Fig. 31 | Population phylogeny of canonical 8-mer HORs reveals shared variation between CEN14 and CEN22.** A maximum-likelihood phylogeny, based on major CenHaps and dominant StVs, shows that canonical 8-mer HORs from CEN14 and CEN22 are intermixed and cannot be distinguished as separate chromosomal clusters. This differs from the distinctness of CEN14 and CEN22 HOR clusters reported in the T2T-CHM13 assembly, indicating potential recurrent inter-chromosomal exchange between these chromosomes.

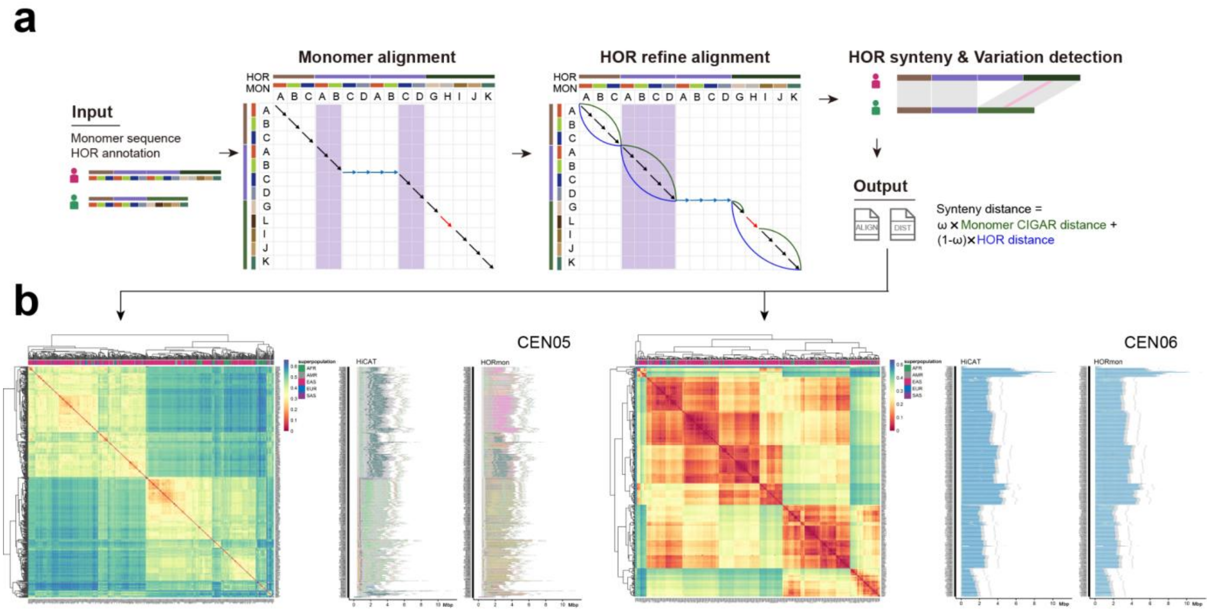


**Supplementary Fig. 32 | Centromeric haplotyping based on synteny distance. a**, Schematic of the HORSCAN pipeline, which consists of monomer alignment, HOR refinement and variant detection modules, and outputs synteny distances and pairwise alignment files. The synteny distance integrates two components: the monomer-level CIGAR distance, which measures the proportion of non-matching positions (mismatches and indels) relative to all aligned monomer pairs, and the HOR distance, which quantifies structural divergence by calculating the Jaccard distance between each matching pair of HOR StVs (see **Methods**). **b**, Variation in synteny distance is mainly driven by two distinct factors: the CIGAR component, which captures differences in HOR array size (e.g., CEN06), and the HOR component, which reflects structural variants within HORs (StVs; e.g., CEN05). These patterns reveal two modes of centromeric haplotype differentiation.

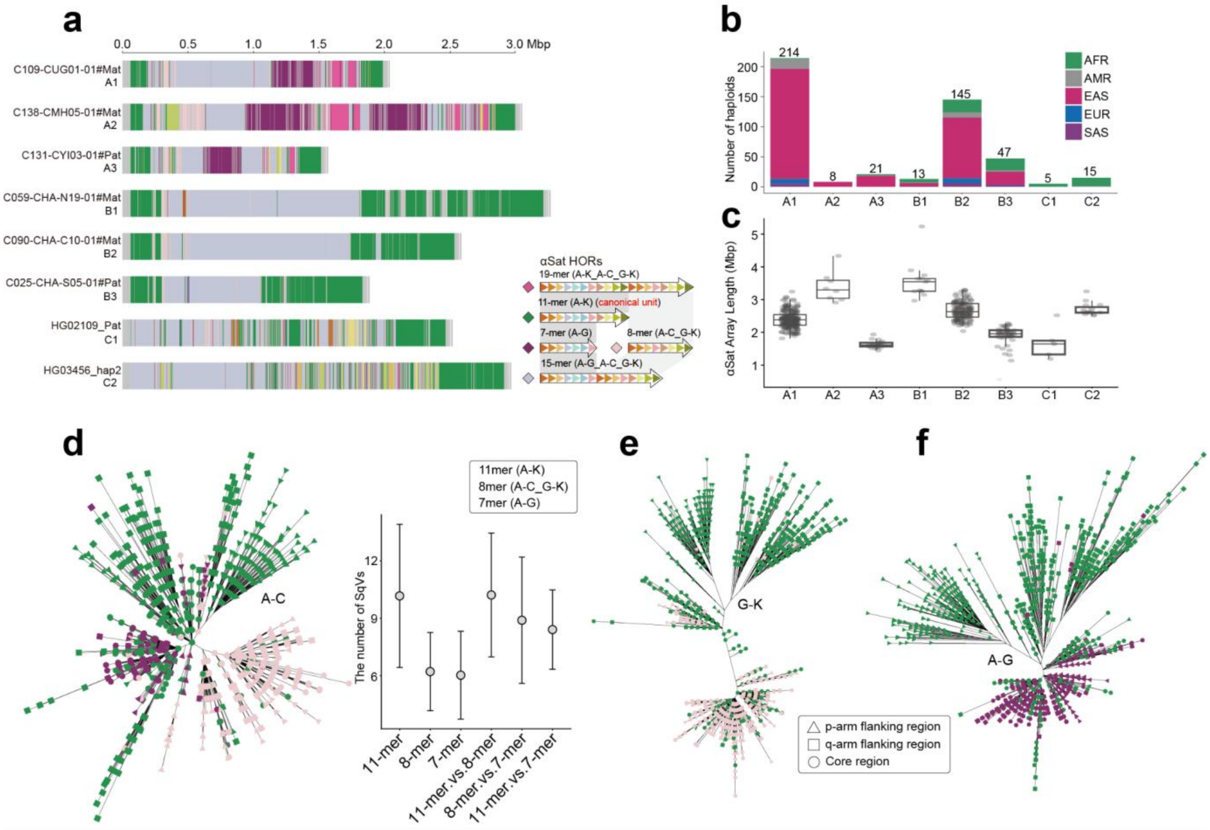

**Supplementary Fig. 33 | Structural composition and superpopulation distribution of three major CenHaps of CEN08. a**, Representative structures of three major CenHaps in CEN08, with highlighted major HOR StVs. **b**, Stacked bar plot showing the distribution of CenHaps across superpopulations. CenHap-A and its subtypes (A1–A3) predominated in EAS superpopulation, while CenHap-B and CenHap-C, particularly subclades B2, B3 and C2, were more common in AFR, reflecting distinct demographic histories. The total number of carriers (*n*) for each CenHap is shown above the bars. **c**, Distribution of αSat array lengths across CenHaps. The center line represents the median, the box limits indicate the upper and lower quartiles (25th and 75th percentiles), and the whiskers extend to 1.5 times the interquartile range from the box. **d**, **e** and **f**, Phylogeny of partial shared HOR sequences (A-C, G-K and A-G) and revealed evolutionary trajectory of HOR StVs. Pairwise alignments of 1,000 randomly selected sequences for each StV show that the H1 (11-mer) is the most polymorphic unit compared to H2 (8-mer) and H3 (7-mer). The circle represents the mean value, and the bar indicates the mean ± standard deviation. Collectively, the 11-mer represents the ancestral configuration, with the subsequent emergence of 7-mer and 8-mer variants, supporting the layered expansion model.

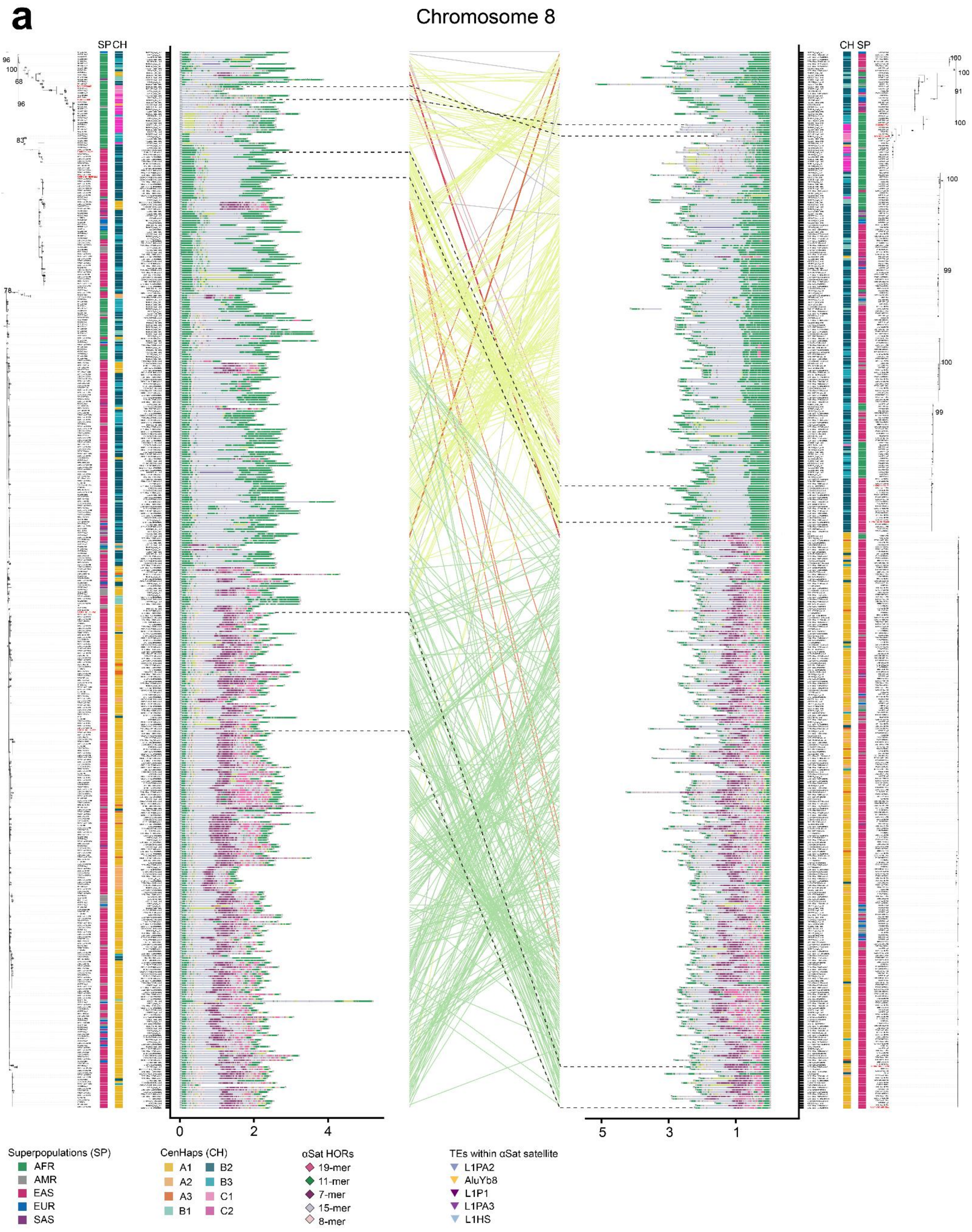
a
Chromosome 8
SP CH
CH SP
0
2
4
5
3
1
Superpopulations (SP)
AFR
AMR
EAS
EUR
SAS
CenHaps (CH)
A1
A2
A3
B1
B2
B3
C1
C2
αSat HORs
19-mer
11-mer
7-mer
15-mer
8-mer
TEs within αSat satellite
L1PA2
AluYb8
L1P1
L1PA3
L1HS

**b**

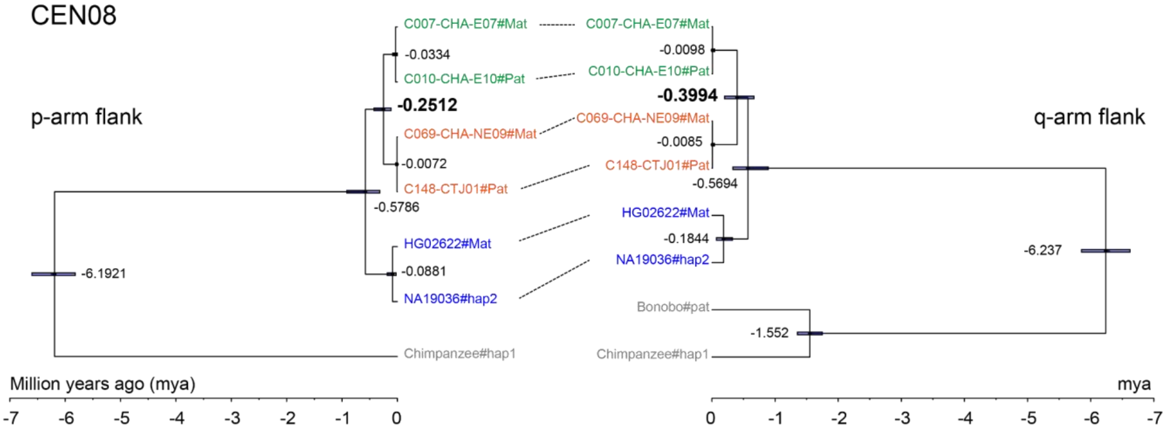


**Supplementary Fig. 34 | Phylogeny congruence of CEN08 centromeric flanks. a**, Maximum-likelihood phylogenetic trees of the p- and q-arm of CEN08 reveal congruent topologies among major CenHaps. Bootstrap values from 60 to 100 for key nodes are indicated. Annotation tracks display the superpopulation (SP) and CenHaps (CH). Corresponding leaves between the p- and q-arm trees are connected by colored lines. Assemblies used for divergence time estimations are highlighted in red and linked with dashed lines. **b**, Divergence time estimation for major CenHaps of CEN08, based on 20 Kbp flanking sequences from the p-arm and 30 Kbp from the q-arm, using chimpanzee and bonobo as outgroups. Bonobo was excluded due to lack of synteny with the human genome in this region. The q-arm region was extended to 30 Kbp to obtain a sufficient number of SNPs for reliable inference.

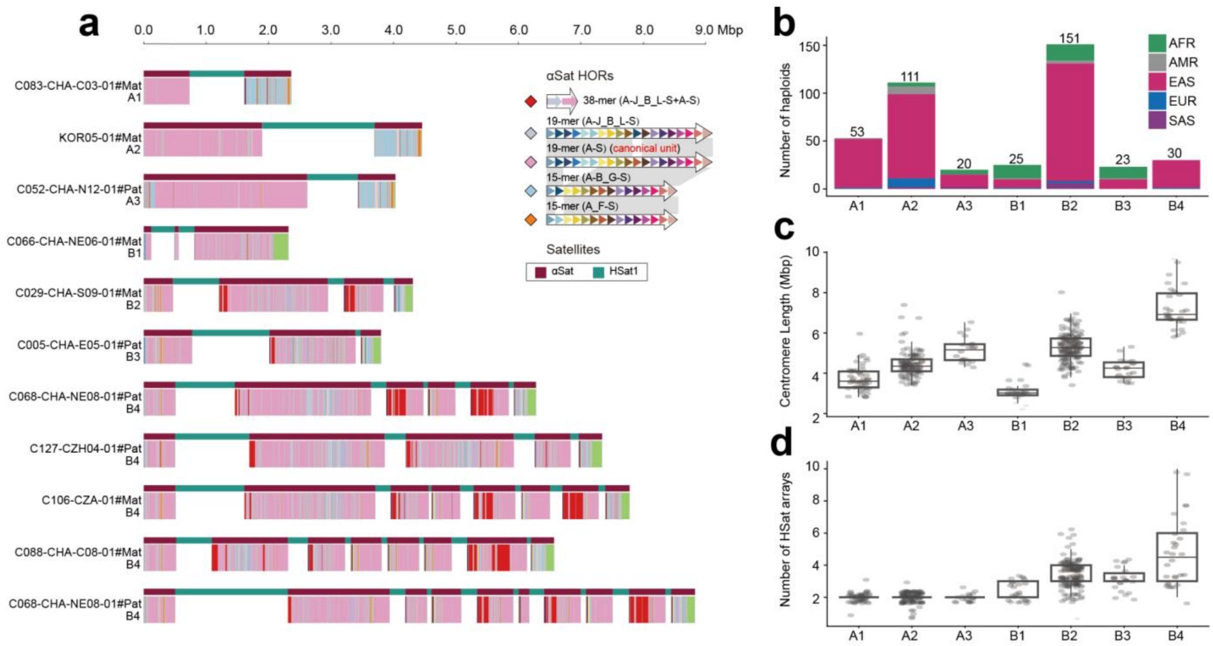


**Supplementary Fig. 35 | Structural composition and superpopulation distribution of three major CenHaps of CEN04. a**, Representative structures of two major centromere haplotypes (CenHap-A and CenHap-B) and their subtypes on CEN04, as defined by hierarchical clustering based on a pairwise synteny distance matrix. Satellite tracks are shown; blank regions in the HOR StVs track correspond to the HSat1 array. **b**, CenHap distribution across superpopulations. **c**, Centromere lengths for CenHaps. **d**, Distribution of αSat-HSat1A interleaved structures across CenHaps. CenHap-B is strongly associated with a greater number of HSat1A block insertions. The EAS-enriched subcluster B4 exhibits frequent HSat1A insertions with larger size, suggesting population-specific duplication events.

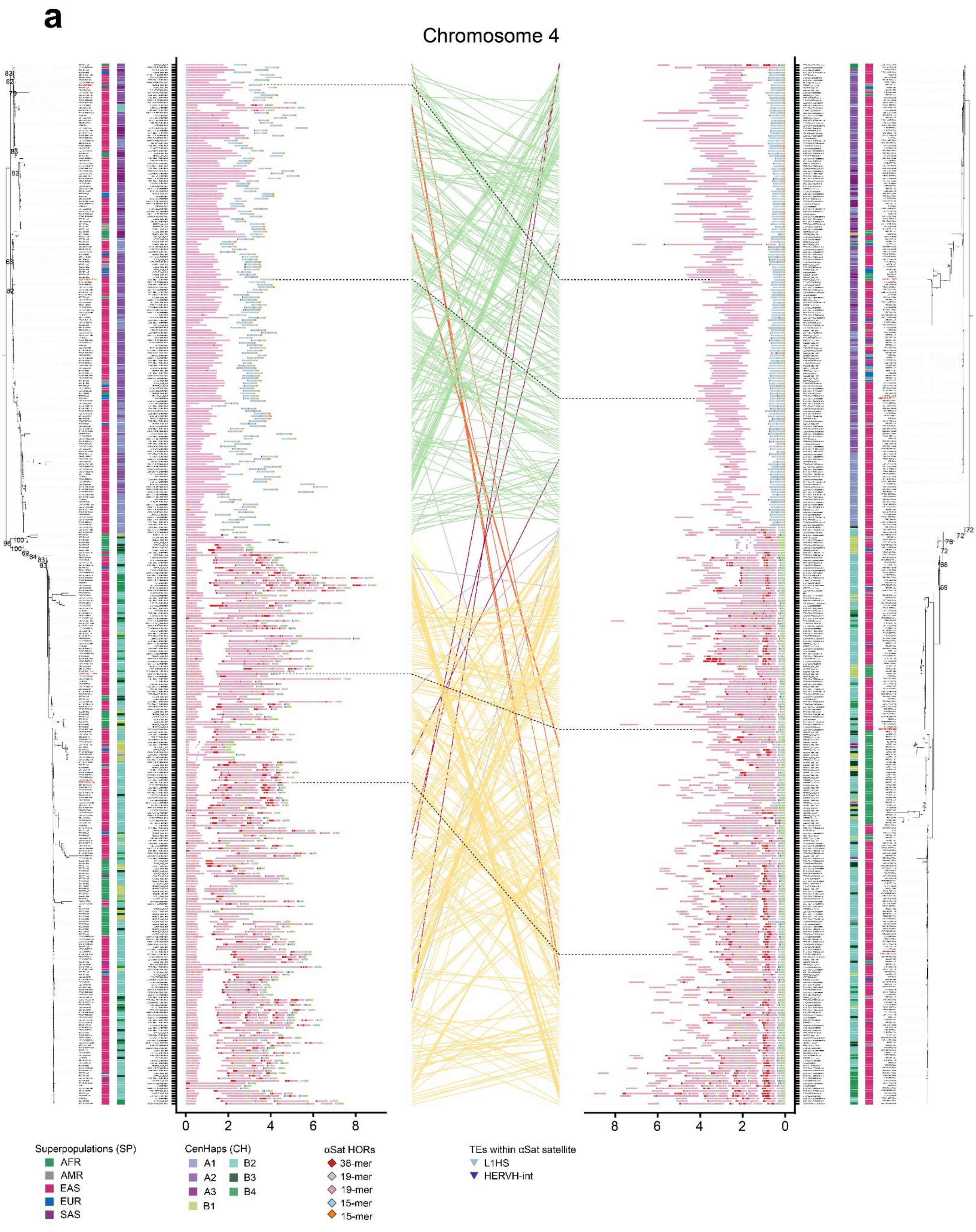
a
Chromosome 4
Superpopulations (SP)
AFR
AMR
EAS
EUR
SAS
CenHaps (CH)
A1
A2
A3
B1
B2
B3
B4
αSat HORs
38-mer
19-mer
19-mer
15-mer
15-mer
TEs within αSat satellite
L1HS
HERVH-int

b

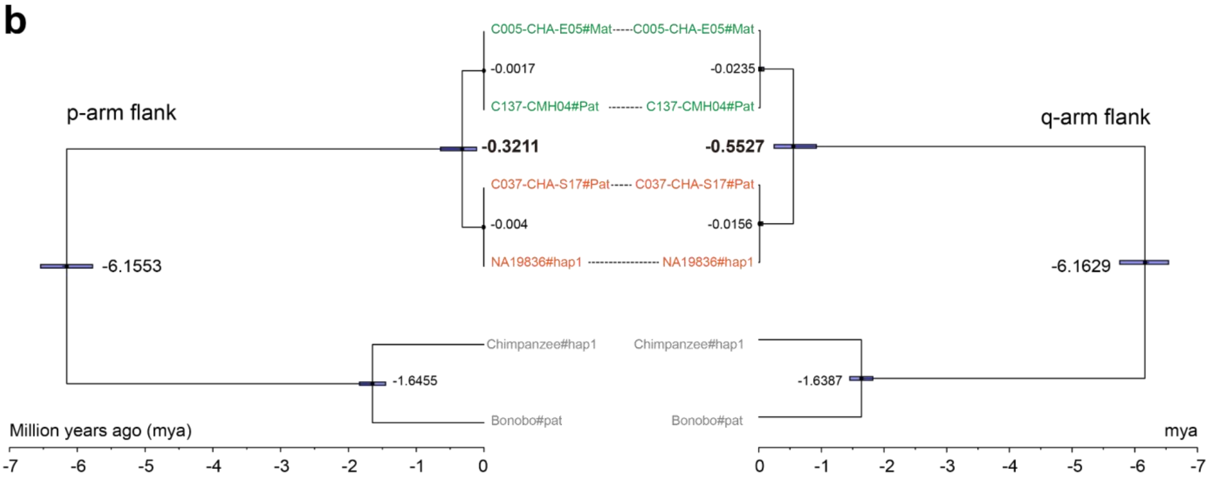


**Supplementary Fig. 36 | Phylogeny congruence of centromeric haplotypes of CEN04. a**, Maximum-likelihood phylogenetic trees for the p- and q-arms of CEN04 reveals congruent topologies among major CenHaps. Bootstrap support values (60%-100%) for key nodes are indicated. Annotation tracks display the superpopulation (SP) and CenHaps (CH). Corresponding leaves between the p- and q-arm trees are connected by colored lines. Assemblies used for divergence time estimations are highlighted in red and linked with dashed lines. **b**, Divergence time estimation for the two major CenHaps of CEN04, based on 20-Kbp flanking sequences of the p-arm and q-arm, using chimpanzee and bonobo as outgroups. The time estimation is performed by BEAST in the calibrated Yule model. Node ages are reported as posterior median estimates, and uncertainty is represented by 95% Highest Posterior Density intervals. The estimates (0.3211 and 0.5527 mya) of the key nodes between two CenHaps are highlighted.

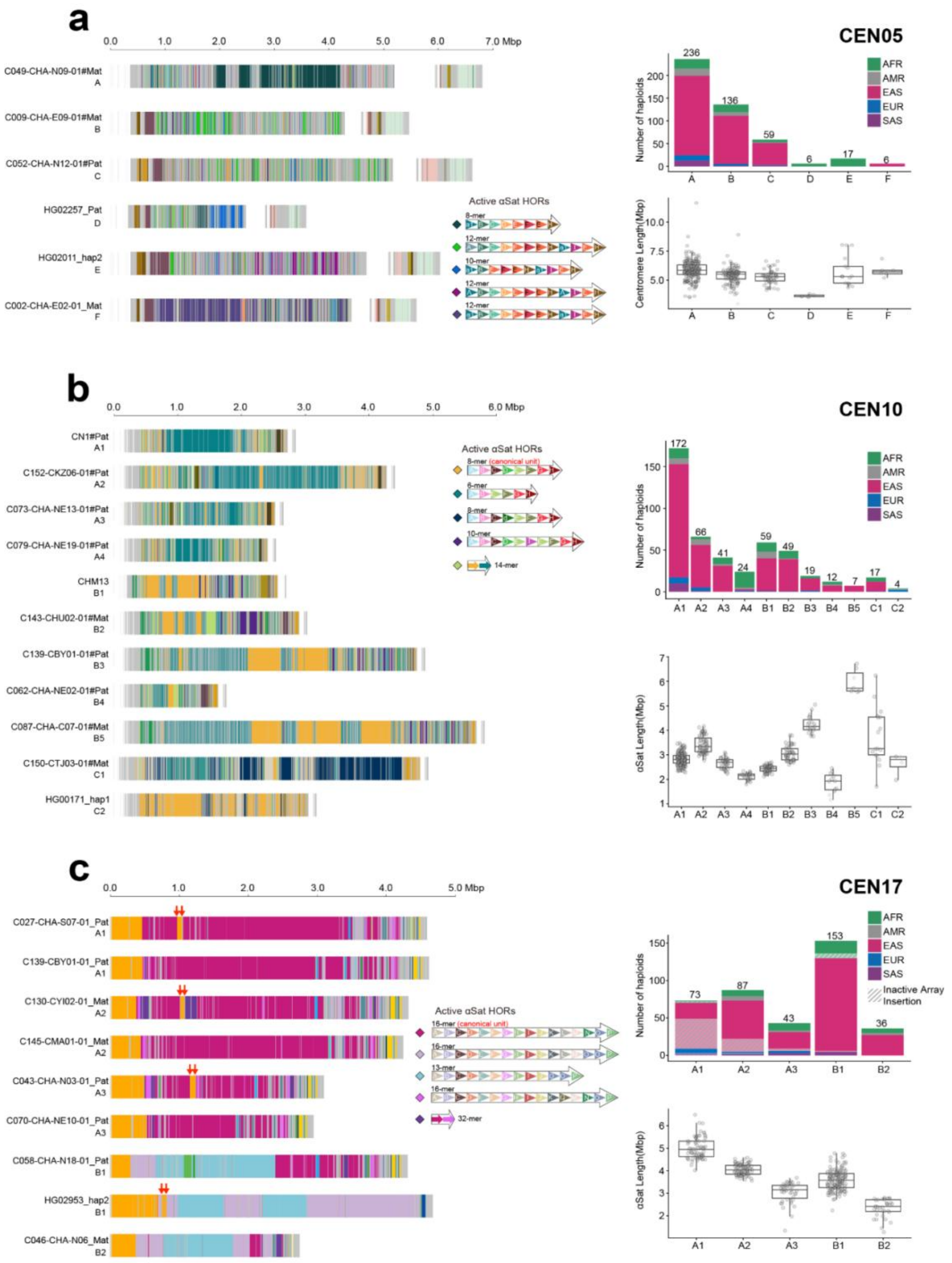


**Supplementary Fig. 37 | Structural composition and superpopulation distribution of major CenHaps in CEN05, CEN10 and CEN17.** Panels **a**–**c** depict representative structures with major HOR sequence variants (StVs), superpopulation distribution, and α-satellite array length for CEN05 (**a**), CEN10 (**b**), and CEN17 (**c**). Notably, in CEN17, insertions of inactive arrays that split the active array were observed, highlighted with red arrows, and were population-specific, being predominantly associated with EAS (CenHap-A) and with AFR (CenHap-B).

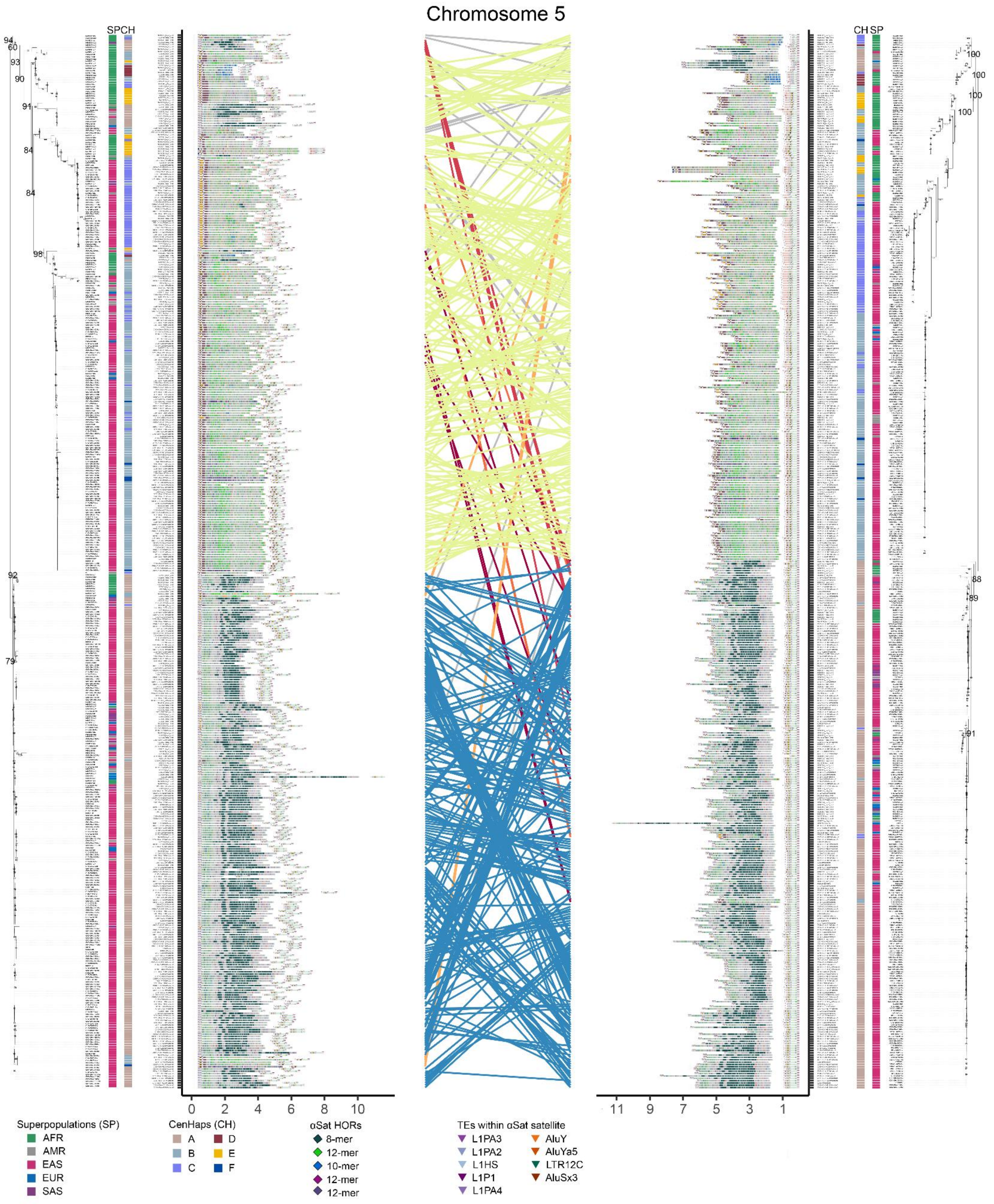


**Supplementary Fig. 38 | Phylogeny congruence of centromeric haplotypes of CEN05.** Phylogenetic analysis based on maximum likelihood for the p-arm and q-arm of CEN05 reveals congruent topologies among major CenHaps. Bootstrap support values (60%-100%) for key nodes are indicated. Annotation tracks record the superpopulation and CenHap assignments. Corresponding branches between the p- and q-arm trees are connected by colored lines.

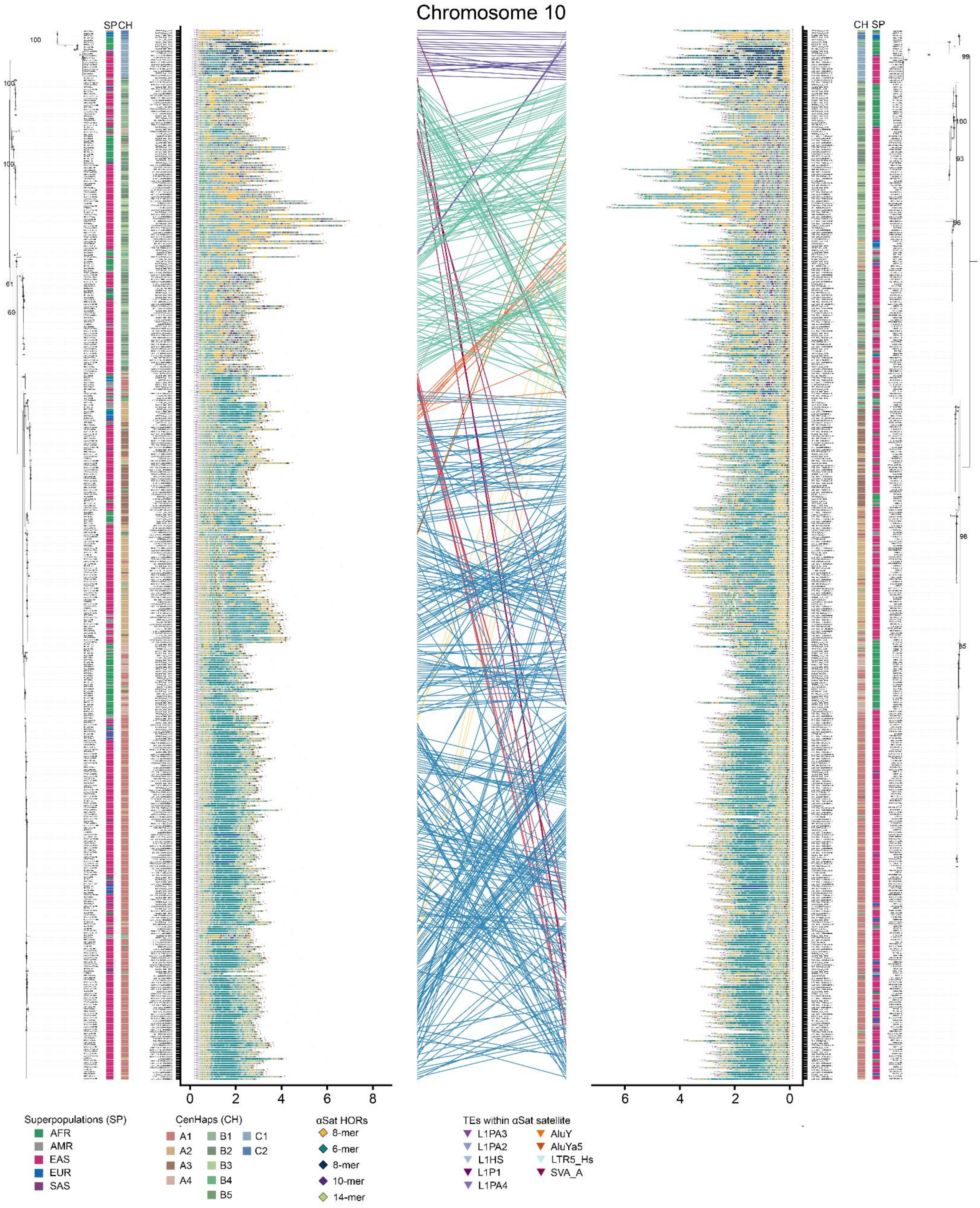


**Supplementary Fig. 39 | Phylogenetic reconstruction of centromeric haplotypes of CEN10.** Phylogenetic analysis based on maximum likelihood for the p- and q-arm of CEN10 reveals congruent topologies among major CenHaps. Bootstrap support values (60%-100%) for key nodes are indicated. Annotation tracks record the superpopulation and CenHap assignments. Corresponding branches between the p- and q-arm trees are connected by colored lines.

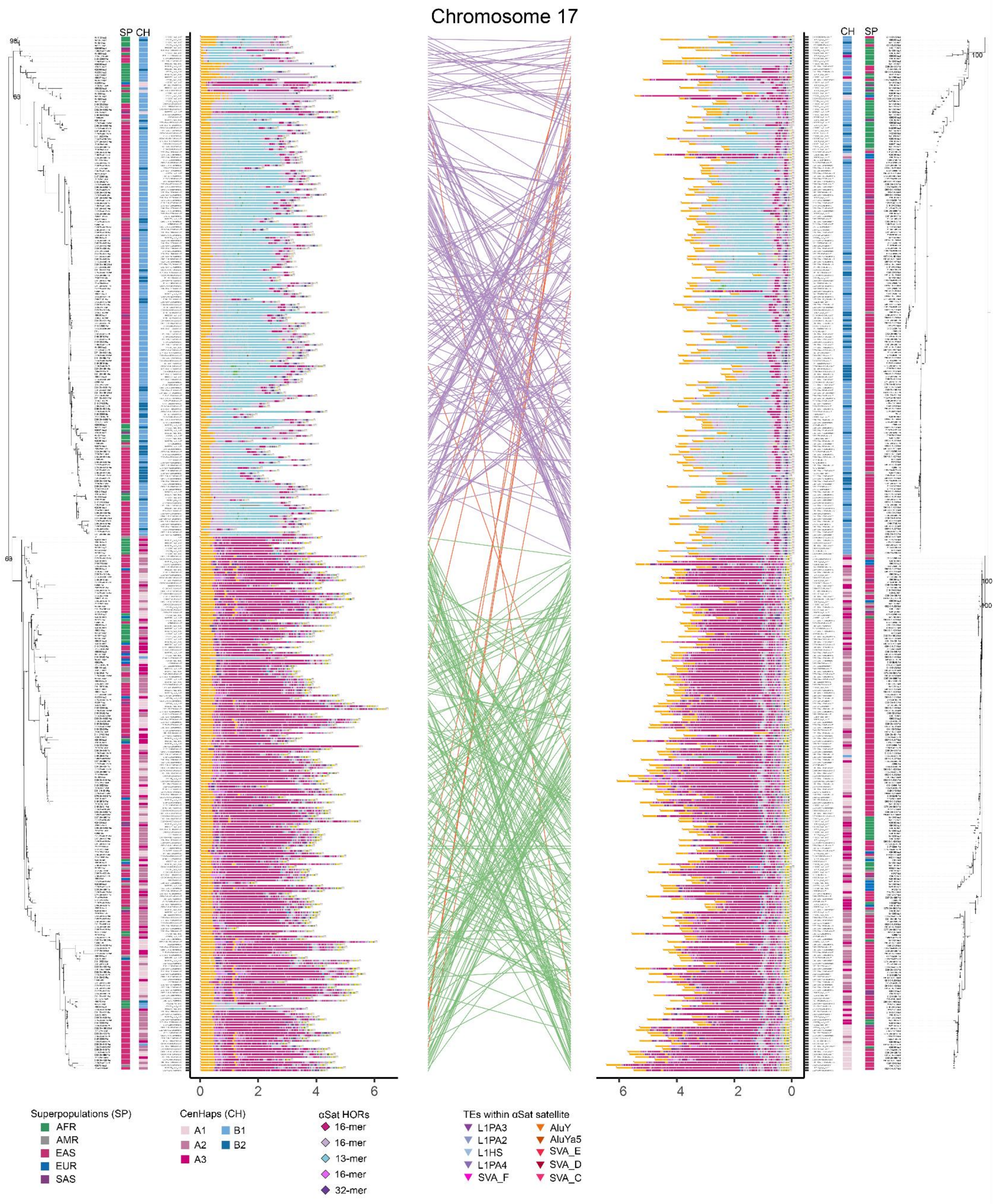


**Supplementary Fig. 40 | Phylogenetic reconstruction of centromeric haplotypes of CEN17.** Phylogenetic analysis based on maximum likelihood for the p- and q-arm of CEN17 reveals congruent topologies among major CenHaps. Bootstrap support values (60%-100%) for key nodes are indicated. Annotation tracks record the superpopulation and CenHap assignments. Corresponding branches between the p- and q-arm trees are connected by colored lines. Topological variations are restricted to terminal branches owing to limited branch lengths that impede reliable resolution; thus, they do not constitute robust evidence for global discordance. ka, thousand years ago.

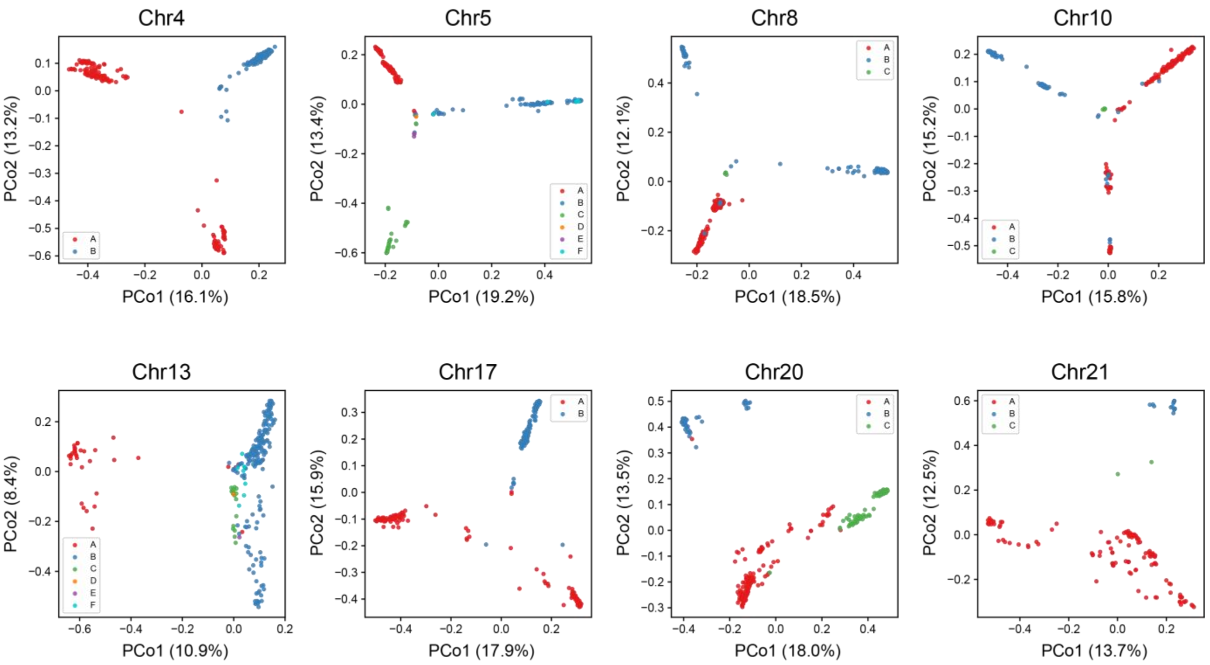


**Supplementary Fig. 41| Unsupervised Principal Coordinates Analysis (PCoA) of centromeric k-mer abundance profiles validating CenHap classification.** Individual panels illustrate PCoA clusters derived from pairwise Bray-Curtis distances, computed exclusively from centromeric αSat *k*-mer copy-number matrices (31-mers; population-informative *k*-mers at 2%-25% frequency). Each dot represents an individual haplotype assembly, projected along the first two principal coordinates (PCo1 and PCo2), with the percentage of explained variance annotated parenthetically. Assemblies are color-coded independently according to their CenHap assignments. The consistency between the *k*-mer clustering and CenHaps across the StV-driven chromosomes (4, 5, 8, 10, 13, 17, 20, and 21) demonstrates that the CenHap classification framework reliably captures discrete, genome-wide sequence-level divergence independently of heuristic clustering thresholds.

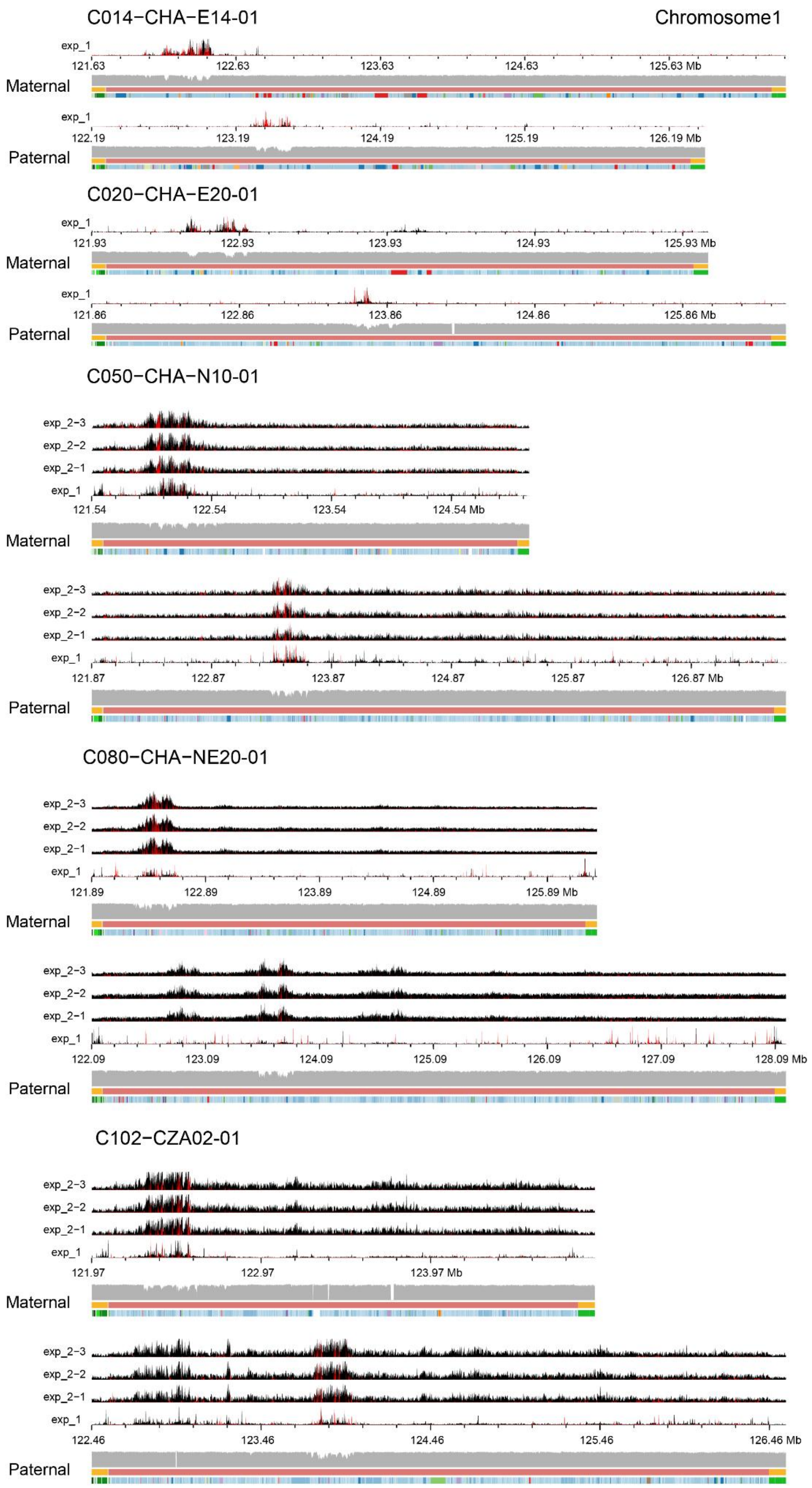

C014−CHA−E14-01
Chromosome1
exp_1
121.63
122.63
123.63
124.63
125.63 Mb
Maternal
exp_1
122.19
123.19
124.19
125.19
126.19 Mb
Paternal
C020−CHA−E20-01
exp_1
121.93
122.93
123.93
124.93
125.93 Mb
Maternal
exp_1
121.86
122.86
123.86
124.86
125.86 Mb
Paternal
C050−CHA−N10-01
exp_2−3
exp_2−2
exp_2−1
exp_1
121.54
122.54
123.54
124.54 Mb
Maternal
exp_2−3
exp_2−2
exp_2−1
exp_1
121.87
122.87
123.87
124.87
125.87
126.87 Mb
Paternal
C080−CHA−NE20-01
exp_2−3
exp_2−2
exp_2−1
exp_1
121.89
122.89
123.89
124.89
125.89 Mb
Maternal
exp_2−3
exp_2−2
exp_2−1
exp_1
122.09
123.09
124.09
125.09
126.09
127.09
128.09 Mb
Paternal
C102−CZA02-01
exp_2−3
exp_2−2
exp_2−1
exp_1
121.97
122.97
123.97 Mb
Maternal
exp_2−3
exp_2−2
exp_2−1
exp_1
122.46
123.46
124.46
125.46
126.46 Mb
Paternal

**Supplementary Fig. 42 | CENP-A CUT&Tag mapping enrichment and CDR analysis on chromosome 1 across five individuals.** Genomic tracks illustrate the locations of functional kinetochore binding sites characterized by CENP-A peaks in two or one experiment batches (exp_1 and/or exp_2) and centromere dip regions (CDRs) inferred from 5mC methylation frequencies. Black coverages display the primary alignments of CUT&Tag short reads and the red lines denote the locations of parental unique *k*-mers between homologous centromeres, serving as reliable mapping anchors within highly repetitive arrays. 5mC DNA methylation frequencies define localized CDRs, with bottom horizontal tracks representing satellite and higher-order repeat (HOR) annotations for each individual centromere. Profiles on other chromosomes are available at the APG portal (https://genome.zju.edu.cn/APG/download).

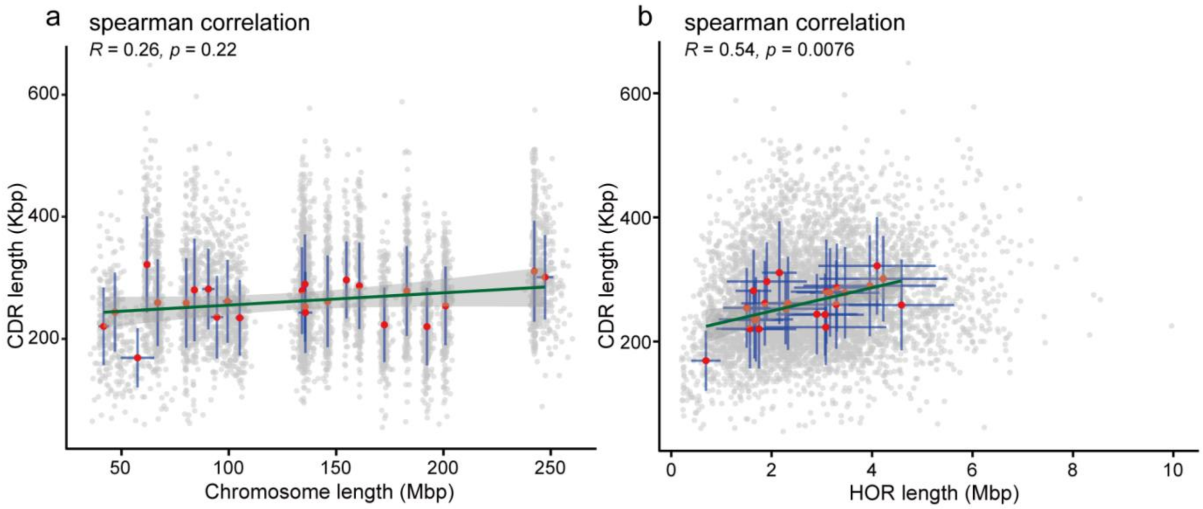


**Supplementary Fig. 43 | Spearman correlation between CDR and chromosome or HOR lengths among APGp1 assemblies. a**, the correlation between the chromosome length and CDR length. **b**, the correlation between the HOR length and CDR length. For both plots, the gray points show the CDR lengths and chromosome or HOR length for each sample; red points indicate the mean length with respect to each chromosome; green lines are the correlation line generated via simple linear regression and blue lines depict the range of Mean ± s.d., standard deviation.

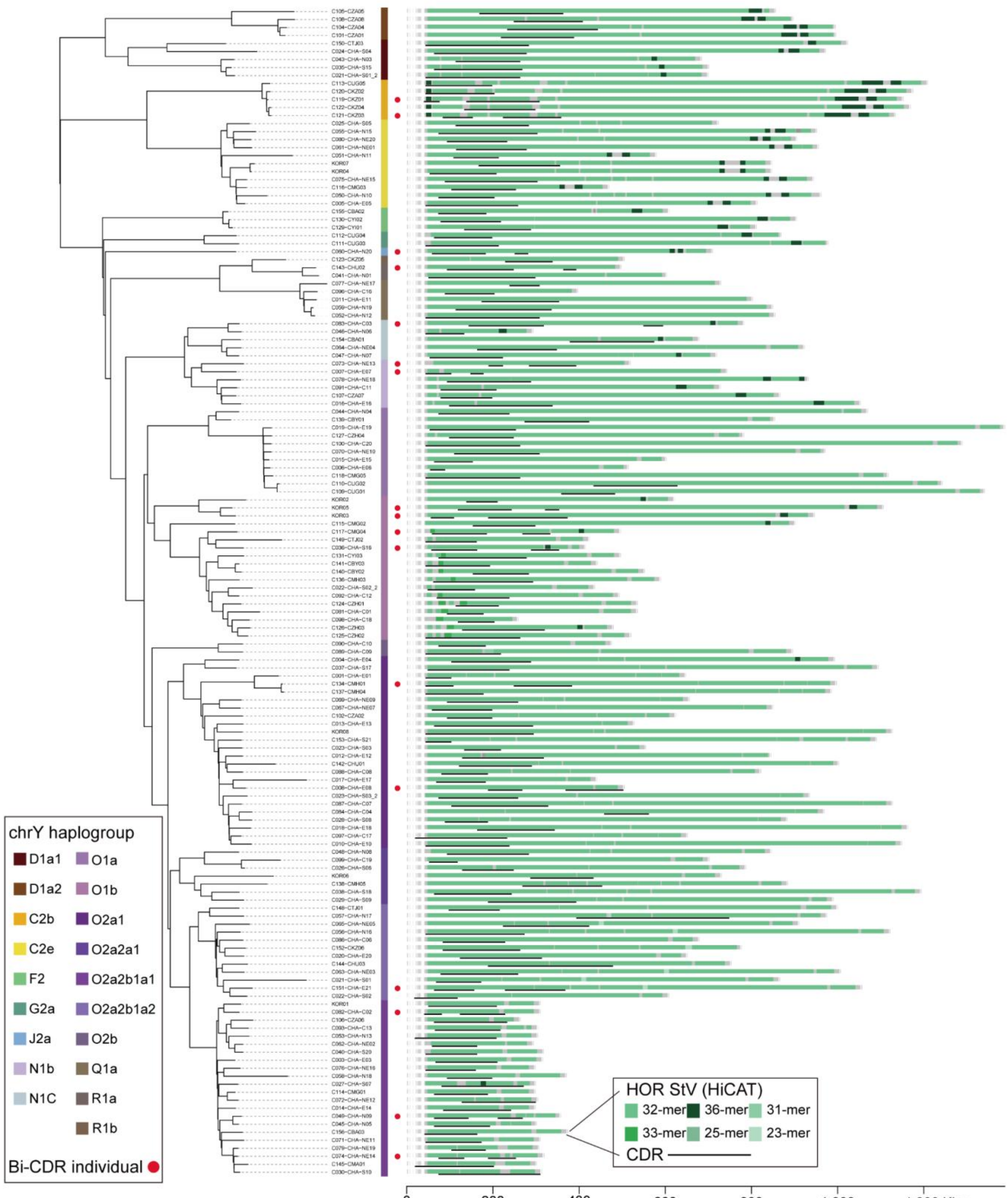


**Supplementary Fig. 44 | Bi-CDRs on chromosome Y.** The bi-CDRs individuals, marked by red points, distribute across 12 haplogroups, suggesting the recurrence of bi-CDRs.

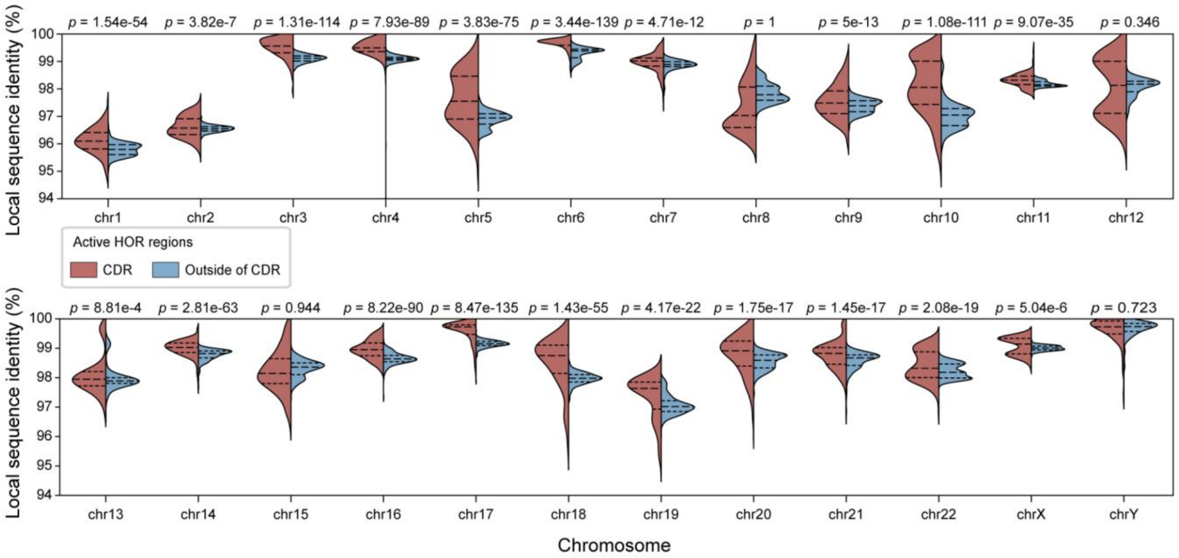


**Supplementary Fig. 45 | Local sequence similarity in CDR and non-CDR active HOR regions for each chromosome.** The dotted lines in each violin plot show the 25%, 50% and 75% percentiles respectively and a one-tail student's *t* test is performed for each chromosome to obtain the *p* value.

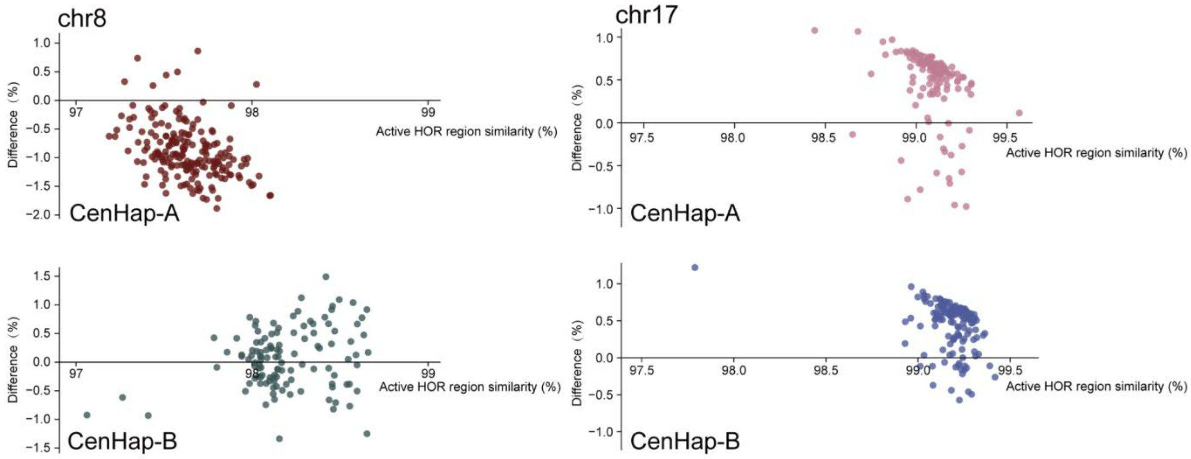


**Supplementary Fig. 46 | Local sequence similarity among CenHaps.** Active HOR regions versus the similarity difference between CDRs and non-CDR active HOR regions for different CenHaps on chromosomes 8 and 17 are shown.

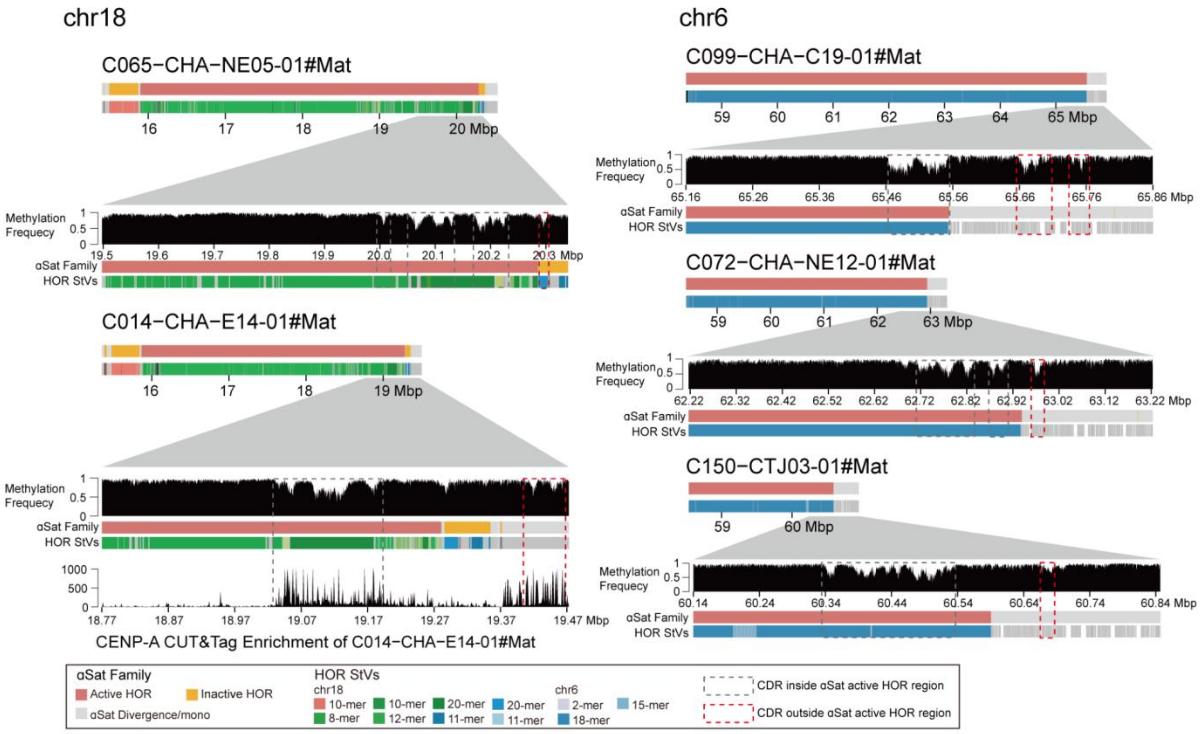


**Supplementary Fig. 47 | The CDRs outside the αSat active HOR regions.** Five centromeres on chromosomes 6 and18 exhibit at least one CDR outside the active HOR regions. The regions containing CDRs outside the active HOR regions are zoomed, with three tracks illustrating methylation, satellite annotation and HOR StVs. Moreover, a track of CENP-A CUT&Tag enrichment is displayed for C014-CHA-E14-01#Mat assembly.

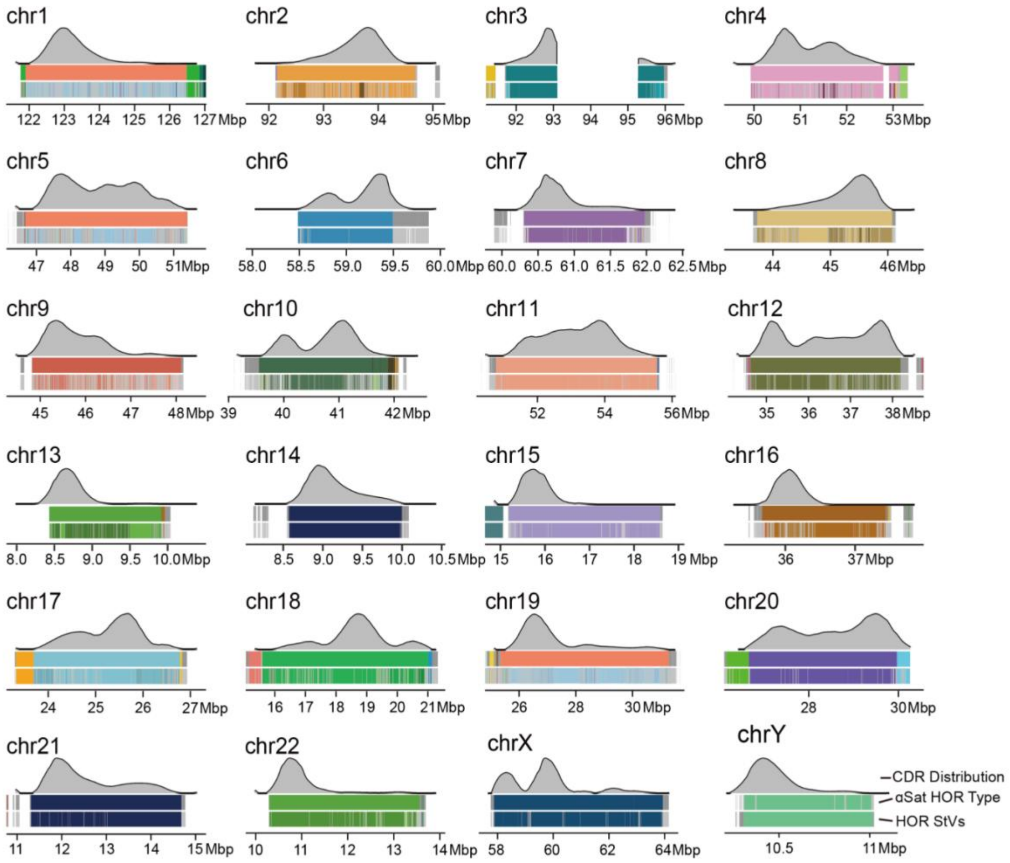


**Supplementary Fig. 48 | CDR distribution across chromosomes.** For each chromosome, the top track shows the CDR distribution when aligned to T2T-CHM13, the middle track displays the αSat HOR types, and the bottom track illustrates the HOR StVs.

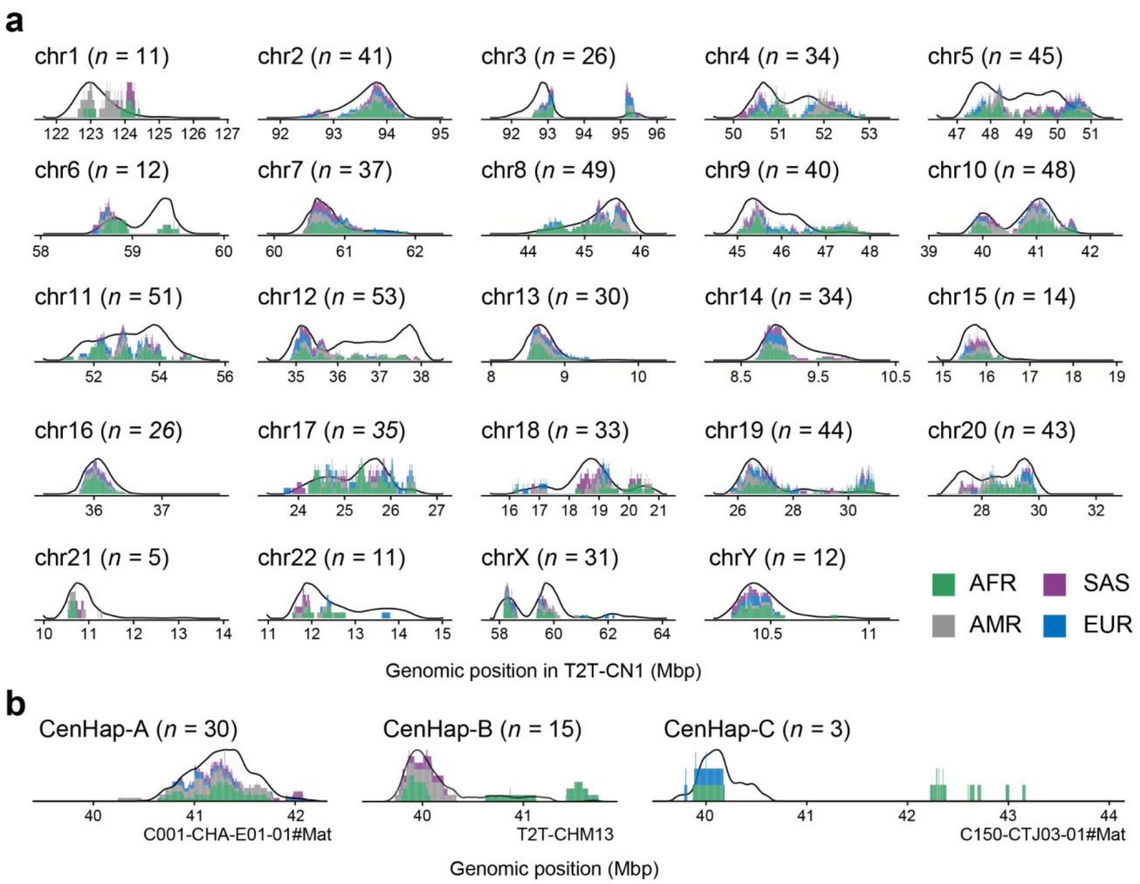


**Supplementary Fig. 49 | CDR positioning across superpopulations from APGp1 and HPRCy1+HGSVC3. a,** Positional distribution of mapped CDRs across chromosomes. **b**, Positional distribution of CDRs stratified across different centromere haplotypes (CenHaps) on chromosome 10. For each panel, the density curves represent the CDR distribution of APGp1 samples (EAS) following coordinate liftover to the reference (T2T-CN1 for **a** or the CenHap-corresponding reference genome for **b**). Stacked bars indicate the CDR positions of samples from the other superpopulations (AFR, AMR, SAS, and EUR) in HPRCy1 and HGSVC3, projected onto the identical coordinate frameworks using the same liftover procedure. Only gapless centromeres are investigated.

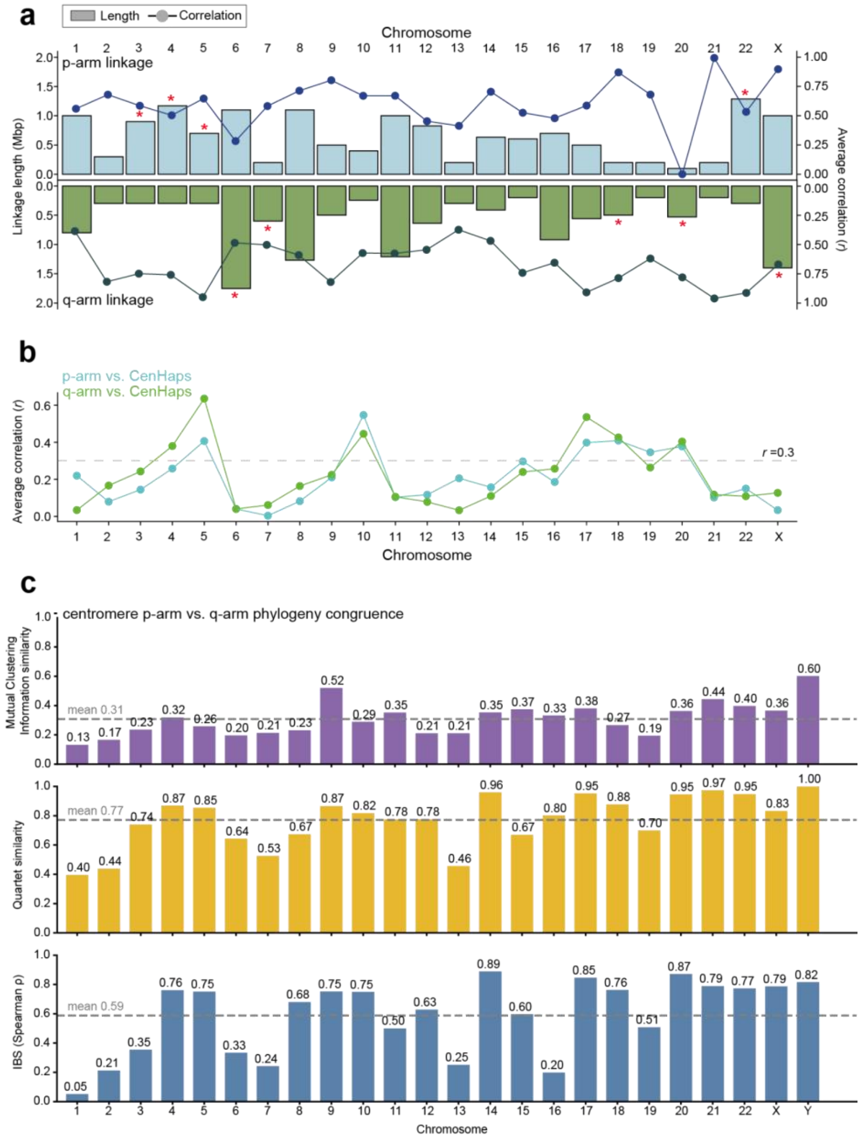


**Supplementary Fig. 50 | Pericentromeric linkage, syntenic distance and phylogenetic concordance across chromosomes. a**, Linkage length and average correlation (Pearson's correlation coefficient *r*) of p-arm and q-arm pericentric regions across chromosomes. Marked asymmetry (>300 Kbp) in length between p- and q-arms is indicated by asterisks (*). **b**, Correlation between linkage and pairwise synteny distance, shown separately for p-arm and q-arm. Interpretation of chromosomes with low correlation ($r < 0.3$) requires caution, which may reflect local expansion or contraction of highly homogenized HOR StV arrays in CenHaps, weakening the synteny–linkage relationship, or limitations in current synteny alignment resolution. **c**, Quantitative phylogenetic concordance between p-arm and q-arm flanking sequence topologies across all chromosomes. Three complementary metrics capture distinct levels of tree similarity. Mutual Clustering Information similarity (top) quantifies shared bipartition information between the two trees via maximum-weight matching of internal splits. Quartet similarity (middle) at bootstrap threshold ≥70, defined as the fraction of randomly sampled four-taxon subsets that induce identical topologies. IBS Spearman rank correlation (ρ; bottom), measuring the concordance of pairwise identity-by-state distance matrices derived directly from the SNP genotype matrix. Dashed grey lines indicate the genome-wide mean values across all chromosomes for each metric. Exact per-chromosome concordance values are annotated above each bar.

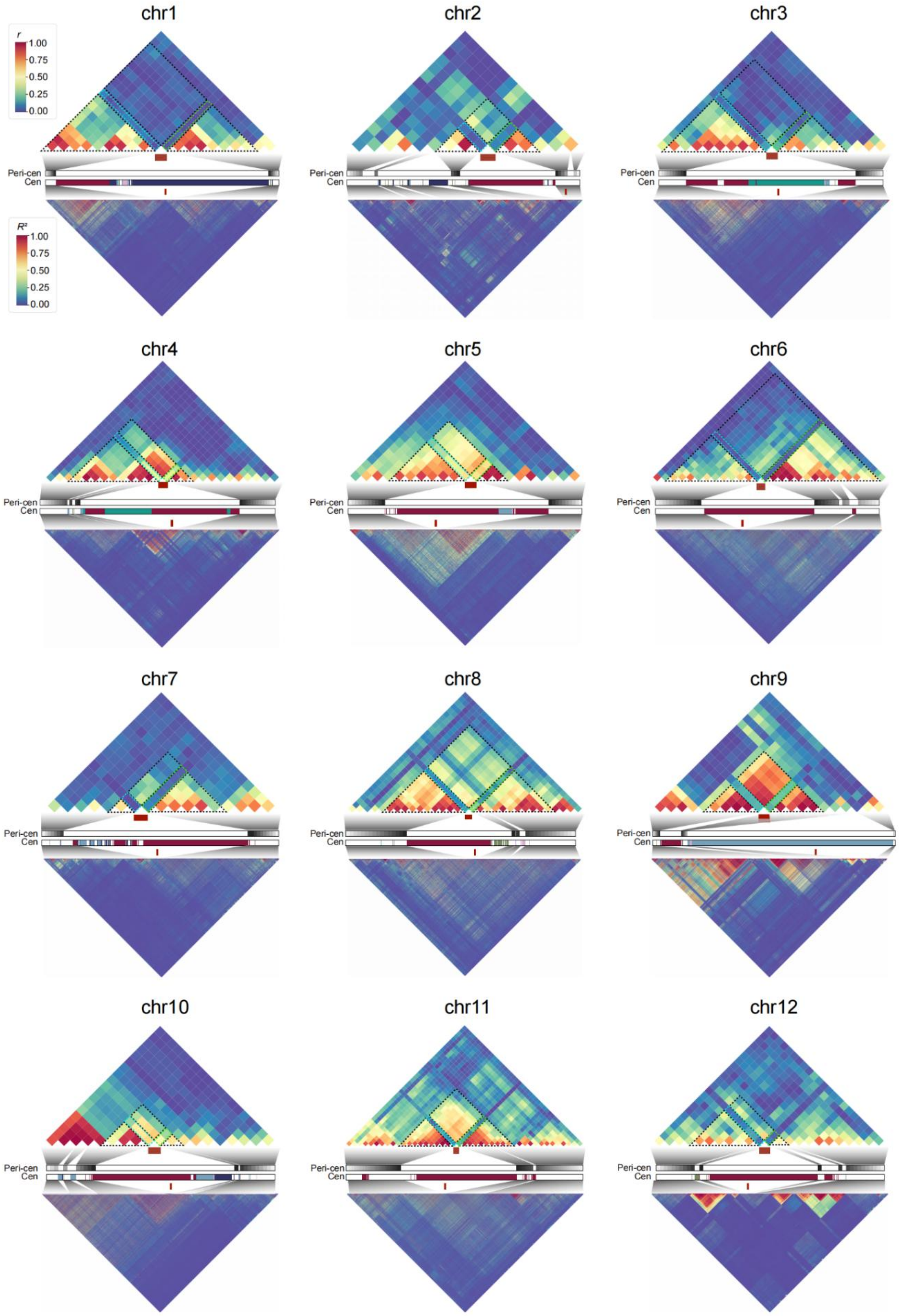

chr1
chr2
chr3
r
1.00
0.75
0.50
0.25
0.00
R²
1.00
0.75
0.50
0.25
0.00
Peri-cen
Cen
chr4
chr5
chr6
chr7
chr8
chr9
chr10
chr11
chr12

**Supplementary Fig. 51 | Linkage landscapes in centromeric regions on chromosomes 1 to 12.** For each chromosome, linkage is depicted from top to bottom as follows: the upper diagonal displays a correlation heatmap (pairwise IBS correlation across populations in 100-Kbp sliding windows), highlighting pericentric linkage. Below, schematic centromere structures and the positions of sliding windows on both flanking regions are shown. The lower diagonal presents a linkage disequilibrium (LD) heatmap based on SNPs from the same sliding window regions. Red rectangles in each triangle denote centromeric positions in the corresponding heatmaps. Blue and green dashed boxes emphasize correlation and linkage length between the p-arm and CenHaps, and the q-arm and CenHaps, respectively.

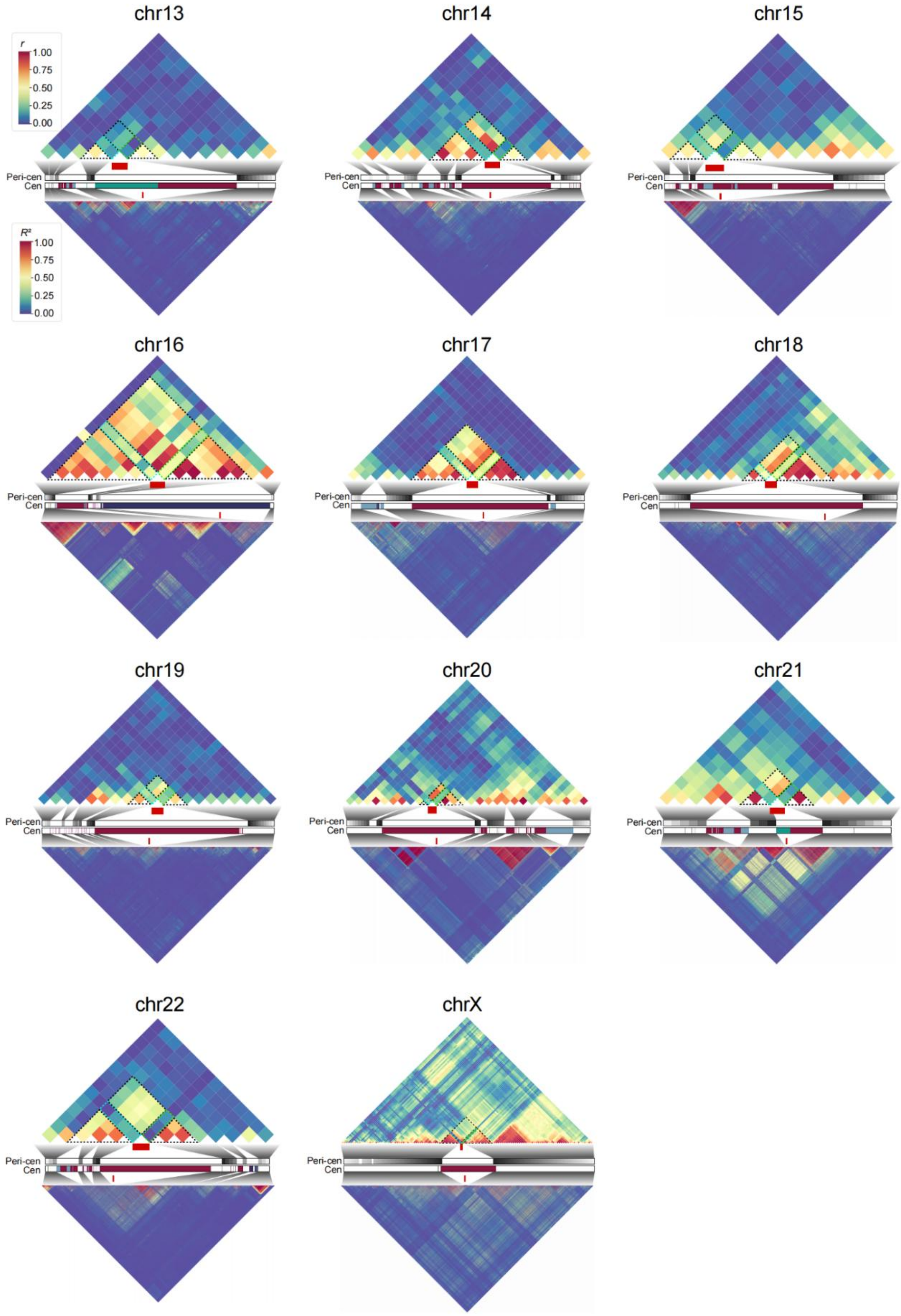
chr13
chr14
chr15
r
1.00
0.75
0.50
0.25
0.00
$R^2$
1.00
0.75
0.50
0.25
0.00
Peri-cen
Cen
chr16
chr17
chr18
chr19
chr20
chr21
chr22
chrX

**Supplementary Fig. 52 | Linkage landscapes in centromeric regions on chromosomes 13 to 22 and X.** For each chromosome, linkage is depicted from top to bottom as follows: the upper diagonal displays a correlation heatmap (pairwise IBS correlation across populations in 100-Kbp sliding windows), highlighting pericentric linkage. Below, schematic centromere structures and the positions of sliding windows on both flanking regions are shown. The lower diagonal presents a linkage disequilibrium (LD) heatmap based on SNPs from the same sliding window regions. Red rectangles in each triangle denote centromeric positions in the corresponding heatmaps. Blue and green dashed boxes emphasize correlation and linkage length between the p-arm and CenHaps, and the q-arm and CenHaps, respectively.

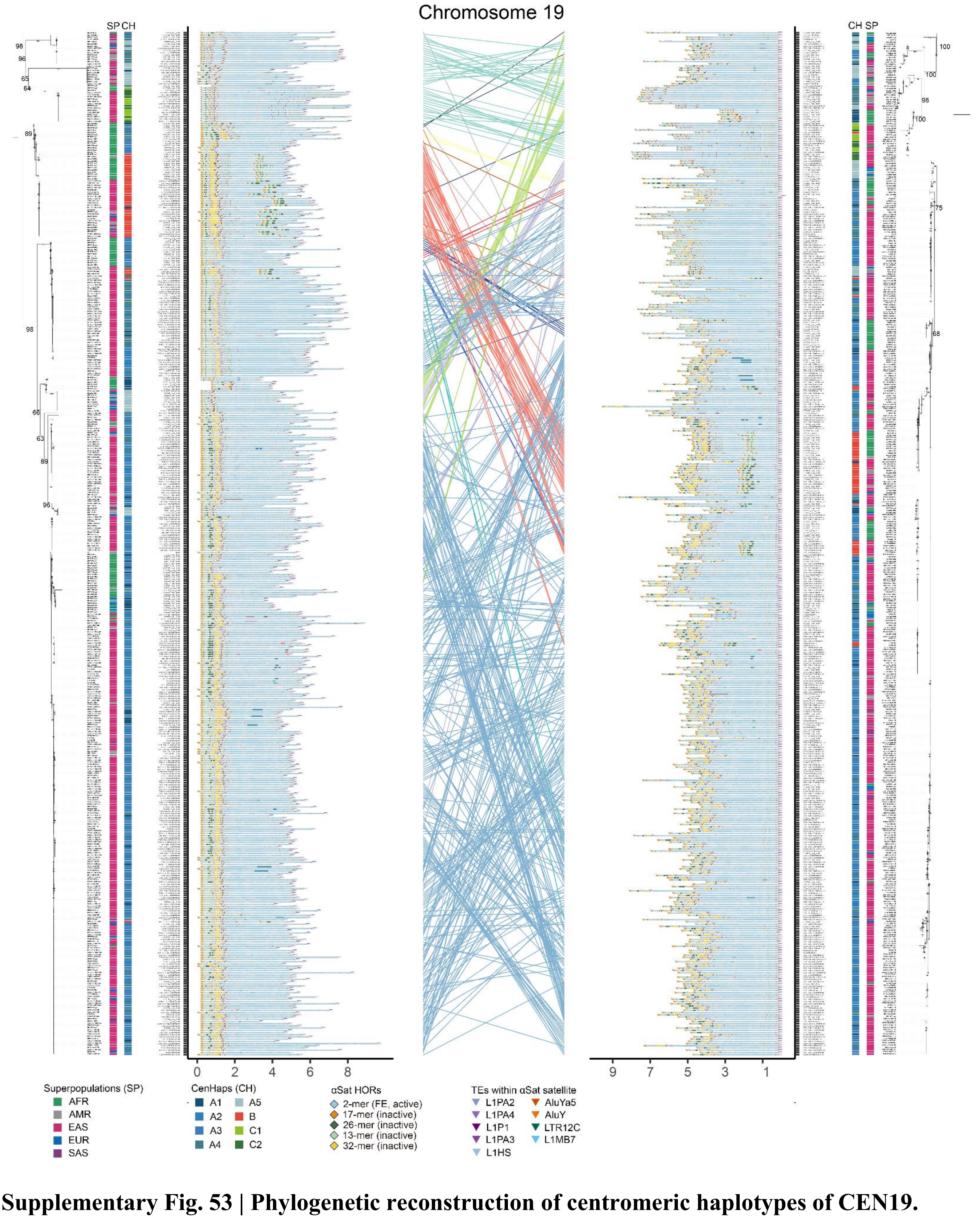


**Supplementary Fig. 53 | Phylogenetic reconstruction of centromeric haplotypes of CEN19.** Phylogenetic analysis based on maximum likelihood for the p- and q-arms of CEN19 reveals discordant topologies highlighted in red line. Bootstrap support values (60%-100%) for key nodes are indicated. Annotation tracks record the superpopulation and CenHap assignments. Corresponding branches between the p- and q-arm trees are connected by colored lines.

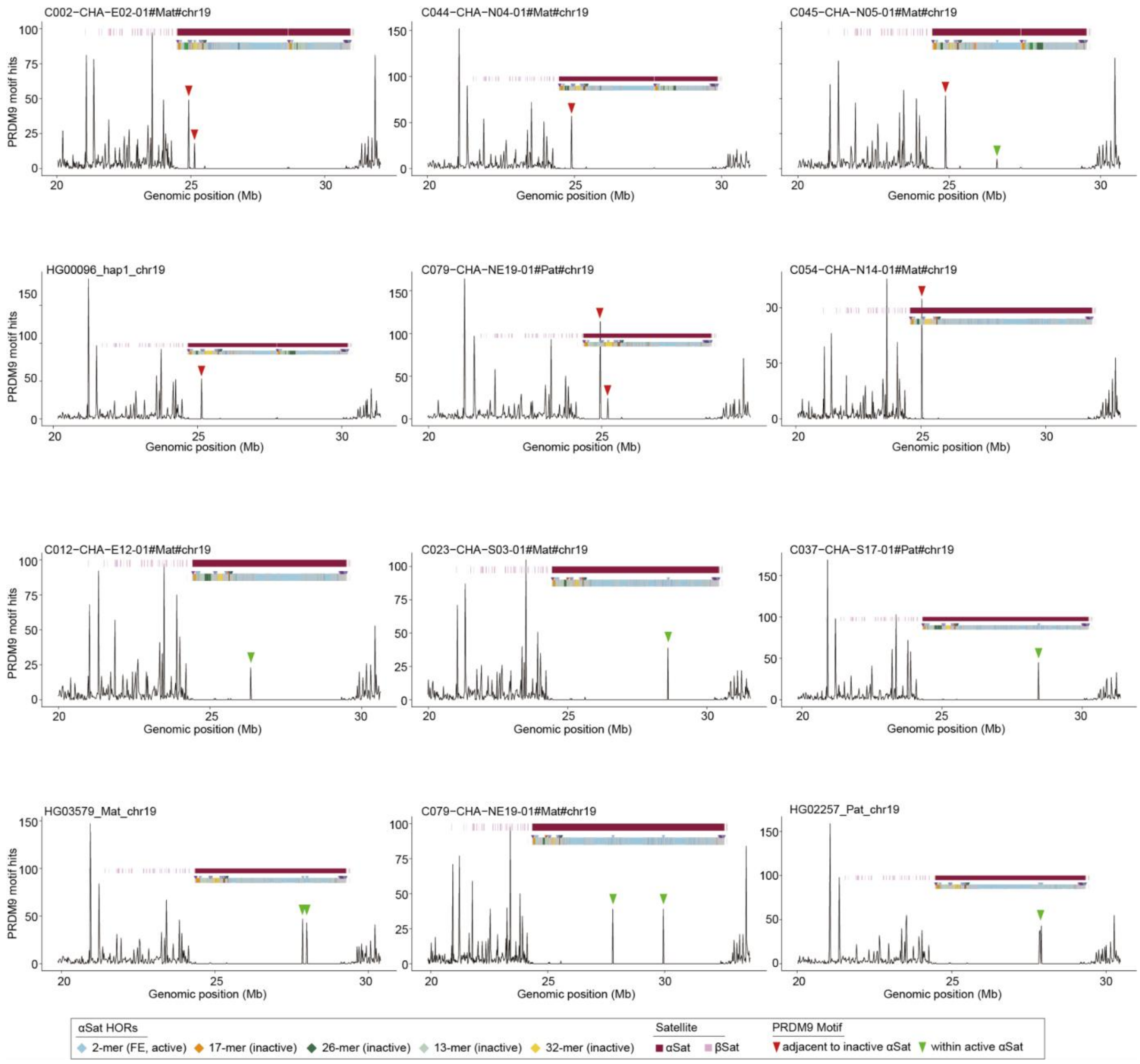


**Supplementary Fig. 54 | Localization of PRDM9 binding sites in human CEN19 αSat arrays.** Identification of two PRDM9 binding sites in CEN19 that precisely colocalize with truncated L1 elements (represented in 12 haplotypes). The first site (red), adjacent to inactive αSat, is enriched in CenHap-B (32/49). The second (green), within the active array, occurs in 17 centromeres (9 with discordant phylogeny).
