## Supplementary Notes for "Multidimensional variation and population stratification across 8000 complete human centromeres"

#### Supplementary Note 1 | Quality evaluation of centromere assemblies

Given the highly repetitive nature of human centromeres, we performed a comprehensive assessment on human centromere assemblies generated in APGp1 in assembly continuity and completeness, using multiple metrics (**Supplementary Table 1**).

##### 1.1 GCI

First, we applied the Genome Continuity Inspector[1] (GCI), a tool specifically developed to evaluate the quality of complete genome assemblies. GCI utilizes alignments of PacBio HiFi and Oxford Nanopore Technologies (ONT) reads that undergo stringent filtering, generating overall continuity scores and flagging potentially problematic regions. Gapless centromeres achieved an average GCI score of 96.7, indicative of high assembly continuity (**Supplementary Fig. 1c**; **Supplementary Table 1**). Analysis of PacBio HiFi and ONT read coverage revealed uniform distributions across majority of centromeres, with ~1.63% (3.58 Mbp) and ~0.71% (1.55 Mbp) of centromere regions per haploid exhibiting limited coverage as identified by GCI for PacBio HiFi and ONT reads, respectively (**Supplementary Fig. 1d**).

##### 1.2 NucFreq

Second, we employed NucFreq[2] to detect heterozygous sites and assembly collapses by analyzing the frequency distributions of the first and second most common nucleotides in aligned PacBio HiFi reads. This approach identified a limited number of high-heterozygosity loci, totaling 2.67 Mbp (2.38% per haploid), where the second most frequent alleles were supported by at least 10% of reads (**Supplementary Fig. 1d**).

##### 1.3 Flagger

Third, Flagger[3] was applied to evaluate assembly quality and identify problematic regions, including those potentially erroneous, duplicated, and collapsed. Considering the sequencing bias of PacBio HiFi sequencing in specific regions[4–6], we adjusted Hsat2 and Hsat3 region assessments based on platform-specific coverage thresholds (PacBio Revio: 0.55; PacBio Sequel II: 1.2), and incorporated Flagger-supported high-confidence mappings to minimize false positives. Following refinement, Flagger analysis indicated minor proportions of such regions per haploid: approximately 1.75 Mbp (0.82%) of potential errors, 3.78 Mbp (1.75%) of duplications, and 3.78 Mbp (1.75%) of collapses (**Supplementary Fig. 1e**).

##### 1.4 GAVISUNK and VerityMap

Fourth, we assessed centromere assemblies using GAVISUNK[7] and VerityMap[8]. These tools assess concordance between the assembly and unique *k*-mers derived from ONT and PacBio HiFi reads, respectively (**Supplementary Figs. 20-21** and **26**).

### 1.5 QV

Additionally, we used Merqury[9] (v1.4.1) to calculate assembly quality value (QV) in centromeric regions under default settings, based on a hybrid *K*-mer database ($k = 21$) constructed from both PCR-free NGS and PacBio HiFi reads. Gapless centromeres in APGp1 achieved an average QV of 87.3, reflecting a superior standard of base-level accuracy.

In summary, regions flagged as potential assembly issues by at least two methods averaged 1.03 Mbp, representing 0.47% of the centromeric sequence per haploid (**Supplementary Table 1**). On average, 19.7 centromeres were validated as completely assembled per haploid with comprehensive chromosomal representation averaging 273 centromeres per euchromosome and 153 per sex chromosome.

### Supplementary Note 2 | Global αSat clustering and HOR *de novo* inference

#### 2.1 Global αSat monomer clustering

To mitigate potential bias using the existing database capturing limited sequence diversity in High-Order Repeat (HOR) inference across multi-ancestry genomes, we developed a novel workflow for global monomer clustering. αSat monomer positions were identified through alignment to a consensus database generated for each monomeric class from the T2T-CHM13 assembly. To minimize potential bias introduced by fragment length variation, monomers shorter than 120 bp or longer than 200 bp were excluded with the command "*seqkit seq -m 120 -M 200 -g all.mn.fasta -o all.mn.filtered.fasta*". Finally, approximately 244 million αSat monomer sequences were identified and extracted from pangenome assemblies of APGp1, HPRCy1, HGSVC3, and reference-level complete assemblies (T2T-CHM13, T2T-CN1, Q100-HG002, and YAO). We then employed VSEARCH[10] (v2.30.0), a fast heuristic-based tool for sequence search and clustering, to remove duplicated monomer sequences with following command: "*vsearch --derep_fulllength all.mn.filtered.fasta --output all.mn.filtered.dedup.fasta --sizeout --minuniquesize 1*". Subsequently, deduplicated sequences were clustered by abundance at identity thresholds ranging from 87% to 99%, using the command: "*vsearch --cluster_sizeall.mn.filtered.dedup.fasta --id ${threshold} --iddef 0 --centroids all.centroid.fasta --uc all.uc --clusters original_seq/*".

Given the greedy nature of this clustering algorithm, we implemented a graph-based refinement procedure (**Fig. 3a**). Centroid sequences from the initial VSEARCH clusters were used to construct a similarity graph, where nodes represent centroids and edges connect node pairs with a sequence identity above the initial clustering threshold with edge weights scaled as similarity ×100. We then applied the Louvain algorithm to this graph to detect community structure, thereby regrouping the initial centroids into more robust clusters (**Fig. 3a**). To further ensure cluster homogeneity and prevent the over-inclusion of weakly linked sequences, we introduced a hierarchical refinement step, which evaluates the cluster integrity by calculating all pairwise identities between centroids within a community. Guided by the triangle inequality principle, we defined a minimum pairwise identity threshold of $2\times\$\{\text{threshold}\}-1$ for cluster coherence. Any cluster failing to meet this criterion was recursively split (**Fig. 3a**). For splitting, centroid similarity graphs were reconstructed within each heterogeneous community using a predefined identity cutoff, followed by Louvain community detection to identify independent subclusters. To evaluate the effectiveness of the splitting procedure, we randomly sampled 1,000 monomers from representative examples before and after splitting and compared intra- and inter-cluster sequence identity distributions, confirming improved cluster separation and internal homogeneity (**Supplementary Note Fig. S1**). Finally, we applied a core-periphery strategy: clusters were ranked by size in descending order, and those accounting for the top 90% of all monomer sequences were designated as core communities. The remaining smaller clusters were assigned to a core cluster if their minimum pairwise identity with it exceeded a defined threshold; otherwise, they were classified as new clusters. The final output of this pipeline is a set of non-redundant, high-integrity monomer clusters, each assigned a unique numeric identifier. These cluster assignments were then encoded as ID strings to serve as the basis for all subsequent HOR inference analyses.

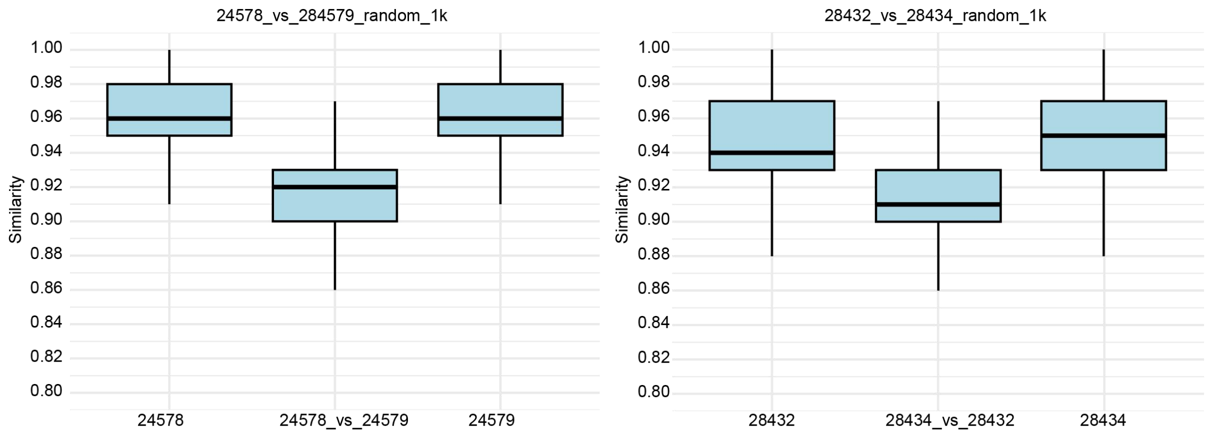


**Supplementary Note Fig. S1 | Evaluation of cluster splitting by sequence similarity analysis.** To evaluate the effectiveness of the recursive cluster splitting procedure, two representative examples are shown for monomer clusters that were separated into distinct subclusters: cluster 24578 was split into 24578 and 24579, and cluster 24832 was split into 24832 and 24834. For each example, boxplots show the distribution of pairwise sequence identities among monomers within each resulting subcluster (intra-cluster similarity) and between the two separated subclusters (inter-cluster similarity), estimated from randomly sampled 1,000 monomers per cluster. These refined clusters were subsequently used to resolve distinct HOR variants, including HORs associated with CEN13 and CEN10 centromere haplotypes respectively, thereby improving discrimination of centromeric sequence diversity.

#### 2.2 Evaluation of global αSat monomer clustering

To assess the accuracy of our monomer clustering, we compared the resulting clusters against the previously established HOR annotations in T2T-CHM13[11]. We quantified the similarity between the two sets of clusters using a weighted Jaccard score. This metric is designed to emphasize the agreement between larger, more significant clusters, rather than treating all clusters equally. The weighted Jaccard score is calculated as follows:

$$Weighted\ Jaccard\ Score = \frac{\sum_{i=1}^{n}\sum_{j=1}^{m} J(A_i,B_j) \times W(A_i,B_j)}{\sum_{i=1}^{n}\sum_{j=1}^{m} W(A_i,B_j)}$$

where $A = \{A_1, A_2, \ldots A_n\}$ represents the set of clusters from the new method. $B = \{B_1, B_2, \ldots B_m\}$ represents the set of reference HOR clusters from T2T-CHM13[11].

$J(A_i,B_j) = \frac{|A_i \cap B_j|}{|A_i \cup B_j|}$ is the classical Jaccard similarity coefficient between cluster $A_i$ and $B_j$, measuring their overlap. $W(A_i,B_j) = \frac{|A_i \cap B_j|}{|A_i|} \times \frac{|A_i \cap B_j|}{|B_j|}$ is the weight assigned to the pair. This strategy ensures that a high weight is only assigned when the intersection constitutes a large proportion of both clusters, thereby prioritizing one-to-one correspondences over fragmented or partial matches.

To avoid excessive fragmentation of αSat arrays at overly stringent thresholds, we selected an identity cutoff of 0.96, which also provided the closest concordance with the T2T-CHM13 annotation, (**Supplementary Note Fig. S2**), with a weighted Jaccard Score of 0.74 for whole centromeres and 0.95 for active arrays.

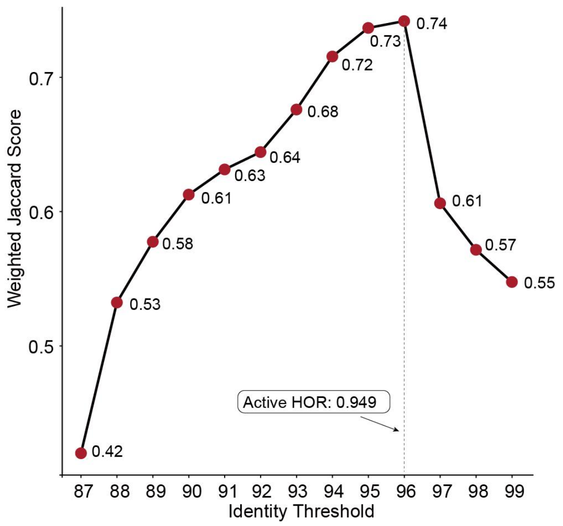


**Supplementary Note Fig. S2 | Evaluating the performance of monomer clustering with different similarity thresholds.** The similarity threshold of 0.96 optimally recapitulates previous monomer classification in T2T-CHM13, yielding a weighted Jaccard Score of 0.74 for whole centromeres and 0.95 for active arrays.

##### 2.3 Suprachromosomal families of αSat monomer clusters

As previously described[11], human αSat monomers were classified into 20 distinct suprachromosomal families (SFs) based on sequence identity and structural features. A systematic nomenclature was proposed for αSat higher-order repeats (HORs), in which each HOR was assigned a name indicating its suprachromosomal family, chromosomal location, and an index number—for example, S1C13/21H1 denotes a HOR composed of αSat monomers from SF1, on chromosomes 13 and 21, numbered as HOR #1[11]. Accordingly, by aligning each monomer class to a consensus database and retrieving the best hit (**Methods**), we obtained the corresponding nomenclature. From this, we extracted the segment following the prefix "S" to assign each monomer sequence to its respective SF category. For each monomer cluster, we quantified the distribution of SF assignments among its constituent sequences. The vast majority of clusters (92.1%, 27,414 of 29,773) were uniformly assigned to single SFs. In cases where a cluster contained sequences assigned to two or more SFs, the cluster was assigned to corresponding dominant SF type if one SF accounted for more than 80% of the sequences within this cluster. Otherwise, the remaining multi-SF clusters were classified as "Mixed". All SF assignment results are provided in **Supplementary Table 12**.

Of all the defined αSat monomer clusters, 3,722 were organized into HOR arrays. Representative monomer sequences were selected using a size-aware sampling approach: from clusters initially containing over 40,000 sequences, three monomer sequences were randomly sampled to capture potential diversity, whereas a single representative was chosen from the remaining clusters. Subsequently, the selected αSat sequences (8,713 in total) were aligned using MAFFT[12] (v7.525), and a maximum-likelihood phylogenetic tree was constructed using IQ-TREE[13] (v2.3.6) with the parameter of "*-m MFP -B 1000 --bnni -T AUT*O". Although the phylogeny recovered eight well-supported clades, it showed a 3.1% discordance with established suprachromosomal family (SF) classifications. (**Supplementary Fig. 22**).

##### 2.4 HOR inference

###### 2.4.1 HORmon (graph-based strategy)

The HORmon algorithm was developed based on the Centromere Evolution (CE) Postulate, which states that each extant centromere has evolved from a single ancestral HOR unit comprising *k* distinct monomers, implying that each monomer occurs only once within that ancestral HOR structure[12]. The algorithm integrates monomer identification and HOR inference by first constructing a *de Bruijn* graph of monomer clusters connected by edges, whose multiplicity reflects the observed succession frequency between adjacent monomers, then refining monomer assignments via positional similarity to resolve the graph into a single cycle representing the inferred HOR, and finally, classifying detected HORs into canonical and partial types.

In the present study, we decoupled monomer generation from HOR inference, adopting a strategy similar to the CentromereArchitect[13]. We posited that a global, population-based classification of monomers could delineate biologically distinct monomer blocks. Using BEDTools[14] (v2.31.1), we firstly merged αSat monomers with intervals less than 10 Kbp into continuous genomic blocks, with each monomer assigned a global cluster ID as input. A *de Bruijn* monomer graph was then constructed to identify cycles representing putative canonical HORs.

Monomer graphs built from these continuous αSat blocks can exhibit substantial complexity in specific genomic contexts, particularly when spanning megabase-scale arrays that encompass both active and inactive centromeric domains, as observed on chromosome 1. This structural complexity is manifested by a high number of nodes interconnected through intricate edge patterns. To resolve such graphs, we implemented a tiered simplification strategy based on edge multiplicity. Beginning from the highest multiplicity level, which is primarily associated with active arrays, we systematically identified cycles, enabling robust HOR inference within these architecturally complex regions (**Supplementary Note Fig. S3**).

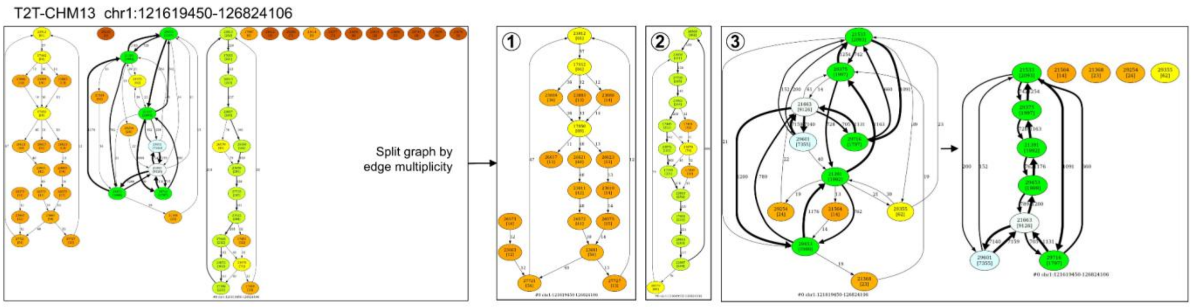


**Supplementary Note Fig. S3 | Monomer graph of CEN01 in the T2T-CHM13 assembly.** The initial monomer graph (left) shows a dense network of nodes connected by numerous high- and low-multiplicity edges. To resolve this architecture, edges were progressively filtered by edge multiplicity. Iterative pruning and cycle detection (middle to right) reduce graph complexity and reveal the underlying HOR structure.

The Pan-HOR structural variants (StVs) dataset identified by HORmon includes both canonical and partial HORs, collectively designated as StVs. All StVs with cyclic permutations—such as 1-2-3-4, 4-1-2-3, 3-4-1-2, and 2-3-4-1—were consolidated into a single canonical form (e. g. 1-2-3-4). During cycle detection, only canonical HORs with a multiplicity greater than 2 (i. e. appearing at least three times within a monomer block) were retained. These pan-HOR StVs were subsequently categorized into five classes based on population frequency: core (>90% in frequency), dispensable (5%-90%), low-frequency (1%-5%), rare (<1% and $n > 1$), and singleton ($n = 1$).

##### 2.4.2 HiCAT (HTRM-based strategy)

HiCAT (Hierarchical Centromere Structure Annotation) enables automated detection of locally nested HORs through hierarchical tandem repeat mining (HTRM), a bottom-up iterative compression approach[15]. We began by merging αSat monomers into contiguous blocks (<10 Kbp) using BEDTools, then converted each monomer into its corresponding global clustering ID as input. The HTRM module was applied to iteratively identify and compress local tandem repeats (TRs), with the maximum unit length was set to 40 monomers for autosomes and chromosome X, and 50 for chromosome Y.

TRs with cyclic permutations—such as 1-2-3-4, 4-1-2-3, 3-4-1-2, and 2-3-4-1—were unified into a single canonical form (e.g., 1-2-3-4). A TR was classified as an HOR if it met two criteria: (1) contained at least two distinct monomers; (2) displayed at least three contiguous repeats (i.e. "nrepeat" ≥ 3 in at least one haploid assembly). TRs with a maximum of two contiguous repeats across all haploids were categorized as dimers, regardless of monomer composition. The resulting pan-HOR StVs encompassed both top-layer and cover-layer HORs, reflecting a hierarchical organization that may correspond to distinct evolutionary stages of repeat compaction. These StVs were stratified into five frequency-based classes: core (>90%), dispensable (5-90%), low-frequency (1-5%), rare (<1% and $n$ >1), and singleton ($n$ = 1).

Given that HiCAT's iterative compression can generate structurally divergent top-layer outputs across individuals, we implemented a decompression and reannotation step to facilitate inter-individual comparison. Each StV was annotated within a parent–child hierarchy (root.hierarchy → child.hierarchy), and coverage relative to the top layer was computed. Cover-layer HORs with >60% coverage replaced their top-layer counterparts for visualization purposes exclusively; all statistical analyses retained the complete set of top-layer and cover-layer HORs.

#### 2.5 HOR array annotation and evaluation

HOR arrays are highly periodic genomic structures in which distinct αSat monomers are organized in a head-to-tail arrangement to form canonical HOR units. In the T2T-CHM13 annotation, αSat arrays are classified as active, inactive, or divergent based on unit composition, size, kinetochore binding potential, and intra-array divergence[11]. Within a given array, monomer-scale deletions generate StVs representing alternative HOR types. We integrated the Pan-HOR StVs inferred from HORmon and HiCAT to identify HOR arrays. Pairwise string alignment (https://github.com/alevchuk/pairwise-alignment-in-python/alignment.py) was applied to quantify monomer composition and calculate the alignment match ratio (max_match_string / length (StV1, StV2)). StVs with alignment match ratio ≥ 0.5 were clustered and considered part of the same array.

Compared with previously reported arrays, we recalled 57 of 60 known HOR arrays in T2T-CHM13, with six merged due to methodological differences. In detail, sister HOR arrays are defined as structurally similar but sequence-divergent (2-5%) in T2T-CHM13, with some monomers different and some conserved, whereas our clustering merges any StVs with match ratio ≥ 0.5 based on string alignment, thus merging sister HORs (annotated as "2_vs_1" in **Supplementary Table 13**). Regarding the three undetected arrays, all were annotated as inactive in T2T-CHM13.The monomeric sequences composing these arrays exhibited divergence exceeding our identity threshold, leading to their assignment to different monomer cluster IDs and preventing the detection of a periodic, recognizable HOR pattern in our analysis.

### Supplementary Note 3 | HOR sharing

##### 3.1 Chromosome-specific HOR StVs in active arrays of CEN01/05/19

Active HOR arrays on chromosomes 1, 5 and 19 are known to be highly similar[11]. To characterize the shared patterns of HOR StVs among them at the population level, we analyzed StVs identified by HiCAT due to the abundant local nested HORs within the canonical HOR units existing in these centromeres. Our analysis revealed both chromosome-specific StVs and StVs shared across all three chromosomes (**Fig. 4d**; **Supplementary Figs. 27** and **28**).

In CEN01, the dominant HOR StV is an 8-mer denoted as "a2-B-C-D-E-F-E-G" (monomer cluster: 21391_29375_21533_29716_21663_29601_21663_29453, corresponding to the T2T-CHM13 annotation "S1C1/5/19H1L.1-5_6/4_5_6"), which contains a local-nested (FE)*n* dimer. Among the 55 haploid genomes in APGp1, this 8-mer on CEN01 was observed in the inverted order "G-E-F-E-D-C-B-a2" that is considered as another StV, resulting from large megabase-scale inversions.

In CEN05, two dominant type of HOR StVs were identified: The first is an 8-mer defined as "a1-B-C-D-E-F-E-H" (monomer cluster: 21377_29375_21533_29716_21663_29601_21663_29179, equivalent to “S1C1/5/19H1L.1-5_6/4_5_2/6”). The other type comprised a set of 12-mer subsets (e.g., "a1-B-C-D-E-F-E-H-a1-I-E-H", "A-B-C-D-E-F-E-H-a1-I-E-H" with monomer clusters “21377_29375_21533_29716_21663_29601_21663_29179_21377_29254_21663_29179”, “21368_29375_21533_29716_21663_29601_21663_29179_21377_29254_21663_29179”), whose key variation is the replacement of the A monomer with a derived allele (S1C1/5/19H1L.1), collectively annotated in T2T-CHM13 as "S1C1/5/19H1L.1-5_6/4_5_2/6_1_2/4_5_2/6".

In CEN19, three distinct structural layers were observed (**Supplementary Fig. 27**). Region 1 (spanning the beginning ~100 Kbp of the active array) and Region 3 (spanning the ending ~100 Kbp) are primarily composed of the most polymorphic and shared StVs. These include diverse HOR StVs such as a 6-mer (*e. g.* "a1-B-C-F-E-R", "A-B-C-F-E-R"; monomer clusters: "21377_29375_21533_29601_21663_29205", "21368_29375_21533_29601_21663_29205"; corresponding to T2T-CHM13 "S1C1/5/19H1L.1-3_6/4_5_2/6"), an 8-mer ("A-B-C-D-E-F-E-R", "a2-B-C-D-E-F-E-G" that shared with CEN01), and 12-mers (shared with CEN05). In contrast, the central Region 2 spans megabases and is predominantly composed of a simple 2-mer "FE" repeat (monomer clusters: "29601_21663", “29759_21663”; T2T-CHM13: S1C1/5/19H1L.6/4_5). A similar "FE"-dominant structure was reported in CEN19 of T2T-CHM13 and was considered as a secondary expansion. This tri-layered architecture was observed in all complete CEN19 in our datasets, suggesting it is a conserved and inherited feature in human evolution.

##### 3.2 Phylogeny of shared HOR segments

Based on chromosomal-specific StVs we described above, we constructed the phylogeny using shared segments of HOR StVs using “A-B-C” and FE dimers, respectively (**Fig. 4e**; **Supplementary Fig. 28**). To perform a spatially unbiased sampling, we extracted shared segment sequences from three regions of the active array: Region 1 (p-arm flanking region), covering 0-100 Kbp from the active array start; Region 2 (central region), spanning the midpoint ± 50 Kbp; and Region 3 (q-arm flanking region), comprising the terminal 100 Kbp. From each region, we randomly sampled 1,000 shared sequences. The pooled sequences were aligned using MAFFT[16] (v7.525), and used to construct maximum-likelihood phylogenies with IQ-TREE[17] (v2.3.6) with the parameter of “*-m MFP -B 1000 --bnni -T AUTO*”. This procedure was repeated ten times to assess the robustness of tree topology. The results showed that CEN19 exhibited the highest sequence variant (SqV) polymorphisms. In summary, our population analysis of the sharing pattern between CEN01/05/19 supports a previous hypothesis that CEN19 represents the most ancestral centromere among these chromosomes.

##### 3.3 A close relationship between CEN19 and CEN16

In CEN16, two dominant HOR StVs were observed (**Fig. 4d**). 362 haploid assemblies displayed massive repeats of the 10-mer canonical HOR unit denoted as "A-K-I-F-E-M-N-O-P-Q" (monomer cluster: 21368_29290_21878_29601_21663_29427_21448_28754_21363_29305, corresponding to T2T-CHM13 "S1C16H1L.1-10"), in which three monomers (A/F/E) are shared with CEN01/05/19. In contrast, two haploid assemblies from AFR exhibited 1172 and 554 copies of 8-mer HOR StVs ("K-L-F-E-M-N-O-P", monomer clusters: 29290_21878_29601_21663_29427_21448_28754_21363, corresponding to T2T-CHM13 "S1C16H1L.2-9") that also comprised of a local-nested (FE)*n* dimer, analogous to CEN01/05/19 (**Supplementary Fig. 27**). The phylogenetic tree revealed that FE sequences from CEN16 were nested within those from CEN19, similar as CEN01 and CEN05, suggesting the origination of CEN16's dimeric HORs from CEN19. Additional evidence comes from their reciprocal monomer incorporation in the flanking regions of their active arrays, a pattern absent between CEN16 and CEN01/05 (**Supplementary Fig. 29**). Taken together, these findings strongly support a uniquely close evolutionary link between these two centromeres of human chromosomes 16 and 19.

### Supplementary Note 4 | Single-base substitution rate analysis in human centromeres

#### 4.1 Overview

Accurately quantifying single-base substitution rates in human centromeres remains a significant challenge, largely due to their extreme repetitiveness and dynamic expansions/contractions that generate extensive large-scale structural variations (SVs) and structural HOR variants (StVs). To address these, we developed a conservative computational pipeline to compare substitution rates between core αSat HOR arrays and their flanking pericentromeric regions, while minimizing confounding effects from structural variations and paralogy. The pipeline's core design principle centers on establishing a structurally stable analytical background by restricting analysis to centromere pairs with extremely high identity, and then defining callable regions with robust orthology support. Through a sequential combination of algorithmic filters (HORSCAN-based alignment score, masking of structurally unstable segments, and pericentromeric collinearity) followed by manual curation, we identified a high-fidelity panel of 93 centromere pairs exhibiting mean structural identity ≥ 0.99 (**Supplementary Table 20**). This curated panel enables unbiased comparison of substitution rates between core centromeres and their pericentromeric genomic context, mitigating the technical artifacts inherent to analyzing repetitive genomic landscapes.

#### 4.2 High-identity centromere pairs

High structural polymorphism and sequence repetitiveness of αSat HOR arrays severely compromises the performance of standard alignment algorithms, such as minimap2[18], Winnowmap2[19] and UniAligner[20]. Therefore, we employed the newly developed tool HORSCAN (v1.0; https://github.com/XDwan/HORSCAN), a monomer-level alignment tool based on multi-level dynamic programming, to perform all-*vs*-all pairwise alignments among the gapless centromeres from APGp1, HPRCy1 and HGSVC3, to identify structurally homologous centromere pairs within each chromosome. This resulted in 1,737,685 pairwise alignments.For each pair, we defined a HORSCAN alignment score to represent structural homology and identity as:

$$Score = \frac{N_{MTH}}{N_{total}}$$

where monomer insertions and deletions are treated as monomer-level structural variants (SVs). Higher scores indicate the greater similarity. To focus on structurally homologous candidate pairs, we filtered the centromere alignments by HORSCAN alignment scores ≥ 0.9 and generated a collection of 94,899 candidate pairs with high identity for subsequent filtration (**Supplementary Note Fig. S4**).

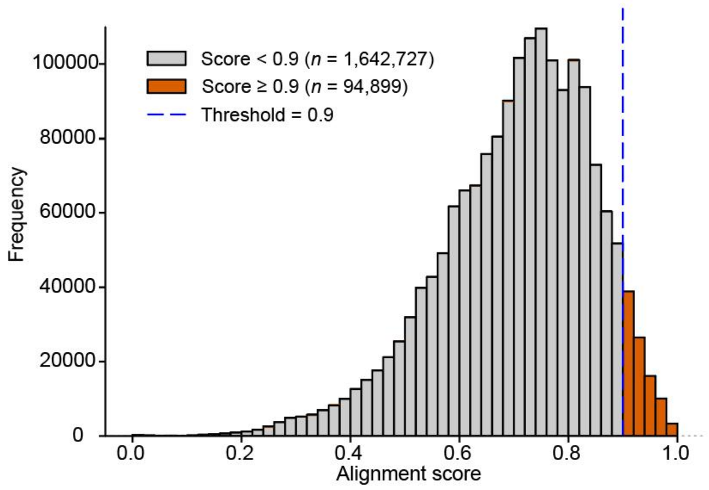

**Supplementary Note Fig. S4 |** Primary filtration of centromere pairs based on HORSCAN alignment scores. The histogram shows the frequency distribution of scores from 1,737,626 pairwise centromere alignments. A score of 0.9 (blue dashed line) is empirically set as the threshold for filtering high-identity candidate pairs.

##### 4.3 Local masking of monomer-level structural variants and periodic variants

Centromeric SVs may artificially inflate the density of adjacent SNPs, due to poor or incorrect alignments of SV-adjacent monomers[6,21]. To mitigate this potential noise, we firstly identified all monomer-level SVs from HORSCAN results, including monomers involved in insertion, deletion or mismatch events. High-density SNPs are strongly co-localized with these monomer-level SVs (**Supplementary Note Fig. S5**). We therefore masked the SVs along with their 10 upstream and 10 downstream flanking monomers. Such masking minimizes the risk that a single monomer-level SV perturbs the alignment of an entire HOR unit, thereby avoiding artificial SNP enrichment around structurally unstable sites.

Additionally, we observed periodic variants in which identical SNPs repeatedly occurred at the same relative positions within a given monomer ID across successive HOR units, even when the compared HOR arrays showed no structural differences. This pattern probably arises from ancient point mutations followed by independent expansions of the two centromeres being compared with identical HOR structural units. Therefore, to eliminate substitution over-estimation caused by this issue, we masked the monomer regions exhibiting periodic variants. Together, direct SV masking and periodic variant masking minimize artifacts from structural variation (e.g., array expansion / contraction and gene conversion), allowing reliable detection of single-base substitutions..

After applying the above masking procedures, we quantified the masked fraction (masked bases divided by total αSat array length) for each centromere pair. Pairs with a masked proportion exceeding 20% were excluded, reducing the candidate set from 94,899 to 978 structurally similar centromere pairs.

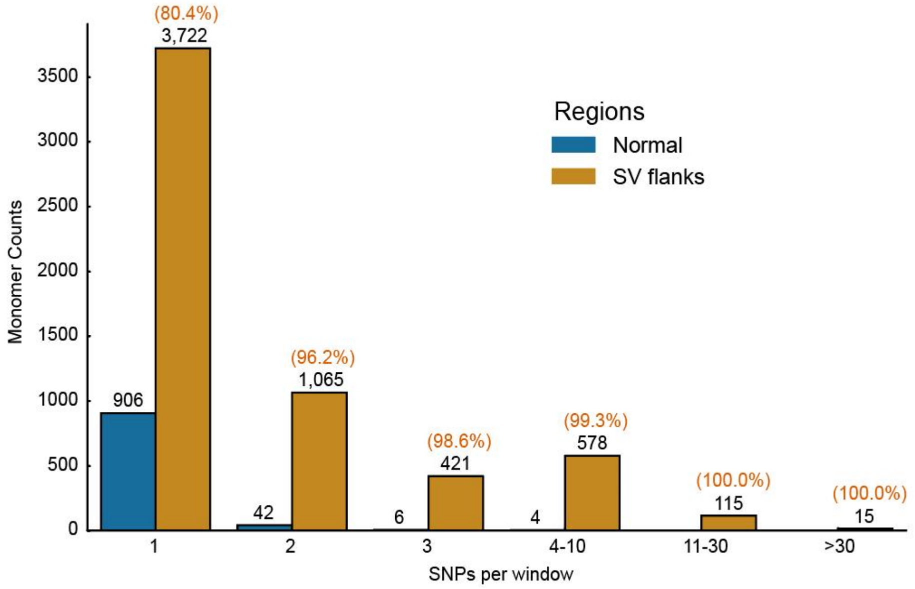


**Supplementary Note Fig. S5 |** High-density SNP clusters strongly co-localize with SVs. This bar chart displays monomer counts grouped by SNP density, comparing monomers in structurally stable ('Normal') regions versus those adjacent to SVs ('Near SV'). A 'Near SV' region is defined as any monomer near an insertion or deletion event, plus its 10 flanking monomers on both sides. The data reveals a stark pattern: while monomers with a single SNP are found in both contexts, virtually all monomers in windows with high SNP density (≥2 SNPs) are located near an SV (96.2%-100.0%). This strong spatial correlation provides the rationale for masking these SV-adjacent buffer zones to filter out SNP clusters that likely represent alignment artifacts rather than true biological variation.

##### 4.4 Manual curation

The final set of 978 centromere pairs underwent rigorous manual curation via visual inspection of HORSCAN alignment plots to identify subtle or complex anomalies that evaded automated detection. Pairs were excluded if they exhibited any of the following distinct features: (a) ≥ 20 monomer InDel events, suggesting potential erroneous alignments between samples; (b) ≥ 40 dispersed, unmasked monomer SNP loci, a frequency exceeding typical background mutation levels and likely caused by shifted annotation of monomer boundaries; or (c) ≥ 3 masked regions of high SNP density (total SNPs >50), indicating fundamental structural inconsistencies between the compared centromeres. Criterion (a) accounted for the majority of exclusions, as numerous InDel events signify a complex rearrangement history that renders such pairs unsuitable for substitution rate analysis. This stringent curation workflow yielded a high-fidelity panel of 93 centromere pairs (mean structural identity ≥ 0.99), comprising 64 family-derived pairs (25 from HG00733 trio, 6 from HG00514 trio, 25 from NA19240 trio, 5 from HG01891 duo, 3 from HG03453 duo) and 29 unrelated pairs (**Supplementary Table 20**).

##### 4.5 High-confidence intervals of centromere and pericentromeres for substitution analysis

Even with high overall structural identity, paralogous sequences or gene conversion events may disrupt monomer-level alignments, confounding SNP calling. To address this, we first computed nucleotide-level edit distances for all aligned monomer pairs (including mismatches), with monomer pairs exhibiting non-zero edit distances designated as containing candidate SNPs. To eliminate false positives arising from paralogy or mis-alignment, each candidate SNP-involved monomer pair underwent phylogenetic validation. Maximum-likelihood phylogenetic trees were constructed using IQ-TREE[17] (v2.1.4-beta) with default parameters, incorporating the monomer pair of interest, their orthologous counterpart, and a same-class monomer pair as a phylogenetic background. A candidate SNP was considered as high-confidence and its position deemed "callable" only if the monomer pair formed a monophyletic clade, providing independent evolutionary evidence of their orthologous relationship and validating the comparison (**Supplementary Note Fig. S6**). Monomer pairs failing this monophyly criterion were masked and excluded from subsequent analyses. This rigorous validation ensures that only evolutionarily orthologous substitutions are included in our rate calculations.

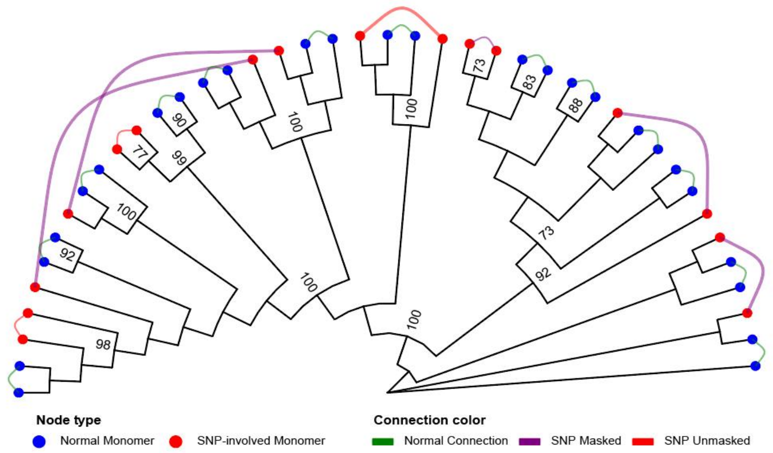


**Supplementary Note Fig. S6** | An example of validating SNPs using phylogenetic congruence between C104-CZA04-01#Pat#chr12 and C107-CZA07-01#Mat#chr12. We filtered candidate SNPs arising from potentially non-orthologous monomer alignments. This tree of neighboring monomers shows individual monomers as tips (blue: Normal; red: SNP) and connects putative orthologs with curved lines. The filtering rule is simple: aligned pairs must also be evolutionary sister taxa. Pairs that violate this, appearing on distant branches of the tree, are considered non-orthologous and are masked (thick purple lines), as they likely represent alignment errors or gene conversion events. Pairs that are sister taxa (thin red lines) demonstrate phylogenetic congruence, and their SNPs are considered true variants.

In pericentromeric regions, we defined callable regions within non-recombination pericentromeric regions, initially specified as the 10 Mbp flanking each core centromere HOR array. We used Winnowmap2[19] (v2.03) to align non-overlapping 50-Kbp windows from the pericentromeric flank to their paired corresponding flank, and performed initial variant calling on these alignment CIGAR values. A two-tiered filtering procedure ensured baseline quality. First, the windows lacking confident, collinear alignment hits in orthologous regions were excluded to eliminate large-scale structural variants. Second, windows exceeding three standard deviations above the chromosomal mean SNP density were masked to remove mutation clusters (**Supplementary Note Fig. S5**) and the boundaries of pericentromeric regions for comparison were refined. This dual filtering effectively removed the regions with both macro-structural variations and micro-scale mutation anomalies, establishing a robust basis for comparative analysis.

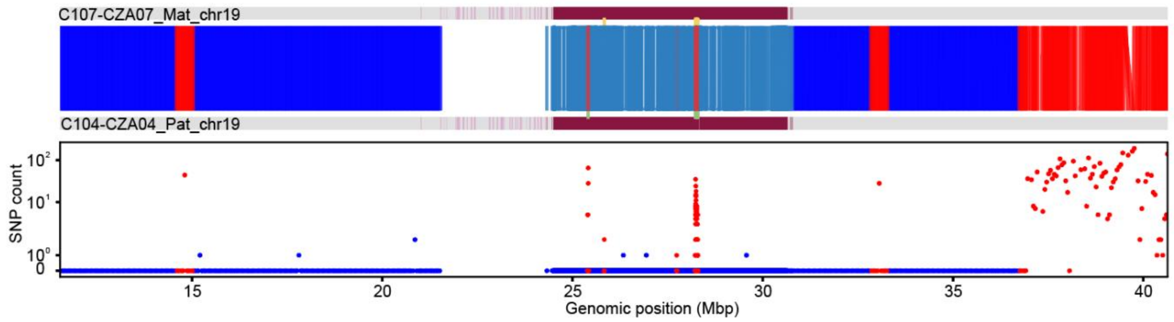


**Supplementary Note Fig. S5 |** A representative example of refining centromeric and pericentromeric regions for substitution analysis. This figure provides a detailed visualization of our filtering process applied to a representative chromosome. The top panel depicts the pairwise alignment of two centromeres (purple) and flanks (gray). The bottom panel shows the corresponding SNP density per window. This example highlights how the workflow effectively removes artifact-prone regions and refine genomic intervals for substitution analysis. Specifically, it removes: (1) a cluster of high-density SNPs (red points) co-localizing with a structurally-masked region (at ~28 Mbp) and (2) a large pericentromeric region characterized by structural variation and anomalously high SNP density (at ~37-40 Mbp).

##### 4.6 Statistical comparison of single-base substitution rates

Substitution rates were calculated as the ratio of high-confidence SNP count ($N_{SNP}$) to callable bases ($L_{callable}$) . For core centromeric regions, $L_{callable}$ was defined as the total length of monomer sequences retained after multi-step masking. For pericentromeric regions, $L_{callable}$ represented the total length of 50-Kbp windows retained after interval filtration and boundary refinement. Core centromere $N_{SNP}$ was derived by summing nucleotide mismatches (edit distance) from HORSCAN alignments of all high-confidence orthologous monomer pairs. Pericentromeric $N_{SNP}$ was similarly obtained by summing mismatches identified via CIGAR strings by Winnowmap2 from all callable windows.

To compare the rates, we calculated the log10-transformed ratio of the centromere substitution rate to the corresponding pericentromeric rate. A log-ratio centered around zero indicates equivalent substitution rates between core centromere and pericentromeric regions. Positive values denote elevated centromeric rate, while negative values indicate reduced rates. To assess the statistical significance, we tested the 93 log-ratios against the null hypothesis of equal rates (log-ratio = 0). As each centromere and its corresponding peri-centromere represent paired data and their rate ratios may not follow a normal distribution, we applied the non-parametric Wilcoxon signed-rank test. A two-sided test was used to evaluate whether centromeric substitution rates significantly differed from pericentromeric rates in either direction.

### Supplementary Note 5 | Reconciliation of centromere variation analyses between this study and Gao *et al.*[22]

Recent independent efforts, including Gao *et al.*[22] have provided complementary perspectives on human centromere diversity through the analysis of large-scale telomere-to-telomere (T2T) genome assemblies. Although both studies leverage high-quality diploid genome resources to investigate previously unresolved aspects of centromeric variation, differences have been reported in HOR diversity, centromere haplotype classification, kinetochore-associated regions, and estimates of centromeric evolutionary dynamics. To facilitate interpretation and enable integration of these complementary resources by the broader community, we systematically examined the analytical frameworks, definitions, and measurements used in the two studies. We further performed targeted comparative analyses using the publicly available HGSVC3 assemblies representing the corresponding samples analyzed by Gao *et al.*, focusing on features for which direct comparisons were supported by shared genomic resources and comparable analytical criteria.

Our analyses demonstrate substantial concordance between the two studies in their major biological conclusions. The observed differences largely arise from distinct analytical frameworks, including annotation strategies, reference baselines, and approaches for estimating evolutionary rates, rather than from fundamental biological discrepancies. Below, we provide a detailed comparison of these aspects.

##### 5.1 Comparison of HOR and structural variant (StV) annotation

The reported differences in HOR structural variant numbers primarily reflect differences in annotation frameworks and discovery baselines rather than biological discordance. Gao *et al.* performed reference-guided αSat annotation using HumAS-HMMER libraries constructed from the T2T-CHM13 reference and previously characterized HOR arrays, with monomer identities assigned according to established HOR models. In contrast, we developed a *de novo* annotation framework that enables the identification of previously uncharacterized HOR architectures through global αSat monomer clustering, HOR reconstruction with HORmon, and HiCAT (**Supplementary Note 2**). This framework enables the detection of non-canonical HOR arrays and StVs that are not represented in existing reference libraries. Detailed comparisons of HOR and StV annotations between this study and the T2T-CHM13 reference models are provided in **Supplementary Tables 13-16**.

The two studies assessed HOR novelty against different reference baselines. Gao *et al.* compared 2,110 centromeric assemblies with the T2T-CHM13 and CHM1 references, identifying 1,870 HOR variants absent from these reference genomes. In contrast, we defined novel HORs relative to the larger HPRCy1 and HGSVC3 pangenome resources, identifying 333 unreported HOR variants within APGp1. Given the substantially different discovery spaces represented by these reference frameworks, the absolute numbers of novel HOR variants are not directly comparable.

Despite differences in annotation frameworks, the two studies reveal highly concordant patterns of population-level centromeric variation. Gao *et al.* reported rare αSat HOR expansions in individuals of African ancestry that altered D7Z1 (CEN07) composition (**Fig. 2e** in their preprint), and we independently observed similar population-specific expansions (**Supplementary Fig. 24**). Likewise, the chromosome 16 αSat variation highlighted by Gao *et al.* (**Supplementary Fig. 18** in their preprint) corresponds to the CenHap-B configuration identified in our analysis (**Fig. 4d**), and their chromosome 21 example (**Supplementary Fig. 22** in their preprint) similarly matches the CenHap-B architecture identified in our study (**Fig. 4e**). Together, these examples demonstrate that independent annotation frameworks converge on the same major patterns of centromeric haplotype diversity.

##### 5.2 Comparison of centromere haplotype (CenHap) classification frameworks

The difference in reported CenHap numbers between the two studies primarily reflects distinct definitions of centromere haplotype identity. In our study, CenHaps are classified based on pairwise synteny distances that integrate αSat monomer-cluster composition and HOR structural variants. In contrast, Gao *et al*. defined CenHaps primarily from phylogenetic relationships inferred from approximately 20-Kbp flanking sequences adjacent to αSat arrays. Using a human-chimpanzee divergence time of 6.2 million years and a 1-million-year divergence threshold for haplotype delineation, they identified 226 CenHaps. Thus, the two frameworks capture complementary aspects of centromere diversity: their approach reflects the evolutionary history of flanking genomic regions, whereas ours resolves structural diversity within αSat arrays.

These two approaches provide complementary perspectives on centromere evolution. Structure-based classification enables direct comparison of HOR organization and is supported by concordance among CenHap assignments, independent k-mer clustering, and flanking-region phylogenies. We nevertheless recognize that the resolution of this approach may be limited in highly homogeneous satellite arrays, given the fixed 96% monomer clustering threshold used for classification (**Methods**). Conversely, phylogenetic classification provides an evolutionary framework for estimating haplotype divergence, although its resolution is constrained by the availability of informative variants within relatively short flanking regions.

##### 5.3 Comparison of centromere dip region (CDR) and kinetochore-associated site annotations

To assess the concordance of kinetochore-associated centromere annotations, we applied our CDR identification framework to 20 HGSVC3 samples and compared the resulting annotations with those reported by Gao *et al*. Because some manually curated HGSVC assemblies analyzed by Gao *et al*. were not publicly available, we performed this comparison using 326 publicly available uncurated chromosome assemblies. The resulting CDR annotations showed strong positional concordance between the two studies: 278 of 326 chromosomes (85.3%) exhibited Jaccard indices greater than 0.7, with reciprocal overlap ratios also exceeding 0.7 (**Supplementary Note Fig. S7**).

The remaining discrepancies primarily reflect differences in CDR filtering and merging criteria rather than divergent identification of kinetochore-associated regions. Our framework retains minor CDRs supported by local sequence features, whereas Gao *et al*. applied additional filtering and manual curation. In addition, the two studies used different rules for merging adjacent CDRs: we generally treated CDRs separated by more than 100 Kbp as independent regions, whereas Gao *et al*. merged some closely spaced CDRs following manual evaluation. For example, we identified a minor CDR in NA19650#hap2#chr20 that was not reported by Gao *et al*., while two CDRs separated by approximately 100 Kbp in NA20509#hap1#chr14 were merged into a single region in their annotation but retained as independent regions in ours (**Supplementary Note Fig. S7**).

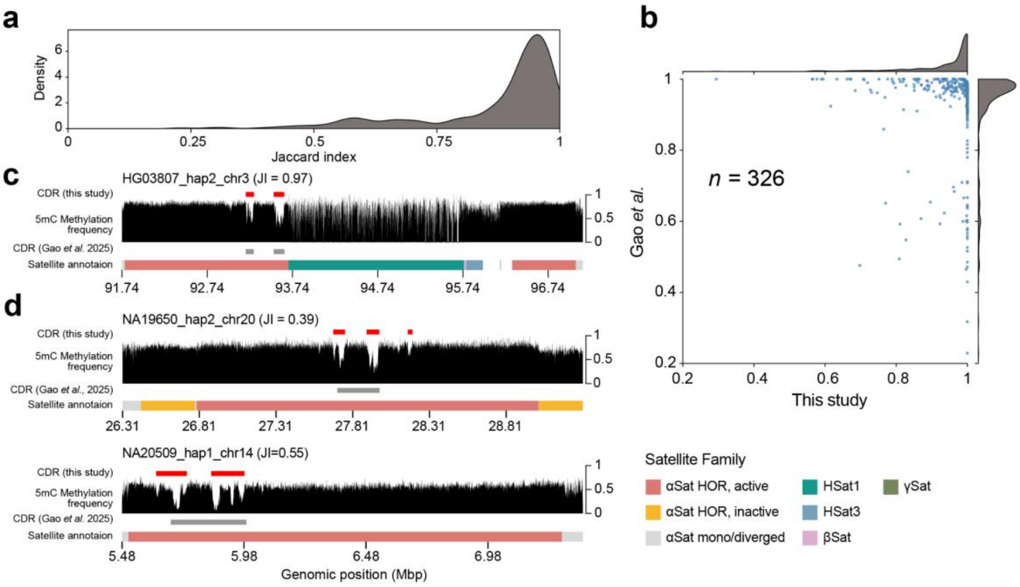

**Supplementary Note Fig. S7 | Comparison of CDR annotation in this study and Gao *et al.* a**, Density plot of Jaccard indices for CDRs annotated by the two workflows. **b**, Overlap proportions between the two pipelines: each point represents the proportion of intersecting CDRs relative to each pipeline's total. **c**, Representative example showing concordant CDR annotations between the two pipelines. **d**, Representative examples showing discordant CDR numbers between our annotations (red) and those from Gao *et al*. (gray).

##### 5.4 Comparison of centromeric mutation-rate estimates

Gao *et al*. reported elevated sequence divergence within centromeres, whereas our analysis detected no significant increase in orthologous single-base substitution rates relative to pericentromeric regions. These measurements capture different aspects of centromeric evolution and therefore are not directly comparable. Gao *et al*. estimated centromeric sequence divergence from whole-array alignments using Kimura's neutral divergence model applied to 10-Kbp windows. This metric reflects the cumulative sequence divergence accumulated over evolutionary time, including nucleotide substitutions and additional processes such as concerted evolution, gene conversion, unequal crossing-over, and copy-number turnover. In contrast, our analysis specifically quantified single-base substitution rates among orthologous centromeric sequences. We established monomer-level orthology through repeat-resolved alignments, excluded structural variants and flanking regions, and retained only monomer groups supported as orthologous by phylogenetic analysis (**Supplementary Note 4**). This strategy provides a conservative estimate of point substitution accumulation while reducing potential effects from paralogous alignments and structural turnover. Together, these measurements capture distinct dimensions of centromere evolution: sequence turnover and homogenization processes shaping satellite array composition, versus the accumulation of fixed single-base substitutions within orthologous repeat units. Both represent biologically meaningful aspects of centromeric evolution but reflect different evolutionary signals. Similar orthology-aware strategies have recently been applied to plant centromeres[23], where repetitive sequence architecture presents analogous challenges for estimating mutation rates.

We could not independently reproduce the pedigree-based *de novo* mutation-rate estimates reported by Gao *et al*., because the underlying curated CEPH-1463 assemblies were not publicly available. Nevertheless, their reported summary statistics highlight an important consideration when interpreting absolute mutation rates in satellite arrays. Several *de novo* events were concentrated within highly localized intervals, including seven variants within a single 177-bp monomer on chromosome 14 and six variants within a 393-bp interval on chromosome 5. Such extreme clustering within highly homogeneous repetitive sequences can be challenging to distinguish from residual paralogous alignment errors or localized gene-conversion events, complicating the interpretation of local mutation-rate estimates.

Collectively, our comparison with Gao *et al.* demonstrates that the two datasets provide complementary perspectives on human centromere diversity. The observed differences in HOR variants, CenHap classifications, CDR annotations, and mutation-rate estimates primarily reflect differences in analytical definitions, reference frameworks, and evolutionary quantities measured. Importantly, both studies independently reveal extensive human centromere diversity, including population-specific HOR expansions, chromosome-specific centromere architectures, and complex evolutionary dynamics within αSat arrays.

### Supplementary Note Reference
